# ALGAL HOMOLOGS OF THE PLANT CER1 AND CER3 PROTEINS ARE FUNCTIONAL HYDROCARBON–FORMING ENZYMES

**DOI:** 10.1101/2025.10.28.685152

**Authors:** Ángel Baca-Porcel, Bertrand Légeret, Mallaury Cabanel, Mathilde Le-Cossec, Damien Sorigué, Yonghua Li-Beisson, Florian Veillet, Fred Beisson

## Abstract

In plants, very-long-chain (VLC) alkanes (C25-C35) are secreted onto the epidermal surface of aerial organs, often forming the primary component of the waterproof cuticular wax layer. This secretion plays a vital role in preventing desiccation in terrestrial environments. The key to VLC alkane biosynthesis in plants lies in the complex formed by two homologous membrane-bound proteins, ECERIFERUM 3 (CER3) and ECERIFERUM 1 (CER1). This complex transforms an acyl-CoA substrate into an aldehyde intermediate, which is then converted into an alkane. The ability to synthesize and secrete alkanes is believed to have been a pivotal event in the evolution of land plants from their green algal ancestors. Interestingly, a single homolog of CER1 and CER3, known as CER1/3, has been identified in certain algae. However, the functionality of this protein remains to be investigated. In this study, we present the functional characterization in yeast of CER1/3 proteins from various algal species belonging to the green lineage. We demonstrate that CER1/3 proteins alone can efficiently mediate hydrocarbon biosynthesis. Furthermore, we demonstrate that point mutations in conserved motifs in the N- or C-terminal domains of CER1/3 lead to impaired hydrocarbon biosynthesis. Additionally, we show that coexpressing plant CER3 with algal CER1/3 results in longer alkanes being formed in yeast. Together, these findings support the hypothesis that the alkane-forming CER1/CER3 complex found in land plants evolved from a green algal CER1/3 bifunctional enzyme through a process of gene duplication followed by protein specialization.

## Introduction

The colonization of terrestrial habitats by land plants has had a profound impact on the evolution of organisms and ecosystems on Earth (Bowman, 2022). This event greatly affected biogeochemical cycles, facilitating the terrestrialisation of various other lineages. Land plants probably evolved from a single clade of streptophyte algae around 470 million years ago (Delwiche and Cooper, 2015; de Vries and Archibald, 2018). The transition from aquatic to terrestrial environments was driven by key cellular and physiological adaptations to the new environmental constraints associated with terrestrial habitats, such as desiccation, ultraviolet radiation, and pathogens. Symbiotic interactions with fungal partners are also believed to have been crucial for the earliest land plants (Delaux and Schornack, 2021).

Limiting losses of internal water was necessarily a key adaptation to terrestrial environments. This was achieved notably through building an extracellular lipid-based structure, the cuticle, onto the surface of the organisms (Delwiche and Cooper, 2015). In land plants, cuticle components are synthesised in the epidermal cells of aerial organs and subsequently exported onto their surface to form a waterproof layer (Yeats and Rose, 2013). The cuticle is made of a cutin scaffold filled and covered with cuticular waxes. Cutin is a polyester that consists mainly of hydroxy-, epoxy-, and dicarboxy C16- and C18-fatty acids, together with glycerol (Pollard et al., 2008; Beisson et al., 2012). However, it is the waxes rather than the cutin itself that confer impermeability to the cuticle. The cuticular waxes of many plants consist mainly of very-long-chain (VLC, > C20) aliphatic compounds (Samuels et al., 2008; Bernard and Joubès, 2013; Lee and Suh, 2015). The C16 and C18 acyl-CoAs are substrates for the fatty acid elongase (FAE) complex, which produces VLC acyl-CoAs up to C30 and beyond (Haslam and Kunst, 2013). Following elongation, VLC acyl-CoAs are converted into cuticular waxes via two major pathways, the alcohol-forming pathway, yielding primary alcohols and wax esters, and the alkane-forming pathway, which produces mainly alkanes, aldehydes, secondary alcohols, and ketones. These two pathways account for approximately 20% and 80%, respectively, of the total stem wax load in *Arabidopsis* (Jenks et al., 1995; Li-Beisson et al., 2013). All these wax components are then exported to the outer cell wall of epidermal cells by ATP-binding cassette (ABC) transporters of the ABCG subfamily, enabling subsequent assembly into intracuticular waxes associated with the cutin polymer and epicuticular waxes present on top of the cutin layer as a smooth film and/or as crystals (Lee and Suh, 2015). In many plant species, alkanes are the major component of cuticular waxes (Bourdenx et al., 2011; Bernard and Joubès, 2013; Lee and Suh, 2015). Alkanes are synthesized in the endoplasmic reticulum (ER) from VLC acyl-CoAs by the ECERIFERUM proteins CER1 (Aarts et al., 1995) and CER3 (Rowland et al., 2007), which form a membrane complex with the cytochrome B5 (Bernard et al., 2012; Pascal et al., 2019).

The CER1 and CER3 proteins of land plants are homologous and possess a bi-domain structure consisting of an N-terminal fatty acid hydroxylase (FAH) domain, resembling the catalytic domain of the yeast sphingolipid α-hydroxylase Scs7, and a C-terminal WAX2 domain, which resembles the cleft region of *Synechococcus elongatus* acyl-ACP reductase (AAR), an aldehyde-producing enzyme (Wang et al., 2019; Chaudhary et al., 2021; Kojima et al., 2024). The N-terminal domain (NTD) of plant CER1 is highly conserved and possesses the histidine-rich motifs necessary for the formation of alkanes by the CER1/CER3 complex (Bernard et al., 2012; Kojima et al., 2024). In contrast, the C-terminus of CER1 is less conserved. Conversely, the C-terminal domain (CTD) of CER3 is highly conserved and contains a potentially catalytic cysteine and a putative NADPH binding site, which may be involved in NADPH utilization. The NTD of plant CER3 is less conserved, and the histidine-rich motifs present are not essential for the activity of the CER1/CER3 complex (Bernard et al., 2012; Kojima et al., 2024). It has long been hypothesized that CER3 is a reductase that mediates the reduction of C_n_ VLC acyl-CoAs into C_n_ aldehyde intermediates, and that CER1 is an aldehyde decarbonylase that produces the alkanes (Jenks et al., 1995; Bernard et al., 2012; Pascal et al., 2019).

A recent landmark study in *Arabidopsis* provides indirect evidence of an aldehyde-forming activity of CER3. This study shows that CER3 also interacts with SOH1, a reductase that converts aldehydes into primary alcohols (Li et al., 2025). The CER1-CER3-SOH1 complex was proposed to participate in the regulation of transpiration through the cuticle by modulating the alkane/primary alcohol ratio of cuticular waxes in response to environmental conditions. Given the importance of alkanes in cuticular waxes, the appearance of the CER1 and CER3 proteins in land plants was likely to be a key to cuticle evolution and terrestrialization. Interestingly, a few algal species, such as the marine chlorophyte *Ostreococcus tauri* (*O. tauri*) (Sorigué et al., 2016), possess a CER1/CER3 homolog, but always as a single-copy gene referred to as CER1/3. Phylogenetic analyses suggest that this CER1/3 algal homolog underwent duplication in ancestral embryophytes to produce the paralogs CER1 and CER3 (Wang et al., 2019; Chaudhary et al., 2021). Given the aquatic habitat of green microalgae, the idea that CER1/3 plays a role in producing extracellular alkanes seemed doubtful, and the function of CER1/3 remained unexplored.

Here, we demonstrate that the CER1/3 protein found in various green algal species is a bifunctional enzyme that can efficiently synthesize hydrocarbons (HCs) by producing aldehydes and transforming them into HCs. These results support the view that an ancestral algal bifunctional CER1/3 protein has evolved into a plant CER1/CER3 protein complex in which each original function is performed by a specialized homolog.

## Materials and Methods

### Bioinformatic analyses

To identify algal homologs to plant CER1/CER3 proteins, BLASTP searches were performed across various databases using the *A. thaliana* CER1 as query protein sequence. The sequences of the various algal CER1/3 and plant CER1 and CER3 proteins used in this study are available in **Table S1**. AlphaFold 3 software (Abramson et al., 2024) was employed for protein 3D structure prediction and modeling. The generated protein 3D structures were visualized using PyMOL software (Delano Scientific LLC). For the construction of a maximum-likelihood phylogenetic tree, protein sequences were first aligned with MAFFT (https://www.ebi.ac.uk/jdispatcher/msa/mafft) using default parameters. The alignments were then trimmed in SeaView 5 using Gblocks with the three options for a less stringent selection. Finally, the tree was constructed in Seaview 5 using PhyML (LG matrix and 1000 bootstrap replicates).

### Algal strains and culture conditions

The *O. tauri RCC4221* strain was obtained from the Roscoff Culture Collection. This strain was cultured in sterile artificial seawater supplemented with F/2 medium (Sigma-Aldrich). Cultures were maintained in T-25 flasks with vented filter caps (CytoOne) for about two weeks at 20-22°C under an illumination of about 50 µmol photons.m^−2^.s^−1^ with a 12-hour light/12-hour dark photoperiod. Cell counts were routinely performed using a Multisizer 3 (Coulter). The *Klebsormidium nitens* (*K. nitens*) *NIES-2285* strain was obtained from the microbial culture collection of the National Institute of Environmental Studies, Japan. This strain was cultured in plates containing C medium (Kodama and Fujishima, 2015) with 2% (w/v) agar, incubated at 22°C under about 50 µmol photons·m^−2^.s^−1^ with a 12-hour light/12-hour dark photoperiod.

### Plasmid construction and yeast expression

Protein sequences from various algal CER1/3 and land plant CER1/CER3 (**Table S1**) were back-translated using yeast codon optimization (Geneious software) and synthesized (TwistBioscience) for cloning into the Golden Gate MoClo system. Site-specific mutations in the coding sequences of the different CER1, CER3, and CER1/3 proteins were generated using specific primers. Each coding sequence was assembled in a transcriptional unit using the yeast copper-inducible promoter (*pCUP1*) and an appropriate terminator sequence (i.e. *tPGK1, tTDH1* and *tADH1*) (Lee et al., 2015). Final plasmids were constructed by assembling at least one transcriptional unit mediating the expression of various algal CER1/3 and land plant CER1 and CER3 to the high-copy 2micron yeast origin of replication and the *URA3* auxotrophic selection marker (Lee et al., 2015). Final plasmids were introduced into the *S. cerevisiae* INV*Sc*1 (ThermoFisher Scientific) diploid strain (*MATa his3D1 leu2 trp1-289 ura3-52 MAT his3D1 leu2 trp1-289 ura3-52*) using the classic LiOAc/PEG3350 method. Yeast cells growing on plates filled with selective dropout media lacking uracil (DOM-URA) were further checked by colony PCR. All the plasmids and strains constructed and used in this work are detailed in **Table S2** and **Table S3**, respectively. For protein expression, cultures (15 mL) were initiated at an optical density (OD600) of 0.2 in DOM-URA media in 50-mL mini bioreactor centrifuge tubes. Cells were grown in an orbital shaker at 30°C, 180 rpm. Protein expression was induced with 0.033 mg.mL^-1^ CuSO_4_ 4 h after starting the culture. The cultures were grown in these conditions for 48 h. The OD600 of the different cultures was then measured before samples were collected for GC-MS/FID analysis to scrutinize fatty acid and HC content within yeast cells.

### Analysis of intracellular HCs and fatty acids by transmethylation and saponification

For the identification of HCs in *O. tauri* cells, 14 mL of cell culture was harvested after 8 days of culture (≈1.3 x 10^8^ cells.ml^-1^) and centrifuged at 3,200 g for 15 min to pellet the cells. For the identification of HCs in *K. nitens* cells, a scoop of *K. nitens NIES-2285* culture (≈4 week-old culture) was collected. Cell pellets were then transmethylated with 2 mL of methanol containing 5% (v/v) sulfuric acid by heating in sealed glass tubes at 85°C for 90 min. After cooling, 3 mL of 0.9% (w/v) NaCl and 200 µL of *n*-hexane were added. The samples were shaken for 5 min and centrifuged at 3,200 g for 5 min to facilitate phase separation and the recovery of HCs and fatty acid methyl esters (FAMEs) in the organic phase. The *n*-hexane phase was then transferred to another glass tube. The *n*-hexane organic solvent was evaporated under a gentle stream of nitrogen gas. Subsequently, the fatty acids and HCs were resuspended in 900 µL of water and 100 µL of NaOH (10 M). The glass tubes containing the samples were sealed, shaken for 5 min, and heated for 90 min at 85°C (for saponification). After cooling, 500 µL of 0.9% (w/v) *n*-hexane was added to the samples. The samples were shaken for 5 minutes and centrifuged at 3,200 g for 5 min to facilitate phase separation and the recovery of the unsaponifiable compounds (such as HCs) in the organic phase. Finally, the *n*-hexane phase was analyzed by GC-MS/FID. For the identification and quantification of HCs and fatty acids in yeast cells, 8 mL of cell culture was centrifuged at 3,200 g for 15 min to pellet the cells. Internal standards (10 µg of *n-*docosane and 10 µg of triheptadecanoyl glycerol) were added for quantification, and cell pellets were transmethylated with 2.5 ml of methanol containing 5% (v/v) sulfuric acid. Samples were homogenized and heated for 90 min at 85 °C in sealed glass tubes. After cooling down, 3 ml of 0.9% (w/v) NaCl and 200 µL of *n-*hexane were added. Samples were then shaken for 5 min and centrifuged at 3,200 g for 5 min to allow phase separation and recovery of FAMEs and HCs in the organic phase (*n-*hexane). Finally, the *n-* hexane phase was analyzed by GC-MS/FID. For the synthesis of the C21:6 alkene (all-*cis*-3,6,9,12,15,18-heneicosahexaene) derived from all-cis-4,7,10,13,16,19-docosahexaenoic acid (DHA), 100 µL of a 2.5 mg.mL^-1^ DHA solution in ethanol (250 µg) and 200 µL of a 10 mg/mL purified recombinant FAP protein (Sorigué et al. 2017) from *Chlorella variabilis* NC64A (2 mg) in a buffer 150 mM NaCl, 10 mM Tris-HCl pH 8.0, and 20% (w/v) glycerol was added to a 10-mL sealed vial containing 1 mL of Tris-HCl buffer (pH 8.0). The vial was then incubated for 1 h under 300 µmol photons.m^-2^.s^-1^ of blue light to trigger the decarboxylation of DHA into C21:6 alkene by FAP. To extract the alkene, 900 µL of the aqueous reaction mixture was subjected to saponification (see above). After saponification, the alkene was extracted by adding 500 µL of *n*-hexane, followed by 5 minutes of shaking and centrifugation at 3,200 *g* for 5 minutes to facilitate phase separation and recovery of the alkene in the organic phase. Finally, the *n*-hexane phase was analyzed by GC-MS/FID to check purity.

### GC-MS/FID analyses

For analysis of transmethylated and saponified *O. tauri* and *K. nitens* samples, as well as transmethylated yeast samples, a GC-FID 7890B Agilent coupled to a 5977B series mass detector was used. For GC, a HP-5MS column (length 30 m, internal diameter 0.25 mm, and film thickness 0.25 mm) was used with helium (1.4 mL.min^-1^) as the carrier gas. One µL of the *n-*hexane phase was injected in splitless mode. The GC parameters were as follows: oven initial temperature, 50°C for 1 min; ramp, 10°C.min^-1^ to 150°C and then 5°C.min^-1^ to 310°C; hold for 8 minutes. The MS was run in full scan over 40-500 Th (electron impact ionization at 70 eV). Peaks were identified based on their retention time and mass spectrum and quantified based on the FID signal using C17 fatty acid methyl ester and C22 alkane internal standards.

## Results

### Green algal species that possess a single CER1/3 protein lack a FAP homolog

Protein sequence similarity searches in various databases using the *Arabidopsis* CER1 sequence allowed to identify single CER1/CER3 homologs (hereafter called CER1/3) in 10 out of 79 genomes of green algae investigated (**Table 1, Table S4 and Table S5)**. Additionally, a CER1/3 homolog was also identified in the genomes of the cryptophyta algae *Proteomonas sulcata* and *Cryptophyceae* sp., both of which result from a secondary endosymbiosis (**Table S5**). Among the 12 CER1/3 protein sequences, 7 of them (from *Ostreococcus tauri RCC4221, Micromonas commoda RCC299, Prasinoderma coloniale CCMP1413, Klebsormidium nitens NIES-2285, Mesotaenium kramstae* Lemmermann *NIES-657, Proteomonas sulcata CCMP1175*, and *Cryptophyceae* sp. *CCMP2293*), along with the CER1/3 protein from *Tetraselmis convolutae* (Xavier Bailly, personal communication), were studied in greater detail in the following paragraphs. Green algal CER1/3 proteins were present in the three phyla of the green lineage (Li et al., 2020): chlorophyte algae (genera: *Ostreococcus, Micromonas, Tetraselmis*), streptophyte algae (*Klebsormidium, Mesotaenium*), and Prasinodermophytes (*Prasinoderma*), which indicated that the ancestral CER1/3 probably existed before the divergence of these three groups. A phylogenetic analysis, which includes these CER1/3 protein homologs, supports the view that the algal CER1/3 gene was duplicated into CER1 and CER3 in the common ancestor of land plants (**Figure 1**), as suggested previously (Chaudhary et al., 2021).

**Table 1:**
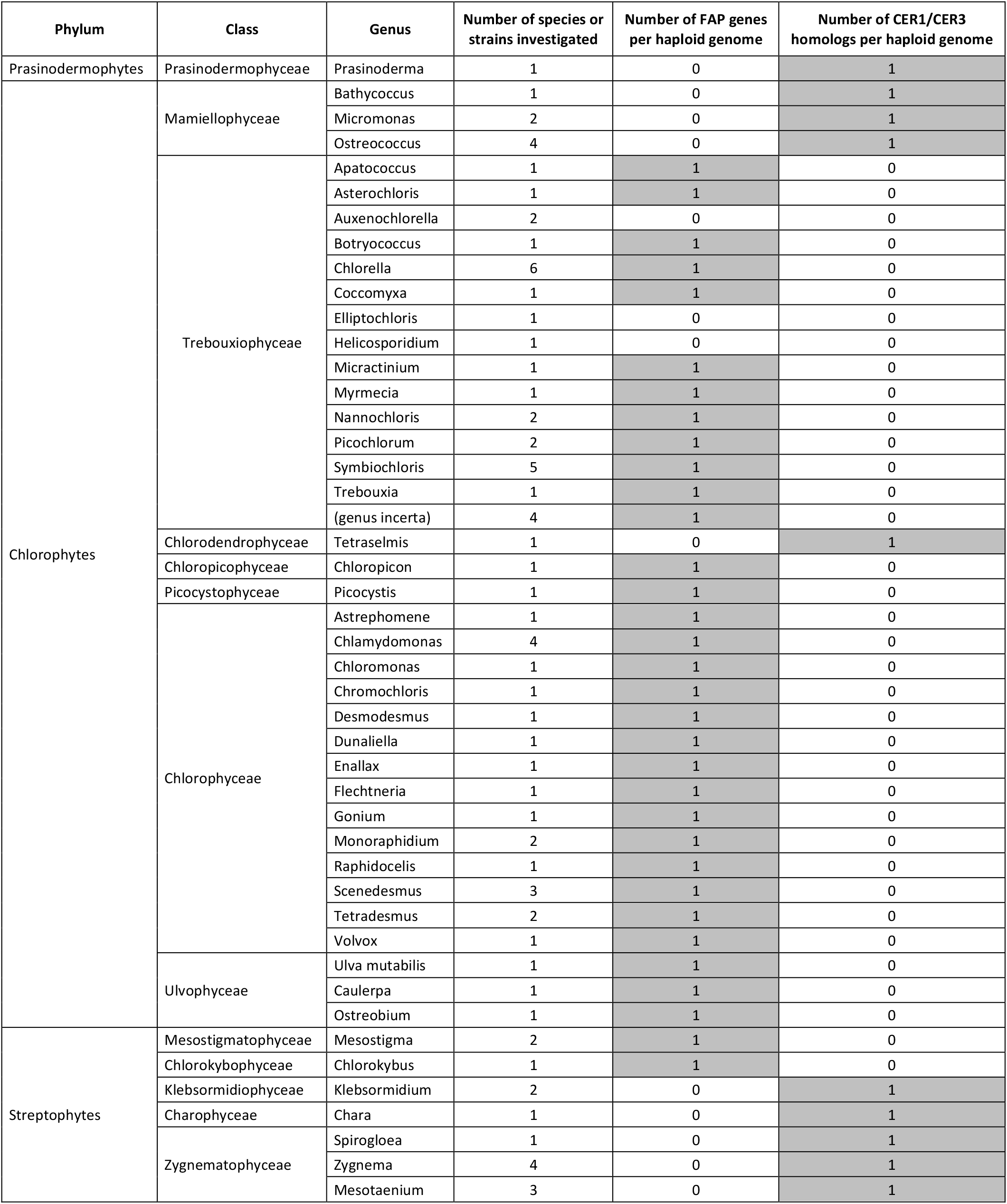
Presence of FAP and CER1/CER3 homologs in green algae genomes. This table shows the genera and the number of species or strains whose genomes were examined for the presence of FAP and CER1/CER3 homologs. The full names of the species or strains used are listed in **Table S4**. The genomes were obtained from the PhycoCosm database (https://phycocosm.jgi.doe.gov/algae/algae.info.html).

**Figure 1:**
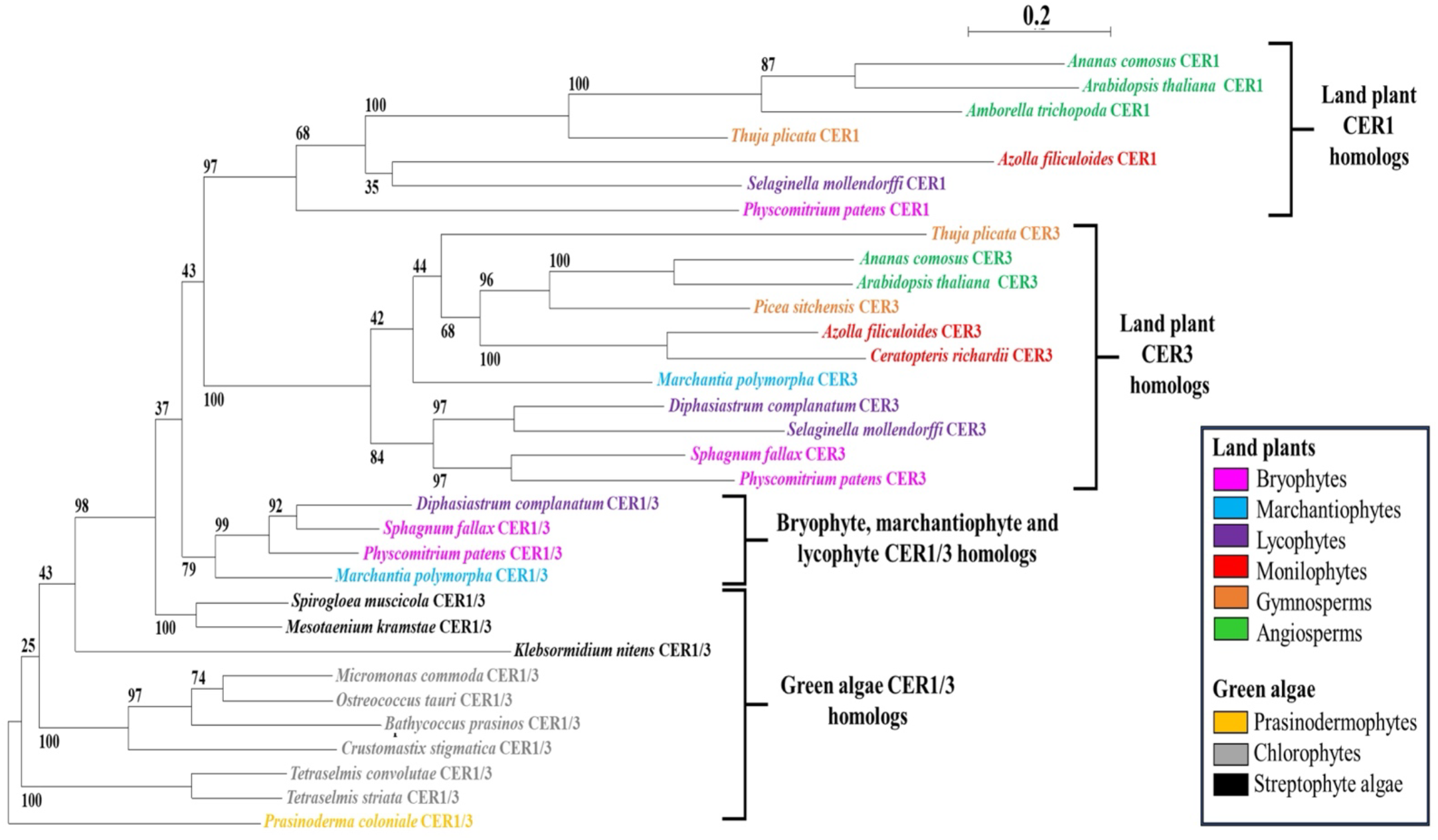
Phylogenetic tree of CER1, CER3 and CER1/3 homologs in land plants and green algae. The tree was constructed using the maximum likelihood method based on CER1/CER3 homologs from various plants and algae. Homologs were classified as CER1, CER3 or CER1/3 prior to tree building based on conserved motifs (see **Figure 3**). The tree was rooted with the *Prasinoderma coloniale* CER1/3 homolog. Bootstrap support values from 1000 replicates are indicated. The scale bar represents 0.2 substitutions per site.

Interestingly, the presence of CER1/3 proteins in these green algal species correlates with the lack of the FAP photoenzyme (**Table 1**), which converts long-chain (LC) free fatty acids into intracellular alka(e)nes (Sorigué et al., 2017; Moulin et al., 2021). We then assessed whether the green algae *O. tauri* and *K. nitens* were able to synthesize HCs. In *O. tauri* cells, we identified in the unsaponifiable fraction the C21:6 alkene previously reported (Sorigué et al., 2016). We further showed that it was very likely to be all-*cis*-3,6,9,12,15,18-heneicosahexaene, which derives from the C22:6 VLC polyunsaturated fatty acid DHA **(Figure 2)**. In the unsaponifiable fraction of *K. nitens* cells, we detected a C17 alkane **(Figure 2)**, a common HC found in various algae. These results strongly suggest that CER1/3 proteins may indeed be involved in HC synthesis.

**Figure 2:**
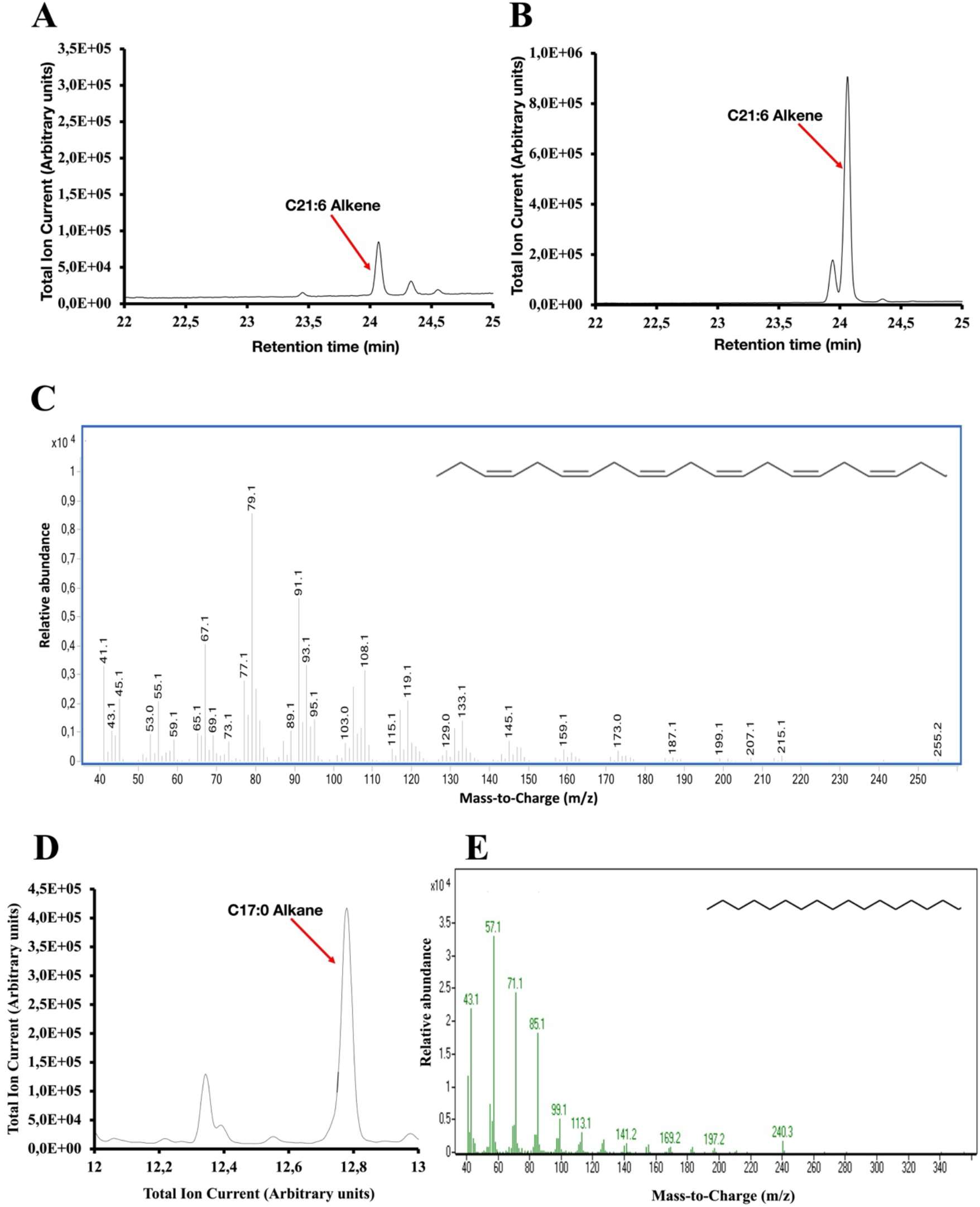
Synthesis of HCs by *O. tauri RCC4221* and *K. nitens NIES-2285* algal species. **A)** GC-MS chromatogram of the unsaponifiable cell material from *O. tauri RCC4221* strains cultured for 8 days at 20°C under 50 µmol photons·m^−2^·s^−1^ with a 12-hour light /12-hour dark photoperiod. The HC is indicated by a red arrow. The C21:6 alkene (all-*cis*-3,6,9,12,15,18-heneicosahexaene) was the only HC detected, with a retention time of 24.1 minutes. **B)** GC-MS chromatogram obtained after injecting a C21:6 alkene (*n*-heneicosahexaene) standard, prepared by the decarboxylation of a DHA (all-*cis*-4,7,10,13,16,19-docosahexaenoic acid) standard using the photoenzyme FAP. **C)** Mass spectrum of the C21:6 alkene peak show in panel A. **D)** GC-MS chromatogram of the unsaponifiable cell material from *K. nitens NIES-2285* strains cultured during 4 weeks at 22°C under 50 µmol photons·m^−2^·s^−1^ with a 12-hour light /12-hour dark photoperiod. The HC is indicated with a red arrow. The C17:0 alkane (*n-*heptadecane) was the only HC detected, with a retention time of 12.8 minutes. **E)** Mass spectrum of the C17:0 alkane peak show in panel D.

### The algal CER1/3 homolog has the key motifs of both land plant CER1 and CER3

Aligning different green algal CER1/3 protein sequences revealed that their NTD and CTD display a high degree of similarity to the key motifs of plant CER1 and CER3, respectively **(Figure S1, S2 and S3)**. Detailed inspection of the multiple alignments was conducted in the region of i) the putative NADPH-binding site and the putative catalytic cysteine found in the CTD of land plant CER3s but absent in land plant CER1s (Kojima et al., 2024), and ii) in the region of the three catalytic histidine-rich motifs (Bernard et al., 2012) highly conserved in the NTD of land plant CER1s but less conserved in land plant CER3s. This analysis showed that all eight of the algal CER1/3 homologs studied possess both the three highly conserved histidine-rich motifs in their NTDs and the conserved putative NADPH-binding site, as well as the putative catalytic cysteine in their CTDs **(Figure 3)**.

**Figure 3:**
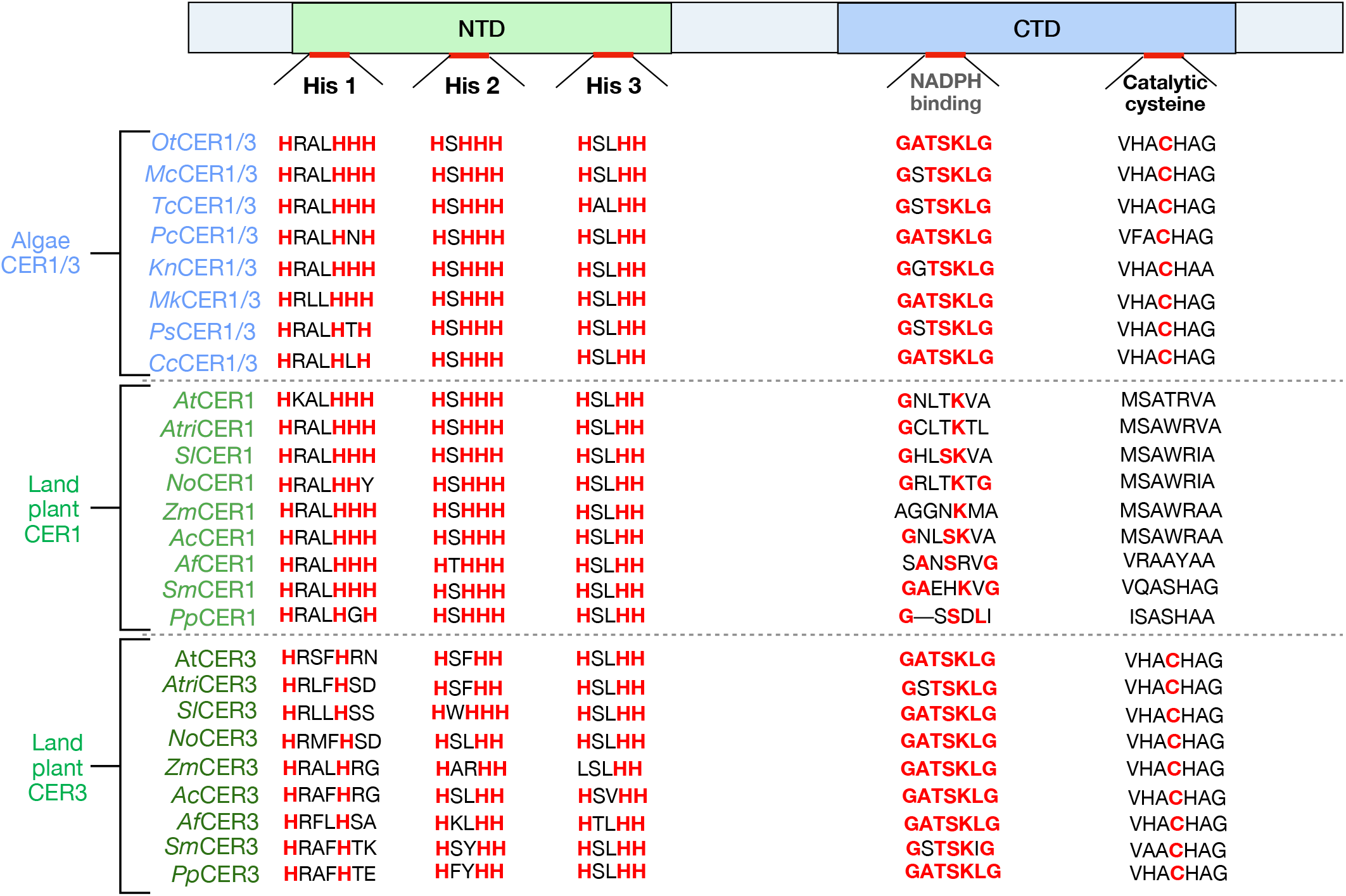
Conservation of putative catalytic motifs in the two domains of land plant CER1 and CER3 and algal CER1/3. The histidines of the histidine-rich motifs, the putative catalytic cysteine as well as the conserved residues of the NADPH binding site are indicated in red bold. *Ac*: *Ananas comosus, Af*: *Azolla filiculoides, At*: *Arabidopsis thaliana, Atri*: *Amborella trichopoda, Cc: Cryptophyceae sp. CCMP2293, Kn*: *Klebsormidium nitens, Mc: Micromonas commoda, Mk: Maesotaenium kramstrae, No*: *Nymphaea odorata, Ot: Ostreococcus tauri, Pc: Prasinoderma coloniale, Pp*: *Physcomitrium patens, Ps: Proteomonas sulcata, Sl*: *Solanum lycopersicum, Sm*: *Selaginella moellendorfii, Tc: Tetraselmis convolutae, Zm*: *Zea mays*.

The 3D structure of the *O. tauri* CER1/3 protein predicted by Alphafold 3 was then superposed onto the predicted 3D structures of *Arabidopsis* CER1 and CER3 **(Figure 4 and Figure S4)**. This analysis showed that the predicted structure of OtCER1/3 is highly similar to those of AtCER1 and AtCER3. The seven other algal CER1/3 proteins have a predicted structure similar to that of OtCER1/3 **(Figure S5)**. The *Arabidopsis* CER1 and CER3 proteins are embedded in the endoplasmic reticulum membrane through the presence of transmembrane domains. Sequence analysis of the OtCER1/3 protein revealed a predicted subcellular localization to the ER and the presence of transmembrane domains (DeepLoc2.1), suggesting that green algal CER1/3 may localize to the same subcellular compartment as their land plant counterparts. Taken together, these results suggest that the green algal CER1/3 homolog may possess the structural features allowing to produce some HCs alone.

**Figure 4:**
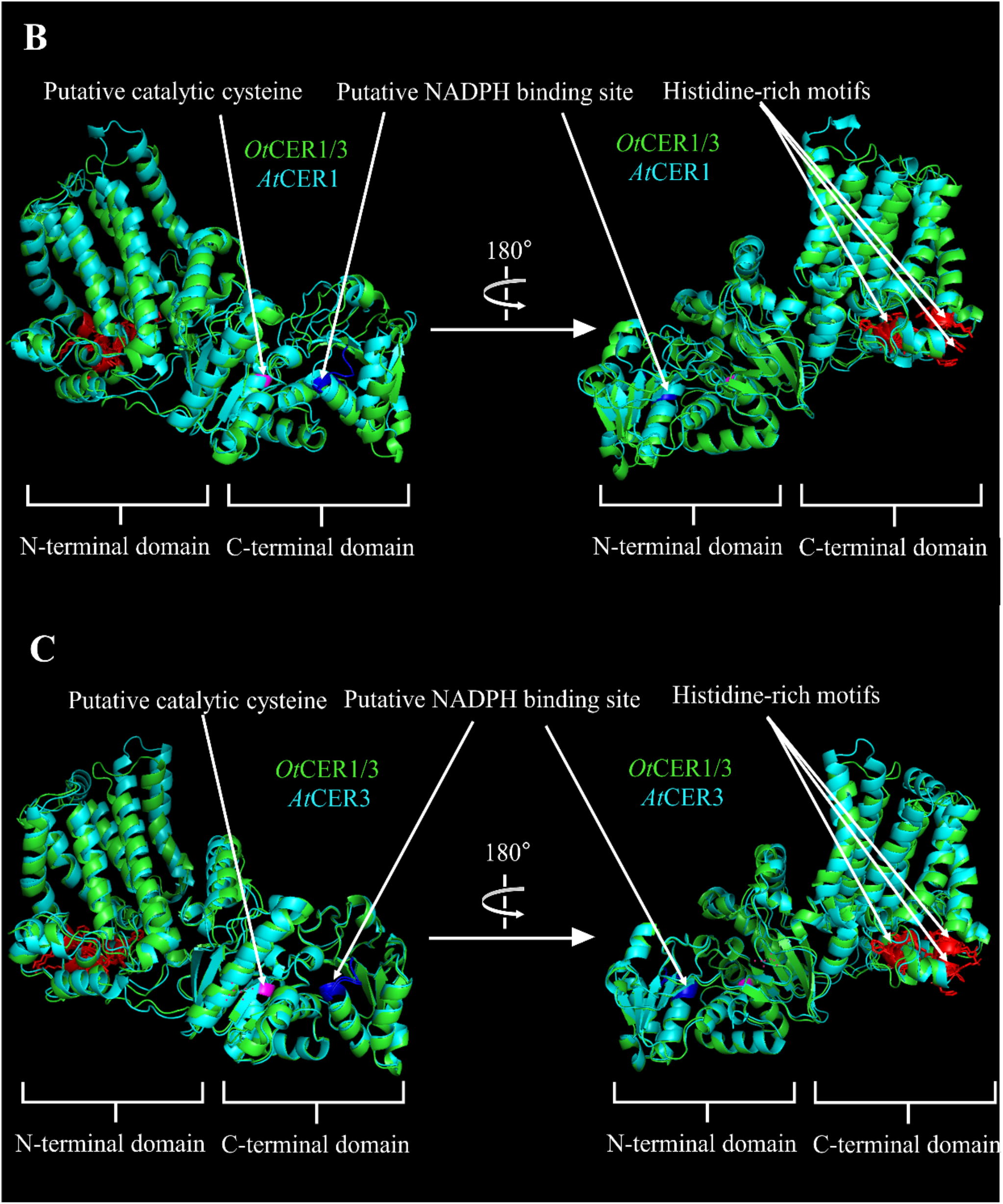
Superposition of *O. tauri* CER1/3 (*Ot*CER1/3) predicted structure with *A. thaliana* CER1 (*At*CER1) (upper panel) and *A. thaliana* CER3 (*At*CER3) (lower panel). Histidine rich motifs are shown in red, while the putative NADPH binding site and the putative catalytic cysteine are shown in dark blue and pink, respectively.

### Algal CER1/3 proteins are functional HC-forming enzymes

To determine whether algal CER1/3 proteins are HC-forming enzymes or have any CER1-like or CER3-like activity, genes encoding several CER1/3 were cloned and expressed in the yeast *S. cerevisiae*. A CER1/3 protein from a species representing each of the three phyla of green algae was selected, namely *Prasinoderma coloniale* (*P. coloniale*) for Prasinophytes, *O. tauri* for Chlorophytes, and *K. nitens* for Streptophytes. *S. cerevisiae* does not naturally produce fatty acid-derived HCs but has VLC fatty acids and acyl-CoAs up to 26 carbons with traces of fatty acids with 28 carbons (Denic and Weissman, 2007; Bernard et al., 2012; Wang et al., 2016). Strikingly, expression in yeast of each algal CER1/3 alone resulted in the production of HCs (**Figure 5**). OtCER1/3 produced three different LC HCs, which were identified based on retention time and mass spectrum as 7-pentadecene (**Figure S6**), 8-heptadecene (**Figure S7**), and *n*-heptadecane (**Figure S8**). Expression of *K. nitens* CER1/3 (KnCER1/3) produced *n*-heptadecane, while *P. coloniale* CER1/3 (PcCER1/3) produced 8-heptadecene and *n*-heptadecane **(Figure 5)**. Interestingly, no VLC HCs (i.e. >C20) were detected in any of the strains expressing the different algal CER1/3 homologs **(Figure 5)**.

**Figure 5:**
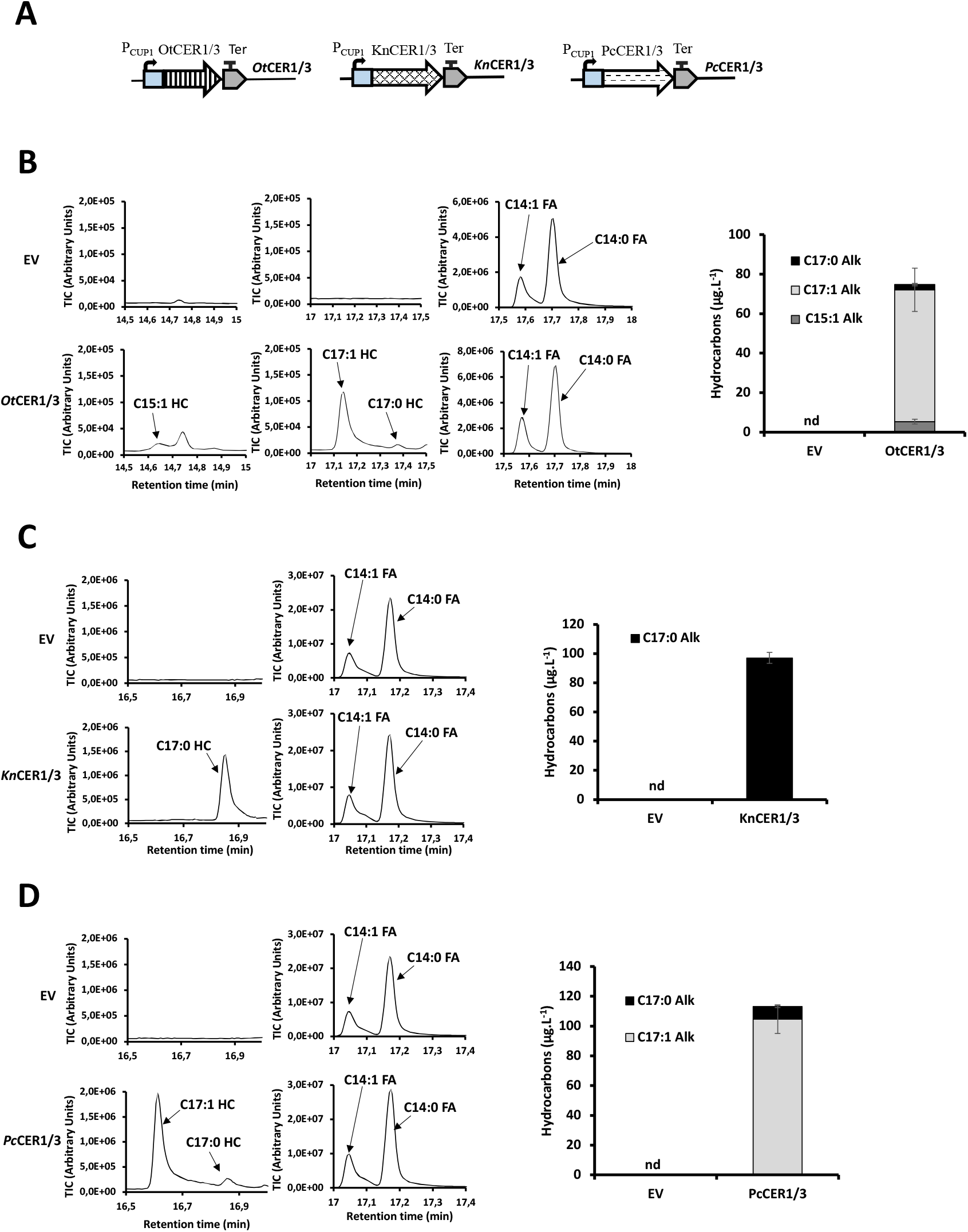
Expression of *O. tauri, K. nitens and P. coloniale* CER1/3 proteins in yeast. **A)** Scheme of the genetic constructs used for the expression of the *O. tauri* (*Ot*CER1/3), *K. nitens* (*Kn*CER1/3) *and P. coloniale* CER1/3 (*Pc*CER1/3) proteins. **B)** <u>Left panel:</u> Chromatograms showing the different HCs, and the C14 fatty acids (used as loading control), present in yeast strains expressing *Ot*CER1/3 protein homolog. <u>Right panel</u>: Quantification of the HCs produced by yeast strains expressing *Ot*CER1/3 homolog. **C)** <u>Left panel:</u> Chromatograms showing the different HCs, and the C14:1 and C14:0 fatty acids (used as loading control), present in yeast strains co-expressing *Kn*CER1/3. <u>Right panel</u>: Quantification of the HCs produced by yeast strains expressing *Kn*CER1/3. **D)** <u>Left panel:</u> Chromatograms showing the different HCs, and the C14 fatty acids (used as loading control), present in yeast strains expressing *Pc*CER1/3 protein homolog. <u>Right panel</u>: Quantification of the HCs produced by yeast strains expressing *P. coloniale* CER1/3 homolog. Error bars represent the standard error based on six biological replicates. ‘nd’ = non detected, ‘HC’ = hydrocarbon, ‘FA’ = fatty acid.

To extend these results, expression of the CER1/3 proteins from another streptophyte alga (*M. kramstae*) and a cryptophyte alga (*P. sulcata*) were also performed. Expression of each CER1/3 protein led to the production of HCs, although at lower levels compared to the first three tested (**Figure S9**). Specifically, *M. kramstae* CER1/3 produced 8-heptadecene, while *P. sulcata* CER1/3 produced both 8-heptadecene and *n*-heptadecane (**Figure S9**).

In order to determine whether the key motifs of the land plant CER1 and CER3 proteins are also important for the catalytic activity in algal CER1/3 proteins, we produced different versions of the OtCER1/3 protein with specific amino acid changes. Single amino acid change in any of the His-rich motifs (H142A, H155A, H243A) abolishes HC production (**Figure S10**). Similarly, amino acid changes in the putative NADPH-binding (A455N and T456L) or catalytic (C581A) sites also impair the production of HCs **(Figure S11)**.

Taken together, these results demonstrate that OtCER1/3 is a functional HC-forming enzyme that has the conserved catalytic motifs of both land plant CER1 and CER3, suggesting a potential bifunctional activity.

### Algal CER1/3 proteins use the same intermediate as the plant CER1/CER3 complex

Since algal CER1/3 proteins expressed alone in yeast can produce HCs, it is possible that these proteins are bifunctional (i.e., possessing both CER3-like activity, converting acyl-CoAs into aldehydes, and CER1-like activity, reducing aldehydes to alkanes). Alternatively, algal CER1/3 proteins may operate through a different mechanism and may convert acyl-CoAs directly to alkanes or through an intermediate different from aldehydes. To evaluate whether an aldehyde is a possible substrate of the different algal CER1/3 protein homologs, coexpression in yeast of OtCER1/3 together with *A. thaliana* CER3 (or *A. thaliana* CER1 as a control) was attempted. AtCER3 has been suggested to produce C24-C32 aldehydes, while AtCER1 is thought to form C27-C31 alkanes from C28-C32 aldehydes (Bernard et al., 2012; Pascal et al., 2019; Kojima et al., 2024). Strikingly, compared to OtCER1/3 expression alone, co-expression of both OtCER1/3 and AtCER3 not only led to an increase in LC HCs (7-pentadecene, 8-heptadecene, and heptadecane), but also resulted in the production of the C25:0 alkane *n*-pentacosane **(Figure 6** and **Figure S12**). By contrast, co-expression of OtCER1/3 with AtCER1 did not yield VLC alkanes and even reduced LC alkanes, producing only 8-heptadecene and small amounts of *n-* heptadecane **(Figure 6)**. Expressing each of the four other CER1/3 proteins from various green algae together with AtCER3 also resulted in the formation of a C25:0 alkane (**Figure S13-S15)**. Additionally, co-expression of PsCER1/3 together with AtCER3 also resulted in the production of a C23:0 alkane (**Figure S14** and **Figure S15**).

**Figure 6:**
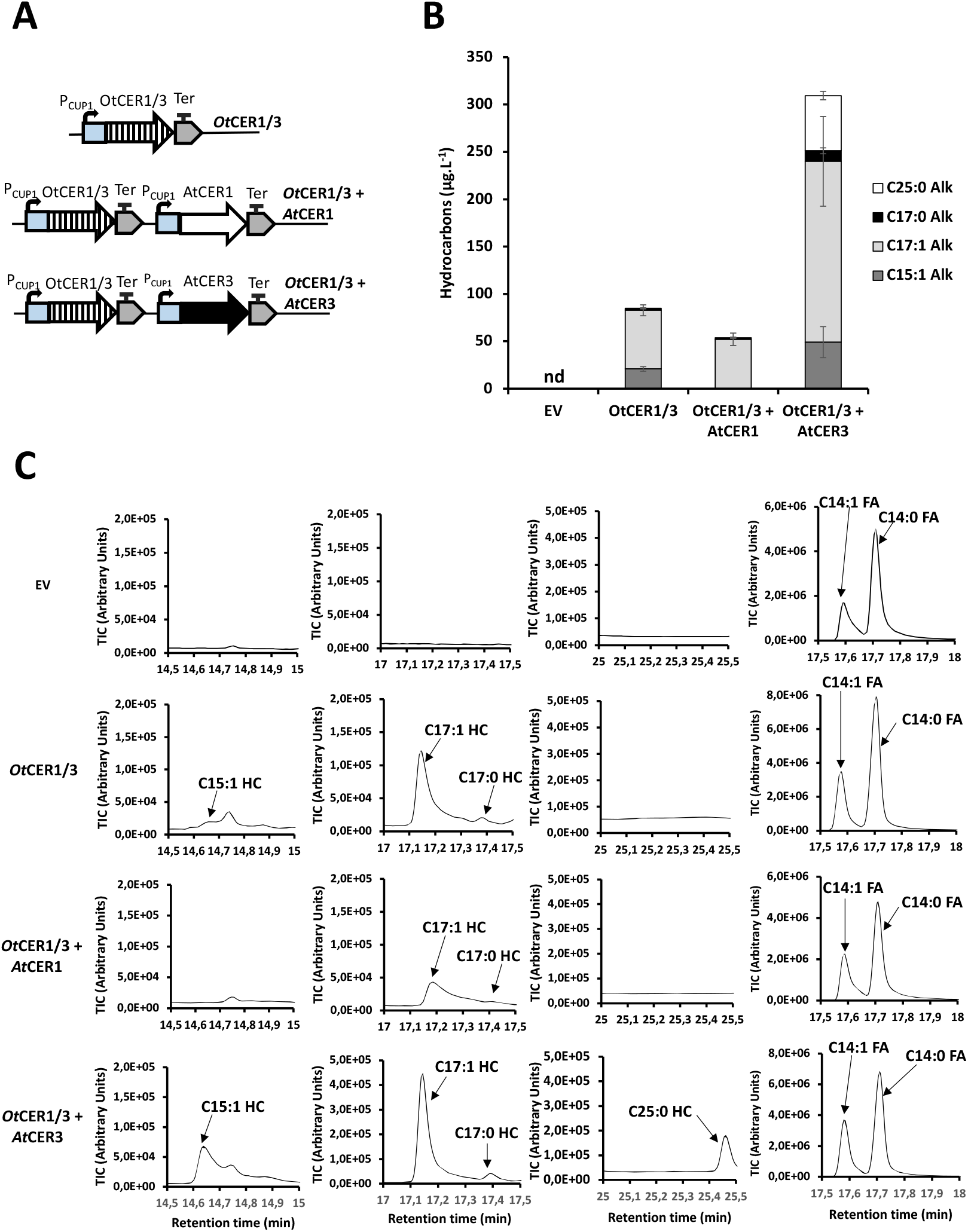
Expression of OtCER1/3 and *A. thaliana* CER1 or CER3 in yeast. **A)** Scheme of the genetic constructs. OtCER1/3 and AtCER1 or AtCER3 were co-expressed in yeast strains under the control of a copper inducible promoter (P_CUP1_). **B)** HCs produced by the different yeast strains. **C)** Chromatograms showing the different HCs, and the C14:1 and C14:0 fatty acids (used as loading control), present in the different yeast strains. Error bars represent the standard error based on six biological replicates. ‘nd’ = non detected, ‘HC’ = hydrocarbon, ‘FA’ = fatty acid.

Taken together, these data therefore demonstrate that, like AtCER1, the algal CER1/3 proteins have the ability to use the intermediate produced by AtCER3 (presumably a C26:0 aldehyde), to synthesize a C25:0 alkane. The CER1/3 proteins thus perform the same catalytic reactions as the plant CER1/CER3 complex.

To determine which domain(s) of AtCER3 are responsible for C25:0 alkane formation when co-expressed with OtCER1/3, AtCER3 variants mutated in the histidine-rich motifs (i.e. H147A, H161A or H250A), the putative catalytic cysteine (i.e. C587A), or the putative NADPH binding site (i.e. A461N and T462L) were co-expressed with OtCER1/3. The production of the C25:0 alkane was unaffected by mutations in the different Histidine-rich motifs of AtCER3 (**Figure S16**). On the other hand, mutations in the putative NADPH binding site or the putative catalytic cysteine of AtCER3 resulted in the abolition of the C25:0 alkane production and a considerable decrease in the alkane content compared to the control construct (**Figure S17**). These results are consistent with those reported in a previous study, where co-expression of AtCER1 with AtCER3 mutated in the various histidine-rich motifs did not abolish the production of VLC HCs in yeast, suggesting that the histidine-rich motifs are not essential for CER3 activity (Bernard et al., 2012). On the other hand, our results suggest that the CTD motifs of CER3 are essential for CER3 activity. In order to support this hypothesis, we co-expressed AtCER1 together with AtCER3 mutated or not in the different motifs of its CTD domain. While the co-expression of AtCER1 with non-mutated AtCER3 resulted in the production of C17:0 and C25:0 alkanes, co-expression with AtCER3 mutated in either the putative NADPH binding site or the putative catalytic cysteine led to the abolition of alkane production (**Figure S18**).

## Discussion

In most algal lineages, the presence of fatty acid-derived HCs is attributed to the presence of the photoenzyme FAP, which converts free fatty acids into HCs in presence of light (350-520 nm) (Sorigué et al., 2016, 2017; Moulin et al., 2021). However, in some green algae, the *FAP* gene is absent from the genome, while homologs of plant CER1 and CER3 proteins, designated as CER1/3, are present. Here we establish that the presence of FAP or CER1/3 genes in the 79 green algal genomes investigated is mutually exclusive. CER1/3 homologs were also identified in some cryptophyte algae, specifically in *P. sulcata* and *Cryptophyceae* sp. *CCMP2293*. Additionally, algal species lacking the FAP enzyme but possessing a CER1/3 homolog, such as *O. tauri* and *K. nitens*, were able to produce HCs, suggesting that these algal CER1/3 proteins function as HC-producing enzymes. After performing multiple sequence alignments and analyzing the AlphaFold-predicted 3D structures of various algal CER1/3 proteins, as well as *Arabidopsis* CER1 and CER3 proteins, we observed that algal CER1/3 proteins possess in their NTD the key motifs of plant CER1 (histidine-rich motifs), and in their CTD, the key motifs of plant CER3 (a putative NADPH-binding site and a putative catalytic cysteine). This suggests not only that algal CER1/3 proteins may be HC-producing enzymes, but also that the enzyme may be bifunctional, performing both CER1 and CER3 activities.

Using heterologous expression in yeast, we further demonstrate that green algal CER1/3s are functional HC-forming enzymes. Our experiments, along with the fact that there is only a single CER1/3 homolog in the algal genomes, reinforce the hypothesis that CER1/3 is a bifunctional protein with both CER1-like and CER3-like activities. Co-expression of various algal CER1/3 proteins with *A. thaliana* CER3, a putative aldehyde-producing enzyme, not only increased HC production compared to expression of the algal CER1/3 proteins alone (except for *P. coloniale* CER1/3), but also altered the HC product profile. In addition to LC HCs, VLC HCs (mainly C25:0 alkane) were also produced in all tested strains. Furthermore, when *O. tauri* CER1/3 was co-expressed together with *A. thaliana* CER3 individually mutated in the putative NADPH binding site and the putative catalytic cysteine, only LC HCs were produced and in lesser amounts than those produced by yeast strains expressing *O. tauri* CER1/3 alone. However, when *O. tauri* CER1/3 was co-expressed together with *A. thaliana* CER3 mutated in the histidine-rich motifs, the strain was capable of producing LC and VLC HCs. These results support the hypothesis that *A. thaliana* CER3 provides a substrate, likely an aldehyde, that algal CER1/3 proteins can convert into HCs. As shown in this study and in previous reports (Bernard et al., 2012), *A. thaliana* CER3, in collaboration with *A. thaliana* CER1, is involved in the synthesis of both VLC and LC HCs when expressed in yeast. Therefore, *A. thaliana* CER3 may not only increase the aldehyde pool available to algal CER1/3 homologs but may also supply VLC substrates—likely aldehydes—for VLC HC production. In contrast, co-expression of *O. tauri* CER1/3 with *A. thaliana* CER1 did not result in increased HC production or in the production of VLC HCs. This suggests that *A. thaliana* CER1 may not provide a suitable substrate for *O. tauri* CER1/3, possibly because *A. thaliana* CER1 functions as an aldehyde decarbonylase rather than an aldehyde reductase like *A. thaliana* CER3.

As demonstrated in this study and in previous work (Bernard et al., 2012), the key motifs of the CTD of land plant CER3 (the putative NADPH-binding site and putative catalytic cysteine) and the key motifs of the NTD of land plant CER1 (the histidine-rich motifs) are essential for HC production, as deletion of these motifs impairs HC synthesis in yeast. In addition, this study investigated the functional importance of these key motifs in the NTD and CTD of *O. tauri* CER1/3. Individual mutations in the histidine-rich motifs in the NTD, as well as in the putative NADPH-binding site and putative catalytic cysteine in the CTD, resulted in impaired HC production. These findings also support the hypothesis that algal CER1/3 proteins are bifunctional enzymes. In algal CER1/3, both CTD and NTD motifs are essential for HC production, whereas in land plants, only the NTD motifs are essential in CER1 and only the CTD motifs are essential in CER3. These findings also support the hypothesis that the land plant *CER1* and *CER3* genes originated from the duplication and differentiation of an ancestral algal *CER1/3* gene. As proposed in another work (Kojima et al., 2024), it seems that after gene duplication, CER1 and CER3 carried out different processes of differentiation. For CER1, the NTD may remained enzymatically active while the CTD may lose its function. On the other hand, for CER3, the CTD may remained enzymatically active while the NTD may lose its function. This specialization of land plant CER1 and CER3 might have helped land plants to colonize more easily terrestrial environments by yielding a more efficient alkane-forming complex, and also possibly by providing a specialized protein producing only aldehydes, which may interact with another alcohol-forming enzyme (Li et al., 2025).

Based on all the *in silico* and *in vivo* experiments, as well as previous works, we propose a model (**Figure 7**) in which the CTD of algal CER1/3 protein homologs may function as an aldehyde reductase, catalyzing the conversion of LC acyl-CoA into aldehydes. The CTD contains a putative NADPH binding site, which may utilize the reducing power of NADPH to supply the electrons necessary for this conversion. Additionally, the CTD harbors a putative catalytic cysteine residue, which may be part of the active site and directly involved in the enzymatic conversion of LC acyl-CoA into LC aldehydes. It is possible, as proposed for AAR enzyme, that the acyl-CoA first forms a thioester bond with the putative catalytic cysteine, and then NADPH provides a hydride to break this bond, releasing the acyl chain as an aldehyde (Gao et al., 2020). On the other hand, the NTD of algal CER1/3 protein homologs may function as an aldehyde decarbonylase, catalyzing the conversion of the aldehydes into LC HCs. This NTD contains conserved histidine-rich motifs that may be involved in the coordination of metal atoms (that may form a dimetal center) at the active site of the NTD, similar to what has been described in the yeast sphingolipid α-hydroxylase (Zhu et al., 2015). These metal atoms (to be determined) might be part of the active site of the NTD. Further experiments will be required to validate this model and elucidate the *in vivo* function of the alkanes produced by CER1/3 proteins.

**Figure 7:**
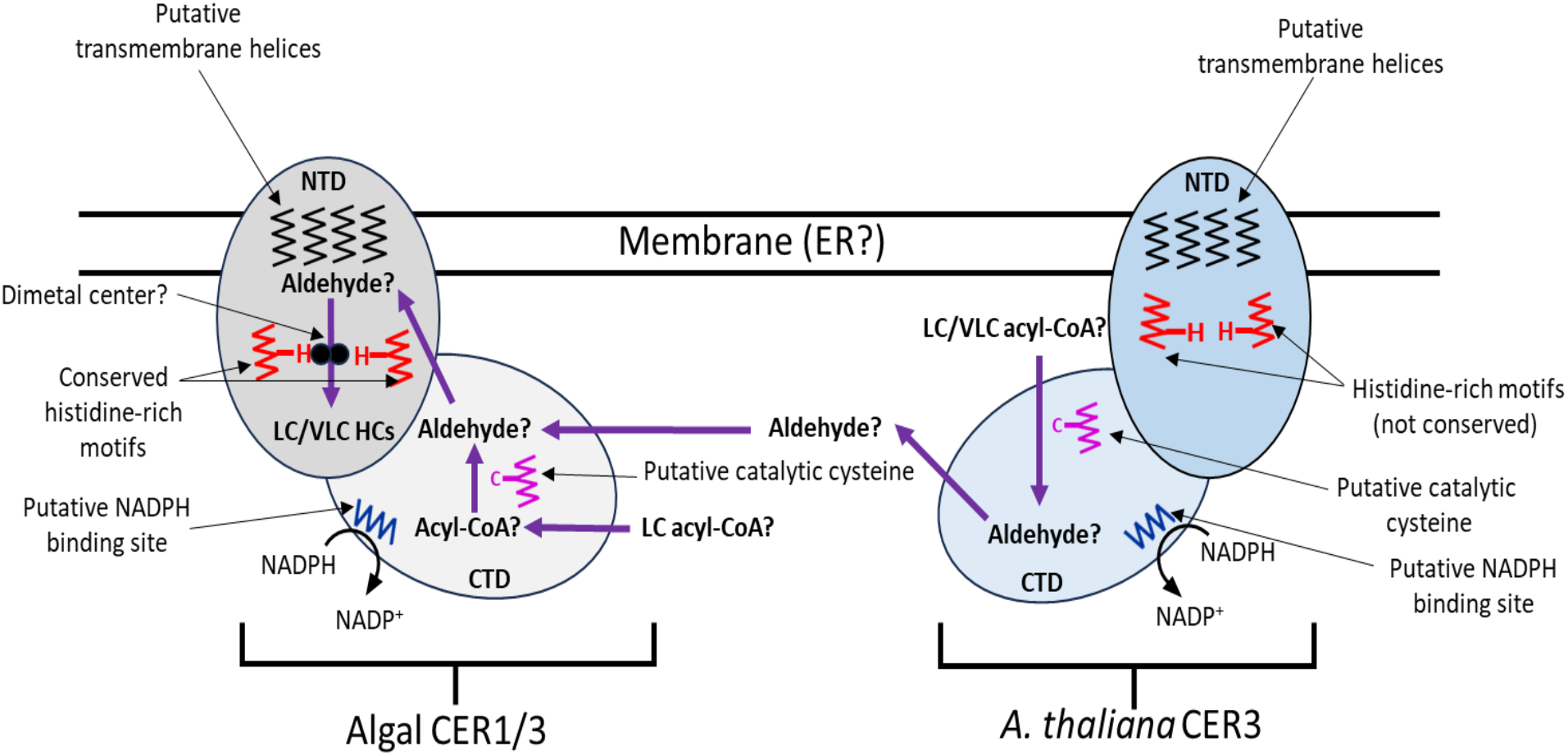
Proposed biochemical mechanism of algal CER1/3 proteins and *A. thaliana* CER3. This scheme represents a biochemical model illustrating the proposed reaction mechanism of algal CER1/3 proteins in the production of LC HCs, and highlighting the synergistic interaction between algal CER1/3 protein homologs and *A. thaliana* CER1/3 proteins in the production of both LC and VLC HCs in a heterologous system.

## Conclusion

In conclusion, the present work identified new algal HC-producing enzymes, CER1/3 in the phyla *Chlorophyta, Prasinophyta*, and *Streptophyta*—as well as in certain algae belonging to the *Cryptophyta* subkingdom. These CER1/3 algae are homologs of the land plant HC-producing enzymes CER1 and CER3. These algal enzymes are found in single copy in algal genomes lacking the FAP enzyme which is the other HC-producing enzyme identified in algae. This CER1/3 enzyme is likely bifunctional since it can produce HCs alone when expressed in yeast, compared to land plant CER1 and CER3 which need to be together for the HC production. Moreover, algal CER1/3 enzymes mainly produce LC HCs while land plant CER1 and CER3 produce mainly VLC HCs. This work also demonstrated that in a heterologous system, algal CER1/3 can function together with *A. thaliana* CER3 to produce VLC HCs, suggesting that algal CER1/3 and land plant CER1/CER3 act through the same aldehyde intermediate. Finally, our results indicate that land plant CER1 and CER3 might be originated from the duplication and differentiation of an ancestral algal CER1/3 enzyme. Further studies are needed to determine the role that duplication of CER1/3 and the subsequent specialisation into plant CER1 and CER3 may have played in helping land plants to colonise terrestrial environments.

## Credit authorship contribution statement

All authors discussed the results and contributed to the final manuscript. ABP, MLC, MC, BL, PAT, PG. and FV. carried out the experiments. ABP, FV, and FB, contributed to the design and implementation of the research. ABP, FV, BL and FB contributed to the analysis of the results. ABP wrote the original draft. Finally, ABP, FV, FB, DS and YLB contributed to the editing and revision of the final manuscript.

## Acknowledgements

We thank Dr. Laurence Blanchard for helpful discussions as well as Louise Ardouin, Dr. Thierry Desnos and Dr. Zhongze Li for technical and IT support. We acknowledge Dr. Xavier Bailly for providing the *T. convolutae* CER1/3 sequence. This work was funded in part by the Agence Nationale de la Recherche project Photoalkane (ANR-18-CE43-0008) and Brown Alkaenes (ANR-24-CE20-3101-01). The funding provided by the European Union Regional Developing Fund (ERDF), the Région Sud, the French Ministry of Research, and the CEA to the HelioBiotec platform is also acknowledged. A.B.P. was supported by a PhD fellowship “Economie Circulaire du Carbone” from CEA.

