## Supplemental data for "ALGAL HOMOLOGS OF THE PLANT CER1 AND CER3 PROTEINS ARE FUNCTIONAL HYDROCARBON–FORMING ENZYMES"

### Supplementary figures and tables

```
KnCER1/3    ---------------------------------MATKPGVLSEWPWQRMGNYKYVLYAPY  27
OtCER1/3    ---------------------------------MATRPGVWYDFPWANVGALKYAVFAPF  27
McCER1/3    ---------------------------------MALRPGPAYKFPWEDMGSFKYLLFVPF  27
PsCER1/3    MWSGIARAGAFAQWFNDSSK--------KQKVRGALKPGILYDSPWESLGSYKYSLFLPF  52
CcCER1/3    MWPGAARLKAMGSWMGGTVTAPNEKEKDSLRHKGATQPGLWYDFPWHAMGDWKYLLYLPF  60
TcCER1/3    ---------------------------------MATKPGPLYEWPWAAAGDLKYLLFTPF  27
PcCER1/3    ---------------------------------MASKPGFMYEFPWEPMGNFKYLLLAPF  27
MkCER1/3    ---------------------------------MATTPGAFYEWPWQSLGNFKYAIYAPF  27
                                              *  **   . **   *  ** :  *:

KnCER1/3    VAAAAYEWGYQGKANEFIAYNAHLGVIAALRYVVMQIWTSIARFPWLVGKYQISSKGIDF  87
OtCER1/3    VLA----VALGK--DDADSFCWHLLAIAAARYVNAQLWISLSRVHAWTRNTRIQAKGIDF  81
McCER1/3    VAT----AALGL--DDADNWAYHMLVIAAIRYVHAQFWISLSRIHAVTQHTKIQAKGIDY  81
PsCER1/3    LYA----VVSGN--DDEDNWCYHMLMIAALRYIQAWIWNFLSRNHHVSGKNRIQAKGVDF  106
CcCER1/3    VVV----LALGL--DDADRWCFHMMAVCVLRFTQAWVWNLLSRNHFISGKTRIQGKGVEF  114
TcCER1/3    IAA----VALGK--DDEDNWASHMLAIAALRYVQNYLWIFVSRCDWLSGRNKIQHKPVTF  81
PcCER1/3    AAC----AALGL--DDADNWCWHMVTLVAVRYALAQLFITASRLHAISGKHRIQKRGIDF  81
MkCER1/3    VAKAVHANLLGG--EDPDNWCLHMVLSSALRMFHGQLWMSASRVHYWTQKHKIQLKGIKF  85
                          ::   :  *:    . *     .:   :*      . :*. : : :

KnCER1/3    KQVDREDSWDDFILFNSLGLTLLGLVLP---------GWQTLPLVNGHGLVVMLLLHASV  138
OtCER1/3    KQVDREDHWDDYILLQTLVIAAVHWMPGL--------GFKDFPLYSGKSFAQLALLHAGP  133
McCER1/3    KQVDREDHWDDYIILQAIIMTLVHKMPYL--------GYNNFPQYNAMGMWQLLLLHAGP  133
PsCER1/3    VQVDREDNWDDYILLQIYVASAVHLLPFLYCWDGECYAYKNFPLHNNKGLIKLLLIHMGP  166
CcCER1/3    KQIDRESNWDDYIILQLYVATAVHCLPFLYCSNGVCHAYRNFPLYDGKGLVKMLLIHAGP  174
TcCER1/3    KQVDRENHWDDYIILQAFVMTAVHWVLP---------GFASFPVNNWNGLWQTLMLHVGP  132
PcCER1/3    EQMDREDNWDDFLILQALVMTAVHWILP---------GFKNFALYDGTGLAQCALLHAGP  132
MkCER1/3    EQVDREERWDDYIILHVLVATFVHAVLP---------GFQNFPLTDGWGLLILLLLHMGP  136
             *:***. ***:::::    : :  :           .:  :   .  .:    ::* .

KnCER1/3    AEWLYYWFHRALHHHWLYTRYHSHHHQSFVTEPITGSVHPFLEHVGYIAIFALPGVLTYV  198
OtCER1/3    TEFIYYWLHRALHHHKLYSAYHSHHHASFVTEPITGSVHPFMEHLMYTANFAIPLLGTWA  193
McCER1/3    TEFIYYWLHRALHHHTLYSWYHSHHHASFVTEPITGSVHPFMEHIMYTANFAIPLVGTWA  193
PsCER1/3    TEFIYYWLHRALHTHQLYANYHSHHHASFVPEVVTGSVHPFMEHLMYTANFAIPLVGTWL  226
CcCER1/3    TEFIYYWLHRALHLHSLYARYHSHHHASFVPEAVTGSVHPFMEHLMYTINFAIPLLGTWL  234
TcCER1/3    TEFIYYWFHRALHHHSLYSQYHSHHHASFVTEAITGSVHPFLEHVGYTANFAIPLLGTWL  192
PcCER1/3    TEFVYYWFHRALHNHALYGAYHSHHHKSFVTEPTSGSCHPFMEHLGYTANFAIPMLGTWF  192
MkCER1/3    AEWVYYWGHRLLHHHYLYTRYHSHHHASFITEPVSGSVHPFAEHVMYTAMFAIPFLGTWA  196
            :*::*** ** ** * **  ****** **: *  :** *** **: *   **:* : *:
```

```
KnCER1/3      GGQGSIAMIYAYLLTFDFLNGLGHCNFEFIPTWFFKAFPPLKYLIYTPSYHSLHHTRVHT   258
OtCER1/3      LGGGDIAMFYTYLIGFDILNAIGHCNFEFVPRWF-MRLPGMKYLIYTPSYHSLHHSRVHT   252
McCER1/3      FGGASIAMFYAYLIGFDLLNNIGHCNFEFMPQWF-MNIPGVKYLIYTPTYHSLHHSKVHV   252
PsCER1/3      WGGASIAMFYTYLLGFDLLNMIGHCNFEFFPVWPFKYIPGLKYLIYTPSFHSLHHSRVHT   286
CcCER1/3      AGGASVAMIYTYLLGFDLMNIIGHCNFEIFPVWPFKVFPFLKYIIYTPSFHSLHHSRVVT   294
TcCER1/3      MGGASWTMFYAYLAAFDFLNAMGHCNFEFVPTGILKACPWIKYLIYTPTFHALHHAHVRT   252
PcCER1/3      LGGASMAMFYAYTIAFDTLNMIGHCNWEFVPVRLYQAVPLLRYLVYTPSYHSLHHSRVHT   252
MkCER1/3      LGGASIAMFYAYWLGFDFLNAIGHCNWEFMPSWLFQAFPPLKYLIYTPSYHSLHHSQVHT   256
               * .. :*:*:*   ** :* :****:*:.*       * ::*::***::*:***::* .

KnCER1/3      NFSLFMPLYDYLYGTVDNDSD-------TLYEQTIKG---VHPGPDAIFLAHGMDVGSFV   308
OtCER1/3      NFCLFMPLYDYVYGTADVTSD-------ELYEKAITGNAVPVKAPEVVFMAHGTELLSVF   305
McCER1/3      NFCLFMPIYDYAYGTNDPSSD-------ELYRKAINGEAAPNKAPDVVFVAHGTELLSLF   305
PsCER1/3      NFALFMPVYDWIGGTLDKKSW-------SLFESAAQGEAVPQKAPDAVFLGHGTEMLSVF   339
CcCER1/3      NFCLFMPMYDWMYGTLDPKSW-------DLFDKAAAGNAVQHAPPETVFLAHGTELLSVF   347
TcCER1/3      NFCLFMPIYDYIYGTWDPDSD-------PWQETAWKG---RQTEEGVVFLAHGTEMLSLF   302
PcCER1/3      NFCLFMPLYDYLGGTLEMERDGRTWTSDKLQAEAMAR---TYEVPDAVFLAHGTDLLSTF   309
MkCER1/3      NFALFMPLYDYLGGTVDGMSD-------LLHSSVRKG---RANQPDFVFLAHGTELLSLF   306
              **.****:**:  ** :                .             :*:.** :: * .

KnCER1/3      HGCFGIRGLASHPYYPSWVVYPLLPLATLLTIFCWLIDAKPWAVDRRSLRGNKIELWTVP   368
OtCER1/3      HLPFVLRSFSSRPFVSEWWLKPFWPLCVPFVLLLRVFG-KSFVADRHRLKTLNCETWVTP   364
McCER1/3      HLPFALRSFSSKPFKSVWWLQPFLPLCIPFVALLRIFG-KPFTADRHRLLHLNTATWVTP   364
PsCER1/3      HLPWMSRSFSAQPYEAQWWMYILWPIALVPLAVIRLVG-KALIQDKYRLGDFNCESWFTP   398
CcCER1/3      HLPFMSREFSSKPFQPSMWMYILWPIAVPMLLLARLFG-TVFVADKHRLGAMRIETWVTP   406
TcCER1/3      HLPFMTRAFASKPYQTTWAMYLLWPLTIPFLLVFWAFG-SVFVTDRNLLGKLKMQTWCTP   361
PcCER1/3      HLPMFFRGFAKNPYSESWLLYPLWPLAWLAMAVLWVVG-EAFPSHVERVNGHPVQTWVTP   368
MkCER1/3      HLPFGIPSFASRPYRPSWVMYLLWPLTLPIMAIIWLIG-KVFVSDSYTLESLRMQTWVVP   365
              *       :: .*:     :  : *:      .   ..      .   :       * .*

KnCER1/3      RMGLQYFFPSERPKINRLILQTLKKCDANGVKVFGLGALNKSEALNQGGEIYRKD---IE   425
OtCER1/3      AWGFQFFMKSEFNHINKKIEEAILDADKSGVQVVGLGALNKNEALNGGGALFVNKH--GK   422
McCER1/3      AWGFQFFIKSEFNHINRQIERAILEADATGTKVIGLGALNKNEALNGGGQLFVDKH--P-   421
PsCER1/3      VVALEFMFKSQWKSINSFIENAIVEADKSGVRVFGLGALNKNEALNGGGLQFIQKH--P-   455
CcCER1/3      AWAIDFFFKSQWGRINRAIDAAITDADKAGVKVFGLGALNKNEALNGGGALFVKNH--P-   463
TcCER1/3      RFGFQYFLKFDKPRINRLIKDAVIKADKLGLQVIGLGALNKAEPLNGGGKAILESALAER   421
PcCER1/3      RYGFQYFLKFERRRINGLIERAIRKADAQGVKVFGLGALNKAEHVNGGGVIFPEA---IK   425
MkCER1/3      RLGFQYFLSGEKKRINKHIEQAILDADELGVKVFTLGALNKNEALNGGGTLFVEK---HK   422
                .:::::  :   **  *  :: ..*  * :*. ****** * :* **    .
```

```
KnCER1/3      GLRMRVVHGNTLTAASILHRLPQDCDEIFLTGGTSKLGRALALYLVRKGVRVLLHTKSQE   485
OtCER1/3      SLKTRVVHGNTLTAAAILQKIPNDCKEIFLTGATSKLGRAIALYCAERGVRVVMYTTSEE   482
McCER1/3      NLRVRVVHGNTLTAAAILKKIPADVKEIFLTGSTSKLGRAIALYLSARGVRVVMYTTAKD   481
PsCER1/3      ELKMCVVHGNTLTAAALLQRINQDVKRVFVLGSTSKLGRAISLYLAKKGVKVMMMTNSRE   515
CcCER1/3      DLTMRVVHGNTLTAAAIIKTIPADVKKAFVLGATSKLGRGISLYLAKRGVTVVMLTQSEE   523
TcCER1/3      DLKVRVVHGNTLTAGCVIKELPAGVKEIFLTGSTSKLGRAIALYMSAKGTRVIMFTKSAE   481
PcCER1/3      DLRTRVVHGNTLTAACLVHELPADVREVFLTGATSKLGRAAALWLCRRGVRVRMLTSSVE   485
MkCER1/3      DLRVRVVHGNTLTAAVILDKIPKDAREIFLTGATSKLGRAIALYLCQRGVRVLMLTAARD   482
                *   *********. ::. :   .   . *: *.******. :*:   :*. * : * : :

KnCER1/3      RFQAIVDEAAPEHRKYLVWSPNLTDGSHCKNWVVGRWHSRKEQAFAPAGTHFHQFVVPLM   545
OtCER1/3      RFEMIRAEAPKKDQHLFVQSTSLTDGANIKDWVIGKHCSMKDQKSAPRGATFHQFVVPPI   542
McCER1/3      RFEKIKAEAREEHRELLVQATTLEEGSGIKDWVVGKFCSARDQAKAPKHATFHQFVVPPL   541
PsCER1/3      RYENTIEDCPKEYRGNLIHSTDMKDGANCKSWVVGRFLSKREQALAPSGTTFHQFVVPEL   575
CcCER1/3      R--------------------------VACWVVGRFLSKEEQHKAPKGAAFHQFVVPPL   556
TcCER1/3      RFAAVQEEAKLEHRHLLVHSTDYTDGSKCKTWVVGTLLGAKQQAFAPKGAFFHQFVVPPI   541
PcCER1/3      RFEAIRAEAAPEHRELLVRVEHHRDGGDCKTWIVGKMCGAKDQACAPPGTHFHQVVVPPI   545
MkCER1/3      RFEAIQREAPEECRHLLVQAESYAEGANCKSWVVGKWMTARQQAHAPPGTHFHQFVVPPT   542
              *                              *::*     .:*  **  : ***.***

KnCER1/3      TPARKDCTYSKQICMRMPKDTQGLGHCELNLQRNTVHACHAAVIVHLLEGWDFHELGSIN   605
OtCER1/3      PESRKDCVYTDLPAFKLPRESKDFRSCEMTMPRGHVHACHAGALVHALEGWDHHEVGAID   602
McCER1/3      EESRRDCAYTDLPAFKLPKEAKDFRSCEMTMERGHVHACHAGALVHALEGWTYNEVGAID   601
PsCER1/3      ERVRTDCSYTALPAFTLPKMAQGLRSCEMTMGRRNVHACHAGALVHLLEGWKHHEVGAID   635
CcCER1/3      DELRKDCVYTKLPAFTLPENAAGFKSCEMTMQRRDVHACHAGALVHALEKWTHHEVGAID   616
TcCER1/3      SETRSDCSYGKLAAMRLPPK-TLMRACEMTMERNCVHACHAGALVHALEGWEHHEVGAID   600
PcCER1/3      AKHRSDCTYGELAAMDLPDTATDVKACELFMPRRRVFACHAGALTHMLEGWDHHEVGAID   605
MkCER1/3      PEVRKDCTYGKLAAMRLPNTVQNMRTCEMTMDRGCVHACHAGGLVHALEGWDHHEVGAID   602
                 * ** *    .: :*      .   **: : *  *.****. :.* ** * .:*:*:*:

KnCER1/3      VDRIDENWEAALKHGFEPVS--------------------------------   625
OtCER1/3      HTRIDLTWEAALKHGFSLA---------------------------------   621
McCER1/3      HTKIDVTWDAAVKHGFALA---------------------------------   620
PsCER1/3      PSRIDATWEAAMRHGFKLVEEEDA----------------------------   659
CcCER1/3      EEKIDRTWDAAMKHGFKMVC--------------------------------   636
TcCER1/3      VSRIDLTWKAAMSHGFKPVFESQAAPVKEAAAAAA-----------------   635
PcCER1/3      IDRIDVTWEAALRHGFRPAVAPAGCAVGCAQPAAVAARASDAAAIPAAVAAA   657
MkCER1/3      PSKIDVTWEAALRHGFQPV---------------------------------   621
                 :** .*.**: ***   .
```

**Figure S1: Sequences alignement of different algal CER1/3 homologs.** The algal CER1/3 homologs compared comes from *Klebsormidium nitens* (*Kn*CER1/3), *Ostreococcus tauri* (*Ot*CER1/3), *Micromonas commoda* (McCER1/3), *Proteomonas sulcata* (*Ps*CER1/3), *Cryptophyceae sp. CCMP2293* (*Cc*CER1/3), *Tetraselmis convolutae* (TcCER1/3), *Prasinoderma* *coloniale* (*Pc*CER1/3), *Mesotaenium* ktamstae (*Mc*CER1/3). Identical aminoacid residues are indicated by asterisks (*), closely related aminoacids are indicated by colons (:) and distantly related aminoacids are indicated by dots (.). The NTDs of the different CER1/3 homologs are highlighted in yellow while the CTDs are highlighted in blue. The histidine-rich motifs are in red bold, the putative NADPH binding sites are in black bold and highlighted in green, and the putative catalytic cysteines are in red bold and highlighted in green. The alignment was performed byu using Clustal Omega (https://www.ebi.ac.uk/jdispatcher/msa/clustalo).

```
AfCER1          ---------MEWMWMWMVLGRRRRRRQYLAYVLVGGAALW-----EGSPWTTNISLNLLL 46
ZmCER1          -MASKPGPLSRWPWQD--LGNY----KYALVAPWAVRSTYRFVRSGSG-ERDLLAFAVLP 52
NoCER1          -MASHPGPLTDWPWEK--LGSY----KYAVLLPFIGHAVHTLYNS-EPADRDYTLLCPFY 52
AtriCER1        -MASKPGPLTDWPWKK--LGNF----KYMVIVPCAIHAVYRIYGA-QKQKRDHATIISLG 52
AcCER1          ----SIHDHFLFPLHM--HACM----QYVVLAPWVLHGLRMVATKGWRELDVITYNAIFP 50
AtCER1          -MATKPGVLTDWPWTP--LGSF----KYIVIAPWAVHSTYRFV-TDDPEKRDLGYFLVFP 52
SlCER1          -MASKPGILTEWPWTW--LGNF----KYVVLAPFVGRSIESLLNREDGSKIDIGYLIIFP 53
PpCER1          MKAIQPGAWTKFPWHS--MGDC----KYLLYLSLVGRLLYG-LIREDR-GRYDLYFHVLL 52
SmCER1          -MAINPGLLTHWPWER--LGSF----KYLLYLPLVANAVRSAMTPEGR-SRDNFSLHILV 52
                           :       .      :*                              :

AfCER1          LATLRYTTFQIWGSVSRLPFFYRPFIIQKKSIPIDQVEHEASWENSVIFTTLLASSFKVM 106
ZmCER1          VLLLRLLYSQLWITVSRHQTARSRHRIVNKSLDFEQVDRERNWDDQILLTALLFYAVNAA 112
NoCER1          LLVTRLVHDQLWISWSRFQNARSKHQIQSRGIEFQQVDRERRWDDQAIMHALAIYVAHVF 112
AtriCER1        LFLSRIIHDQLWITLSRFQTARSKHRIQSRGIDFEQVDRERNWDDQAILHAIIFYIGELF 112
AcCER1          SLLLRMIHNQIWITLSRFQNARSKHQIVDRSIEFEQVDRERNWDDQIIFNGLLLYVGFTF 110
AtCER1          FLLFRILHNQVWISLSRYYTSSGKRRIVDKGIDFNQVDRETNWDDQILFNGVLFYIGINL 112
SlCER1          FLLFRMLHNQIWISLSRYKTAKGDNRILDKTIEFDQVDRERNWDDQILLNGLLFYYGYMK 113
PpCER1          LAVLRHFFGQLGISLSRWPYLSSRYQIQKKGFSFDAVDLSSNWDDYIILDTLLLSVTVMI 112
SmCER1          LAALRYIQGQLWITVTSVHDIVKKHQVQTKGMKFDQLDRERDWEDFILLQALMLLAYQFS 112
                    *    *:  : :          :  : : :: :: .  *::  ::  :

AfCER1          FPSI-FTLPLWDGKGVALALLLHVGPVEWLYYWGHRALHHHYLFSRYHTHHHSSFVTQPV 165
ZmCER1          VPVA-QSVPWWDSRGLLVAALLHVGPVEFLYYWLHRALHHHYLYARYHSHHHASIVTEPI 171
NoCER1          IGGA-SHLPLWNSKGLLFTALVHAGPVEFVYYWAHRALHHYWLFTRYHSHHHSSFVTEPI 171
AtriCER1        LPGA-QNLPLWNTKGVIIAVLCHAGPVEYIYYWAHRALHHHFLYTRYHSHHHSSFVTEPI 171
AcCER1          IPSSAQRLPMWRTDGAVMIALLHAGPVEFLYYWFHRALHHHFLYSRYHSHHHASIVTEPI 170
AtCER1          LPEA-KQLPWWRTDGVLMAALIHTGPVEFLYYWLHKALHHHFLYSRYHSHHHSSIVTEPI 171
SlCER1          LEQS-HYLPIWRSDGIILTTLLHIGPVEFLYYWLHRALHHHFLYSRYHSHHHSSIVTEPI 172
PpCER1          PMFGNRYYPPWDWTGLVICALLHMGPAEAIYYWLHRALHGHYLYTRYHSHHHSLFVTEAN 172
SmCER1          PLCL-PNHAVSDWRGLVITILWHLGPVEFLYYWFHRALHHHSLYRRYHSHHHLSFVTQAV 171
                              *  .  * * **.* :*** *:*** : *: ***:***  :**:

AfCER1          SANNHPMLEIMFYTMLFAIPIVGAVLVGSASIAMYHIYVFFVDTMNAWGHCNFEFIPTSV 225
ZmCER1          TSVIHPFAEELVYFTLFAIPLLTMVGTGTASVAVANGYLAYIDFMNYLGHCNFELVPRLL 231
NoCER1          TSVVHPFAEHLLYLAIFAPGFVVPWITGTGSLIALCGYMTFIDLMNNLGHCNFEFIPKWA 231
AtriCER1        TSVVHPFAEHILYALIFAVPFLATVLTYTASYTVLIGYMTFIDFMNNMGHCNFEFFPKWT 231
```

```
AcCER1      TSVIHPFAEHVVYYALFTIPMLTTVLTGRGSIIALLAYTTYLDFMNNMGHCNFELVPKWL 230
AtCER1      TSVIHPFAEHIAYFILFAIPLLTTLLTKTASIISFAGYIIYIDFMNNMGHCNFELIPKRL 231
SlCER1      TSVIHPFAEHIAYFLLFSIPLLTTVVTKTASIVSFGGYITYIDFMNNMGHCNFEIIPKWM 232
PpCER1      SGTVHPFLEHLMYASNFAIPLFGTWALGRFSISTLYVYTLTFDTLNAIGHCNVEFVPSWL 232
SmCER1      TGNVHPFAEHLSYAVLFGSTLIVNLFLGTASLALIYSYMLWFDFMNYIGHCNWEFMPSWM 231
            :.   **: * : *    *    :.          *        *    .* :*   **** *:.*

AfCER1      YDAFPPLKYLIYSPSFHSLHHSKVHTNFALFMPLYDYMYGTADPTSDELHRQVREGN--H 283
ZmCER1      FDVFPPLKYLMYTPSFHSLHHTQFRSNYSLFMPLYDHLYGTADKSSDDLYERALQGRAGE 291
NoCER1      FTIFPPLKYIMYTPTFHSLHHSQVHINFCLFMPIYDYMYGTVDKTTDTLYETSISGR--E 289
AtriCER1    FKIFPPLKYLIYTPSYHSLHHSQVHTNFSLFMPIYDYMYNTVDKTTDSVYESSIEGR--E 289
AcCER1      FHAFPPLKYLMYTPSFHSLHHTQFRTNYSLFMPFYDYIYGTMDKSSDDLYESSLKGK--E 288
AtCER1      FHLFPPLKFLCYTPSYHSLHHTQFRTNYSLFMPLYDYIYGTMDESTDTLYEKTLERG--D 289
SlCER1      FSTFPPLKYLMYTPSYHSLHHTQFRTNYSLFMPIYDYIYGTLDKSSDTLYEKSLERQ--G 290
PpCER1      FDAFPPLKYLIYTPSYHSLHHSQVHTNFCLFMPIYDYWGGTMDKNSDALYRSVRR-SDSQ 291
SmCER1      FQALPLLKYLVYTPSFHSLHHTQVHTNFCLFVPLYDYIYGTVDKTSGQLHLAAR--QGRT 289
            :   :* **:: *:*::*****::.: *:.**:*:**:   .* *  .:. ::

AfCER1      DKVDCVYLTHPTDLPSIFHQPFAFQTFATQPY-SKPW-------------YVWLTWPFSF 329
ZmCER1      DAPDVVHLTHLTTPASLLRLRLGFASLAAAPA-PPASRYGAGSSSSSSSSLAAVACPLA- 349
NoCER1      QMTDVVHLTHPTSIHSIWQIRFGFAYLAAEPY-STKW-------------YFWLLWPFTA 335
AtriCER1    DTTDVVHLTHPTSLHSIYHLRLGFAYLAAEPY-SSKW-------------YLWLMCPLSF 335
AcCER1      EAPDVVHLTHPTTLQSIYHLRLGFASLASRPY-NSKW-------------YMLIMWPISS 334
AtCER1      DIVDVVHLTHLTTPESIYHLRIGLASFASYPF-AYRW-------------FMRLLWPFTS 335
SlCER1      KSPDVVHLTHLTTPESIYHLRLGFASFASQPY-TSKW-------------YFWLMWPVTL 336
PpCER1      ERADNVYLTHGMDLLHMMHVTLGIQSFAATPYKGPNW-------------RLWLLYPLAL 338
SmCER1      ELVDFVFLTHPTDPLSIFHLSFGIPSFAAQPY-GRRW-------------YIWLLYPLAL 335
            .   * *.***       : :   :.:   :*: *                         :   *.:

AfCER1      VIMVAFWLFAKPFVATSQLIGKSFELQTWMIPRYNFQYRMKPVSAKIKRLVEQCVLDAQE 389
ZmCER1      --ALLGWT-RTAFRSEANRLHK-LKLETWVVPRYTSQYLSKQGLYAVGRVVEKAVADAEA 405
NoCER1      ALALLTWMFGATFTVEKIRLDK-LKIQTWAIPRFGFQYNVASQKQPINSMIKKAIKDADS 394
AtriCER1    VLMLVTWIFGCTFVVEKNKLYE-HKMQTWAIPRYNFQYSLKWQKAPINNMIEKAILEADA 394
AcCER1      VSMLLTWIYGSSFTVERNTLKK-LKLQTWAIPRYNFQYGLKWEKEAINDLIEKAILEADQ 393
AtCER1      LSMIFTLFYARLFVAERNSFNK-LNLQSWVIPRYNLQYLLKWRKEAINNMIEKAILEADK 394
SlCER1      WSMMVTWIYGHTFTVERNVFNN-LNLQTWAIPKYRVQYFMQWQRETINNLIEEAIMEADQ 395
PpCER1      IAMPLLWILGQPFAADKYWIPRTLRGETWLIPRYRFHYSLPVEKVRINALIEQAIVMAED 398
SmCER1      PVMLLLWAFGSPFTVEEHTVDK-VLAQTWAIPRFSFHFGMTSEIGSLNALIERAILAAQD 394
                          *        . .     ::* :*::   ::          :   :::..:   *:

AfCER1      AGAKVISLGLLNKELL----------DGD--FLRTRSIEIPIVTGETLTSAIIVKKIEAQ 437
ZmCER1      SGARVLTLGLLNQA-------NELNKNGELYVIRKPSMRTKIVDGTSLAAAAVLHMIPEG 458
```

```
NoCER1      KGVKVITLGLHNQS-------EELNENGKSYLDSVGNMKVKVVDGCSLAAAVVMNNIPQG 447
AtriCER1    MGVKVISLGLLNQG-------EEFNKNGELYLHNREKLKTRIVDGSTLSAATVLNGIPHG 447
AcCER1      RGVRVLSLGLLNQACYSLALAKELNGSGELYIEKHPKLRVRIVDGSSLAAAVVLKSIPAG 453
AtCER1      KGVKVLSLGLMNQG-------EELNRNGEVYIHNHPDMKVRLVDGSRLAAAVVINSVPKA 447
SlCER1      KGIKVLSLGLLNQD-------EKLNKNGEVYIRRHPQLKVKLVDGSSLAVAVVLNSLPKG 448
PpCER1      EGCRVVSLGQLNKE-------MRLNGSGAAIVVRNPHLKVRIVTGLTLTAAVVINRLPKQ 451
SmCER1      KGAKFICLGLHNKD-------EHLNASGALFLKNHPDLSIKVVDGSTLTSAIVLDKLPKD 447
             * :.: **  *:             .*   .     :   :* *  *: * ::. :

AfCER1      KVDKVFLASANS--RVGQAVATYLCQHGVTVVALLPTKSCFHELQANIDAEFQKNLVFAT 495
ZmCER1      T-DEVLLLGDAGGNKMAGVLASALCEREIQVHVVDK--DLYESVKQQLRPETHEHLLHLA 515
NoCER1      A-RQVLVCGRLT--KTGYAVVRALCQRGTKVLTVTE--EELQGLKSKIPAELHDRFEL-- 500
AtriCER1    T-QKVLIRGCLT--KTLFTTALALLERRTKIVVIRT--EEYENLKMRIPSKYQSGISL-- 500
AcCER1      T-NEVLLAGNLS--KVARAVAAALCQKGVQVIMTRK--HEFKMLKGQMTEKAANYLVF-- 506
AtCER1      T-TSVVMTGNLT--KVAYTIASALCQRGVQVSTLRL--DEYEKIRSCVPQECRDHLVYLT 502
SlCER1      T-TQVVLGGHLS--KVANAIALALCQGGVKVMTLRE--EEYKKLKSSLTPEAATNLLL-- 501
PpCER1      T-KEVFLVGSSD--LIR-SVEIYLVRRGVRVLVLTNSPRYFGSTQPKVTKVNQQLIVNVM 507
SmCER1      A-SEVFLVGAEH--KVGRAIANYLCRHRAT---------EVTSLKKSVPQESQHKLVAVE 495
               .*.: .              * .                  :  :       :

AfCER1      TY--KNGKDCKVWVVNDMVAAEEQRWAPAGTCFHHLGVFPLSTTTRTSDCTYEYIPAMRL 553
ZmCER1      EWWSHSAKTTKVWLVGDRLTGEEQRKAQGGAHFVPYSQF-PPGAVVRADCVYHSTPALVV 574
NoCER1      ----VHYYDCKVWLVGDGLSTQVQRKAPKGTLFVPFSQF-PTKA-VRSDCTYHTTPAMAI 554
AtriCER1    ----SNNYDTKVWLVGEDLRAEEQSRANRGTLFIPFTQF-PPRA-VREDCIYHTTPALVI 554
AcCER1      ----SRNYTTKVWLVGDGLEHEEQKRAPKGARLIPYSQF-PPKK-IRNDCSYYTTPCMKI 560
AtCER1      ---SEALSSNKVWLVGEGTTREEQEKATKGTLFIPFSQF-PLKQ-LRRDCIYHTTPALIV 557
SlCER1      ----SKTYTSKIWLVGDGLNEDEQLKVPKGTIFIPFSQF-PPRK-TRKDCFYFHTPAMIT 555
PpCER1      SF--QEGQHCREWILDEYVEGKDLKWAPPGADLHHVCQGSKPLPRTRKDCTYAMYPAMHV 565
SmCER1      SL--EHGRHCKAWIVGEPLRAMEQLHAPSGACFYQFT--EEAMEETRPDCLYAKLPAMRL 551
                      : *::.:         .  *: :               ** *   *.:

AfCER1      PSNARTMNGCEDVLPKYVVRAAYAAGMLHALEGWKHSDFSNEISTYLLDKTWEAAVKHGF 613
ZmCER1      PDAFEDLHACENWLPRRVMSAWRAAGIVHALEGWDAHECGAR--VTGVDKAWRAALAHGF 632
NoCER1      PKALENVHSCENWLPRRVMSAWRIAGIVHALEGWDAHECGEK--MLDMKKVLDAAVSHGF 612
AtriCER1    PRTLEDVHSCENWLPRRVMSAWRVAGIVHALEGWDSHECGNM--TLDVEKVWSTTLSHGF 612
AcCER1      PETLQNMHSCENWLPRRVMSAWRAAGIVHALEGWDVHECGET--VQDVEKTWSAAIRHGF 618
AtCER1      PKSLVNVHSCENWLPRKAMSATRVAGILHALEGWEMHECGTSLLLSDLDQVWEACLSHGF 617
SlCER1      PKHFENVDSCENWLPRRVMSAWRIAGILHALEDWHEHECGNL--MFDIEKVWKASLDHGF 613
PpCER1      PKSMKGLRSCEGGLPRGVISASHAAGVVHSLEKWTHNEVGP-IDVERIDTVWAAALKHGF 624
SmCER1      PPEYKGIRACEGSMPRGVVQASHAGGILATMENWNHHEVGNTIDVDKIDAVMRAAVNRGF 611
            *     : .**. :*: .: *   .*:: ::* *   : .       :. .  : : :**
```

```
AfCER1      SIYSLNGDDQAKH   626
ZmCER1      RPYDRYGAN----   641
NoCER1      RPLGVVRSM----   621
AtriCER1    KPINNVKFN----   621
AcCER1      LPVTQI-------   624
AtCER1      QPLLLPHH-----   625
SlCER1      QPISVVSASESKA   626
PpCER1      QMAV---------   628
SmCER1      VPYY---------   61
```

**Figure S2: Sequences alignement of different plant CER1 homologs.** The plant CER1 homologs compared comes from *Azolla filiculoides (Af*CER1), *Zea mays* (*Zm*CER1), *Nymphaea odorata* (*No*CER1), *Amborella trichopoda* (AtriCER1), *Ananas comosus* (AcCER1), *Arabidopsis thaliana* (AtCER1), *Solanum lycopersicum* (*Sl*CER1)*, Physcomitrium patens* (*Pp*CER1) and *Selaginella moellendorfii* (*Sm*CER1). Identical aminoacid residues are indicated by asterisks (*), closely related aminoacids are indicated by colons (:) and distantly related aminoacids are indicated by dots (.). The NTDs of the different CER1 homologs are highlighted in yellow while the CTDs are highlighted in blue. The histidine-rich motifs are in red bold. The alignment was performed by using Clustal Omega (https://www.ebi.ac.uk/jdispatcher/msa/clustalo).

```
SmCER3      -------------------MEESKILGCWPWQRMGTYKYHLFLPIFLSAAHSHY---LGT 38
PpCER3      ------------------MVAKEAFLAEWPWERLGHFKYLVYAPFVGRLLQTII--YGTG 40
AfCER3      ----------------MALPSAPAPLYSWPWRWMGCFKYLLYLPIAWAAYSSLE--Q-KT 41
SlCER3      MEKQNEEVENGND---TLLRKTDAALFAWPWNNLGNYKYLLYGPFLAKFIHSMY--WKES 55
AtCER3      ---------------------MVAFLSAWPWENFGNLKYLLYAPLAAQVVYSWV--YEED 37
NoCER3      ---------------------MVAPLSAWPWNELCNLKYFLYAPLLANCITSWG--NDEA 37
AtriCER3    ------------------------------------------------------------ 0
AcCER3      ---------------CDKAKMTSAPFSSWPWENLGVYKYVLYGPLIAKAFTSKAWEFG-- 43
ZmCER3      MGK--PRIIQGKQTTTRAAALLILGSLLFVWTGV-HAQYLLYGPLVAKVA--HAWRETGS 55

SmCER3      SPRDNWCFHILVIAALRYALYQAWSSFARLHAVVKHHQIISYALTYEQVDREFDCDNGII 98
PpCER3      LEPDNWAMHMFFLMVARYFHQQLWVSASRVPWLTEKFVVDERQSGYEQVDREYHSDNHLM 100
AfCER3      PISQNFSLHILILTAARLILYQLWQTFSRFLPLTESIQIRPQGLSYAQVDREWHCDDHHL 101
SlCER3      M-EDIWCLHILVLCSLRGLVHQLWSTFSNMLYLNSTRRVSYEGIDYDQIDNEWDWDNFLI 114
AtCER3      ISKVLWCIHILIICGLKALVHELWSVFNNMLFVTRTLRINPKGIDFKQIDHEWHWDNYII 97
NoCER3      ---GGWCFHILALCALRGVVHQFWYSYSCMLFITDKYRVLKQGVDYKQIDREHNWDNFII 94
AtriCER3    -----------------------------MLFLTKKLRVLQQGVDFKQIDREWHWDNFIL 31
AcCER3      -NPNKWCLHILILFALRGAVHQLWYTFSNMLFLTRRRRIFTDSVDFDQIDKEWDWDNFLI 102
ZmCER3      LPLGSWCLHLLLLLALRSLTFQLWFSYGNMLFFTRRRRVVKDGVDFRQIDAEWDWDNMVI 115
                                             .  .     :      : *:* * . *:  :
SmCER3      LHSLLAYAL-----GPN---DISGFSIWNLRGLVYLIAFHAGVTESAYYWLHRAFHT-KS 149
PpCER3      LQLIFISV--AHSWFPGF----SNVVAWNTQGFLYVLLFHVGVVEVLYYWIHRAFHT-EV 153
AfCER3      LQALVLFG------AHSWLGFLKGAPVWRGSGLLMTLLLHTGPAEMIYYWFHRFLHS-AP 154
```

```
SlCER3      LQAVVGSF--VYLNFPSL----ANLPVWDVRGLISCLILHIGISEPLFYWMHRLLHS-SY 167
AtCER3      LQAIIVSL--ICYMSPPLMMMINSLPLWNTKGLIALIVLHVTFSEPLYYFLHRSFHRNNY 155
NoCER3      LHAFMAAI--ACYS-STF---MDSLPLFNFRGYIYALILHMGITESLYYFIHRMFHS-DY 147
AtriCER3    LQALMATF--ACYYFPSF---TKSLPLYNTTGFICVLLLHMGFSESLYYGVHRLFHS-DY 85
AcCER3      LQCLIGAM--SLYTFPAL----RDLPQWDVWGILLAFFLHVTISEPLFYFVHRAFHR-GH 155
ZmCER3      LQTLVAAVAMGSAAFPAV----SELRAWDPRGWALALLLHMAVSEPVFYWTHRALHR-GP 170
            *: ..                      :   *    : :*    *  :*  ** :*

SmCER3      LFRSFHSYHHASTAPEPATAFTHTFLEALLQTVLMSVPIFASCFLGGSCLALFYVYPLAF 209
PpCER3      LFRNYHFYHHMSVVPEPPTGSITTMLEQILQSLLVCVPLLGAAALGGGSMAMIYIYLIAF 213
AfCER3      MFERYHKLHHESVTPEPSTPGLSTFLEQISMVVIMAVPIVGGSVKGQGSLSALYAYVLLF 214
SlCER3      LFPLYHWHHHESKITHPFTAGHGTFLEHLLLCVVIGIPTLGTAFIGYGSISVMYSYILAF 227
AtCER3      FFTHYHSFHHSSPVPHPMTAGNATLLENIILCVVAGVPLIGCCLFGVGSLSAIYGYAVMF 215
NoCER3      LFQNYHSLHHLSVVAQSYTAGTASILENLVMGFLMGIPMLGASWMGGASVTMFYGYVLVF 207
AtriCER3    LYTNYHSFHHKSMVSQSFTAGSASLLEHIVLCFIMGIPIVGASWMDCASLSMVYAYIMAF 145
AcCER3      LFSLYHSLHHSSKVPQSFTAGFATPLEHLILSVVMGVPLLVPCLVGNGSLGLIYGYVLLF 215
ZmCER3      LFSQYHARHHSSPVTQPFTAGFGTPLEALLLTLAMGAPLAGAFLAGAGSVSLVYGHVLLF 230
            ::  :*  ** *   .  *    : ** :   .    *       . ..:  .* : : *

SmCER3      DFFKYLGHFNCEIVPLWAFQKLPLLKYLIYTPSYHSLHHLDLKSNFCLFMPLYDYLGGTQ 269
PpCER3      DFFKCWGHSNFEFVP-EWFRGFPGVKYLLYTPSYHSLHHLEQNSNFCLFMPLFDYLGGTV 272
AfCER3      DCMRCMGHCNAEMFSPTFLANFPLIKYLIYTPSYHTLHHEERDSNYCLFMPIYDYMFGTV 274
SlCER3      DFLRCMGHSNVEIIPHSLYQRVPLLRYVIYCPTYHSLHHQEMKTNFCLFMPLYDMLGNTL 287
AtCER3      DFMRCLGHCNVEIFSHKLFEILPVLRYLIYTPTYHSLHHQEMGTNFCLFMPLFDVLGDTQ 275
NoCER3      DFIRCMGHCNVEVMPVSLFDHMPVLRYLIYTPSYHSLHHTDMSSNFCLFMPLYDALGNTL 267
AtriCER3    DFMRCLGHCNVEIIPHSLFEWMPLLRYIFYTPTYHSLHHNEKSTNFCLFMPIYDALFNTL 205
AcCER3      DFLRCMGHSNVEVFPHKLFEALPLLKYFIYTPTYHSVHHMEKNSNFCLFMPLFDLLGGTL 275
ZmCER3      DCLRCLGYSNVEVISHRAFAAFPPLRYLVYTATYLSLHHREKDCNFCLFMPLYDALGGTI 290
            * ::  *: * *..       .* ::*..*  :* ::** :   *:*****::* : .*

SmCER3      HPNTHAFYRSIR--------------------------KDGREAVPQFVFLVHCIDILSS 303
PpCER3      DPKTESLYAELR--------------------------KGRLLKVPDFVFLAHCIDVLSS 306
AfCER3      SKNREAFLKEIR---------------------------ERVLRVPDYVFLLHAVDFLSC 307
SlCER3      NTASWSLHKEIS--------------------------SRTNERAPDFVFLAHIVDIMSS 321
AtCER3      NPNSWELQKKIRLS------------------------AGERKRVPEFVFLAHGVDVMSA 311
NoCER3      NKNSWEFHRRIR--------------------------TGEDVRVPDFVFLAHVIDVHAS 301
AtriCER3    NPKSWDLHRTVR--------------------------SGCGDKVPDFVFLGHVIEFDSA 239
AcCER3      DNKTWKLHKEISLEVFTSLVSTVVHVWRMVTEERSKFLAGRNDQVPDFVFLAHVLDLVSA 335
ZmCER3      SSRSWGLQREVD--------------------------QGMNDRVPDFVFLAHVVDVVSS 324
                  :   :                                 .*::*** * ::. :.

SmCER3      LHVAFSGRTASSVPFRGEWYAWLVFPIGLVSCFCVWIWGKTFVATKYLLDGLHAQSWVVP 363
```

```
PpCER3      LQVSFCCRTMAAHPYKCHWFIWWTWPITVFFLMIFWYWGQTFTAMTIYVNKLKCTSWVIP 366
AfCER3      LHVHFLFRAFASHGYRTLLFLIPIWPFVIPIAIAMWIWGKPFKGWNYYLDNRFHQIRLIP 367
SlCER3      MHAPLLFRSFSSVPFSTRLFLLPMWPFAFVVVLTMWLKSKTFLFSFYNIRGRLNQTWIVP 381
AtCER3      MHAPFVFRSFASMPYTTRIFLLPMWPFTFCVMLGMWAWSKTFLFSFYTLRNNLCQTWGVP 371
NoCER3      MHAPFMFRSVSSKAFKANLLVLPLWPVALVVLQLMWAWAKPFLVSFYCLRNRLLHTWIVP 361
AtriCER3    LHVSFLFRSISSMPCVFRPILLPTWPVVFVVMLFMWASATTFLISFYTLRDRLHQTWVIP 299
AcCER3      MHVQFIFRWHSSMPFAVKPLLLVFWPVAIVIMLCMWAWSKTFLVSMYRLRGRLHQIWAVP 395
ZmCER3      MHVPFAFRSCSSLPWAMRPVLLPLWPVAFAFMLLQWFFSKTFTVSFYFLRGRLHQTWSVP 384
            ::. :  *  ::            :*. .      *  .  *      :         :*

SmCER3      RYGFHYFIPACAAGINRHIERAILDADELGVKVISLAALNKNESLNGGGLLFVKKHPNLK 423
PpCER3      KHGFQFFLPFGLDSINKHIEKAILEADKQGVKVISLAALNKNEALNGGGLLFVKKHPNLK 426
AfCER3      RFGFMYFLPFARNNINNLIEKAILDADRMGVKVLSLAALNKNEALNGGGNLFVMKHPNLN 427
SlCER3      RAGFQYFLPFAAEGINKLIEEAILRADRIGVKVISLAALNKNESLNGGGTLFVNKHPNLR 441
AtCER3      RFGFQYFLPFATKGINDQIEAAILRADKIGVKVISLAALNKNEALNGGGTLFVNKHPDLR 431
NoCER3      RYGFQYFLPFAKDGINRSIEEAILRADKIGVKVISLAALNKNEALNGGGTLFVGKLPHLR 421
AtriCER3    RFGFQYFLPFAKDGINKHIEDAILSADKLGVKVISLAALNKNEALNGGGIIYVQKHRNLK 359
AcCER3      RFGFQYFLPFAKDGINHHIELAILRADKMGVKVLSLAALNKNEALNGGGMLFTSKHQDLR 455
ZmCER3      RYGFQYFIPSAKKGINRQIELAILRADKMGVKVISLAALNKNEALNGGGTLFVNKHPNLR 444
            : ** :*:*    .**  ** *** **. ****:*********:***** ::. *  .*.

SmCER3      VRVVHGNTLTAALVLRELPAETSEVFLTGSTSKIGRAIALYLCRRNVRIMMLTTSRERYQ 483
PpCER3      VRVVHGNTLTAAVIIKTLPPDVKEVFMTGATSKLGRAIALYLCARGIRVLMLTTSTERFD 486
AfCER3      IRVCHGNTLTAAVIIKEIPDHVTEVFLTGATSKLGRAIALYLCQKGVRVLMLTESRKRFE 487
SlCER3      VRVVHGNTLTAAVILNEIPRDVNEVFLTGATSKLGRAIALYLARRRVRVLMLTKSTERFM 501
AtCER3      VRVVHGNTLTAAVILYEIPKDVNEVFLTGATSKLGRAIALYLCRRGVRVLMLTLSMERFQ 491
NoCER3      VRVVHGNTLTAAVILHEIPQDVKEVFLTGATSKLGRAIALYLCRKAVRVMMLTHSTERFQ 481
AtriCER3    VKVVHGNTLTAAVILNELPNGVKEVFLTGSTSKLGRAIALYLARKGVRVLMLTLSQERFQ 419
AcCER3      VRVVHGNTLTAAVILNEIPKDVQEVFLTGATSKLGRAIALHLCRKKIRVLMLTLSTERFQ 515
ZmCER3      VRVVHGNTLTAAVILNEIPSSVREVFLTGATSKLGRAIALYLCRKRIRVLMLTLSTERFL 504
            ::* ********:::  :*  . ***:**:***:******:*. : :*::*** * :*:

SmCER3      SIVDEAPADCRHNLVQVTKYQAGQTCKTWIVGKWATSQDQSWAPHGSHFHQFVVPPVHEY 543
PpCER3      AIQREAPADCRNNLIHVTKYQAGKNCKTWIVGKWTFAKDQQWAPPGTFFHQFVVPVISEV 546
AfCER3      SITSESPAEFRHNLVQVTKHQAGKNCKTWILGKWTTYSDQMFAPPGTHFHQFVVPPVIPF 547
SlCER3      KIQREATVECQKYLVQVTNCKEAKQCKTWIIGKWSTPREQSWAPSGTHFYQFVVPPIIPF 561
AtCER3      KIQKEAPVEFQNNLVQVTKYNAAQHCKTWIVGKWLTPREQSWAPAGTHFHQFVVPPILKF 551
NoCER3      AIQKEAPTECQKFLVQVTKYQVAQNCKAWIVGKWLSPMEQAWAPSGTHFHQFVVPPILEL 541
AtriCER3    IIQKEAPVEYQKNLVQVTKYQAGQNCKTWIVGKWILPREQKWAPSGTHFHQFVVPPILRL 479
AcCER3      KIQKEAPPQFQQYLVQVTKIQAAKNCKTWIAGKWLSPREQLWAPPGTHFHQFVIPPIIGF 575
ZmCER3      KIQREAPPEFQQYIVQVTKYQAAQGCKTWIVGKWLSPREQRWAPPGTHFHQFVVPPIIGF 564
```

```
                     *  *:  : :: :::**: : .: **:** ***    :* :** *:.*:***:* :
SmCER3               RKDCTYGKLAGMKLPQS-VEGVHSCEYTFDRGVVAACHAGGLVHALENWTHHEVGSIDID 602
PpCER3               RKDCTYGQLAGMVLPKEGVKGLRTCEFTMERGAVHACHAGGMIHTLEGWTHHEVGSIDVS 606
AfCER3               RRDCTYGALAAMRLPPN-TKGLASCEYTLDRNVVHACHAGGVVHALEGWTHNEVGALDVD 606
SlCER3               RRDCTYGKLAAMRLPKD-VTGLGTCEYTMGRGIVHACHAGGLVHLLEGWTHHEVGAIDVD 620
AtCER3               RRNCTYGDLAAMKLPKD-VEGLGTCEYTMERGVVHACHAGGVVHMLEGWKHHEVGAIDVD 610
NoCER3               RRDCTYGKLAAMRLPDD-VEGLGNCEYTMGRGIVHACHAGGLVHLLEGWEHHEVGAIDLD 600
AtriCER3             RRDCTYGKLAAMHLPDD-VEGLGTCEYTMGRGIVHACHAGGVVHLLEGWTHHEVGAISVD 538
AcCER3               RKDCTYGRLAAMRLPKD-VQGLGMCEYTLERGVVHACHAGGVVHLLEGWTHHEVGPIDVD 634
ZmCER3               RRDCTYGKLAAMRLPKD-VEGLGSCEYTMERGVVHACHAGGVVHCLEGWEHHEVGALEVD 623
                     *::**** **.* ** . . *:  **:*: *. * ******::* **.* *:*** :.:.

SmCER3               HIDLVWEAALKHGLEPVL----    620
PpCER3               RIDVVWEAAMRHGFAPIGS---    625
AfCER3               RIDLVWEAAMRHGFRPA-----    623
SlCER3               QIDVVWEAALKHGLKPLYNSS-    641
AtCER3               RIDLVWEAAMKYGLSAVSSLTN    632
NoCER3               RIDVVWECALKHGLRPLS----    618
AtriCER3             RIDLVWEAALKHGLKPV-----    555
AcCER3               RIDVVWRAALKHGLAPQLHY--    654
ZmCER3               RIDVVWEAALKHGLTPA-----    640
                     :**:**..*:::*:
```

**Figure S3: Sequences alignement of different plant CER3 homologs.** The plant CER3 homologs compared comes from *Selaginella moellendorfii* (*Sm*CER3), *Physcomitrium patens* (*Pp*CER3), *Azolla filiculoides (Af*CER3), *Solanum lycopersicum* (*Sl*CER3), *Arabidopsis thaliana* (*At*CER3), *Nymphaea odorata* (*No*CER3), *Amborella trichopoda* (*Atri*CER3), *Ananas comosus* (*Ac*CER3) and *Zea mays* (*Zm*CER3). Identical aminoacid residues are indicated by asterisks (*), closely related aminoacids are indicated by colons (:) and distantly related aminoacids are indicated by dots (.). The NTDs of the different CER3 homologs are highlighted in yellow while the CTDs are highlighted in blue. The histidine-rich motifs are in red bold, the putative NADPH binding sites are in black bold and highlighted in green, and the putative catalytic cysteines are in red bold and highlighted in green. The alignment was performed byu using Clustal Omega (https://www.ebi.ac.uk/jdispatcher/msa/clustalo).

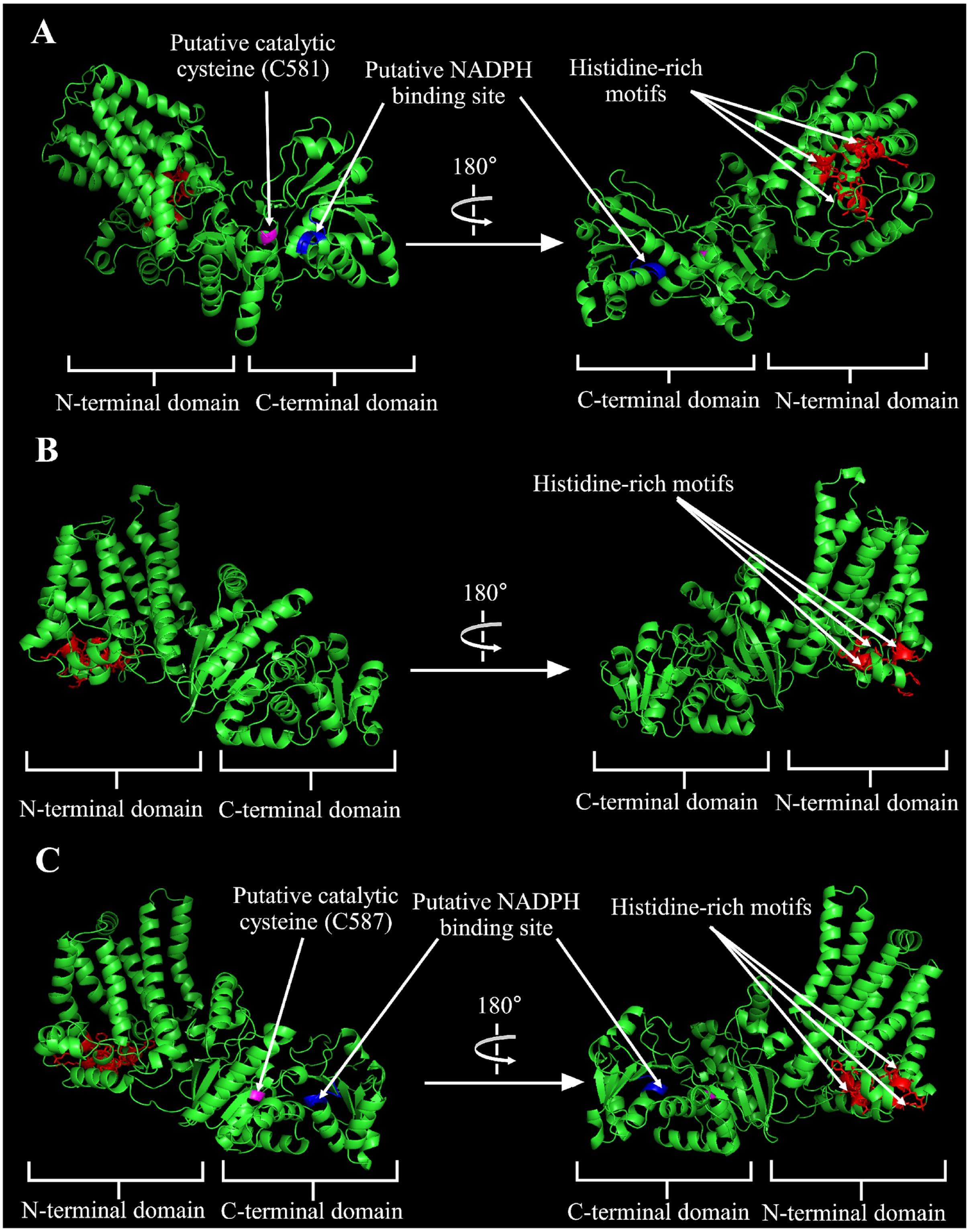


**Figure S4: Predicted structures of *A. thaliana* CER1, CER3, and *O. tauri* CER1/3 proteins. A)** Predicted 3D structure (AlphaFold 3) of *O. tauri* CER1/3. **B)** Predicted 3D structure (AlphaFold 3) of *A. thaliana* CER1. **C)** Predicted 3D structure (AlphaFold 3) of *A. thaliana* CER3. His rich motifs are shown in red, while the putative NADPH binding site and the putative catalytic cysteine are shown in dark blue and pink, respectively.

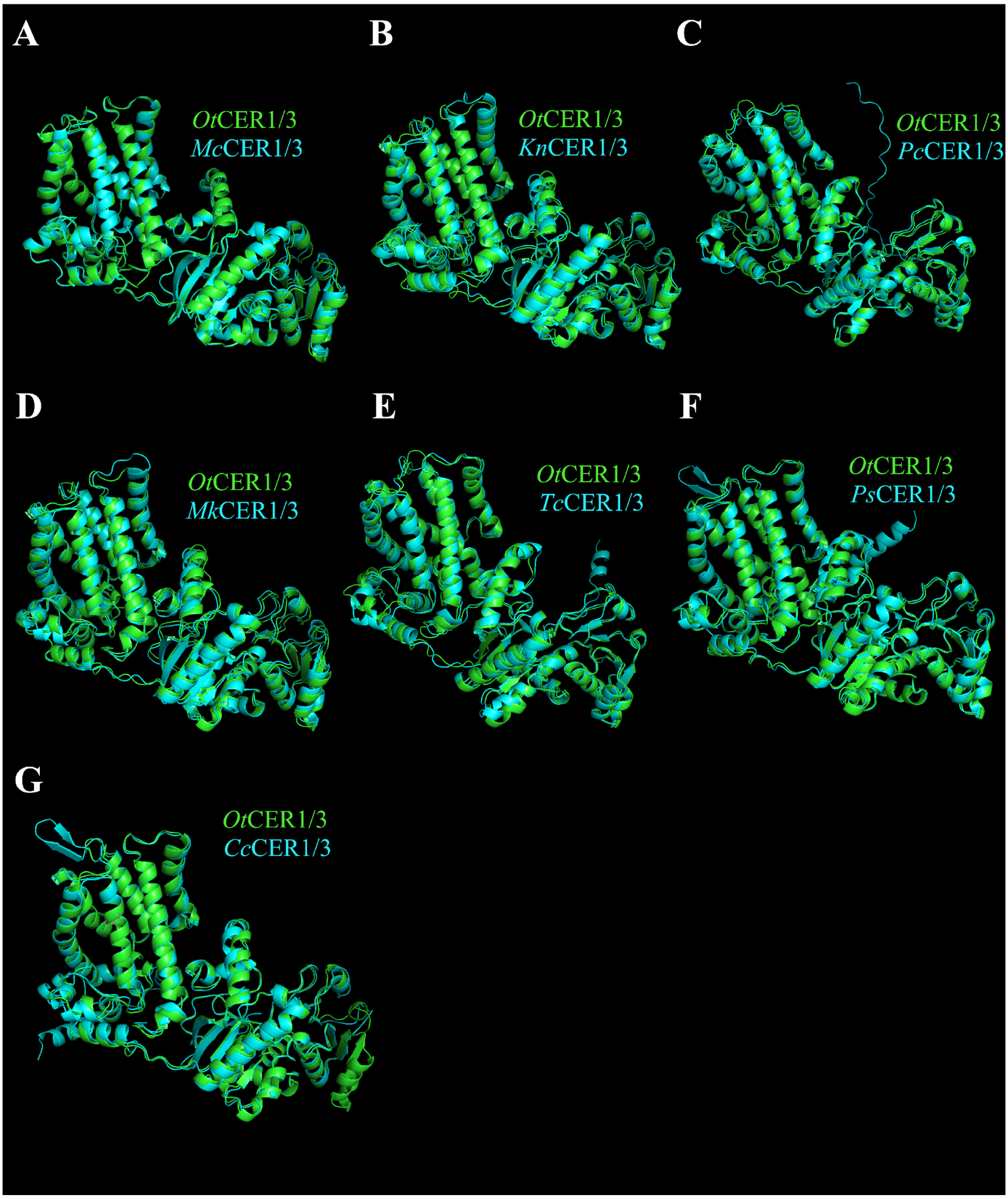


**Figure S5: Superposition of the predicted 3D structure of *Ot*CER1/3 (*O. tauri* CER1/3) (generated using AlphaFold 3) with the predicted 3D structures of CER1/3 homologs from other algal species.** *Mc*CER1/3 (*M. commoda* CER1/3), *Kn*CER1/3 (*K. nitens* CER1/3), *Pc*CER1/3 (*P. coloniale* CER1/3), *Mk*CER1/3 (*M. kramstae* CER1/3), *Tc*CER1/3 (*T. convolutae* CER1/3), *Ps*CER1/3 (*P. sulcata* CER1/3), *Cc*CER1/3 (*Cryptophyceae sp. CCMP2293* CER1/3);

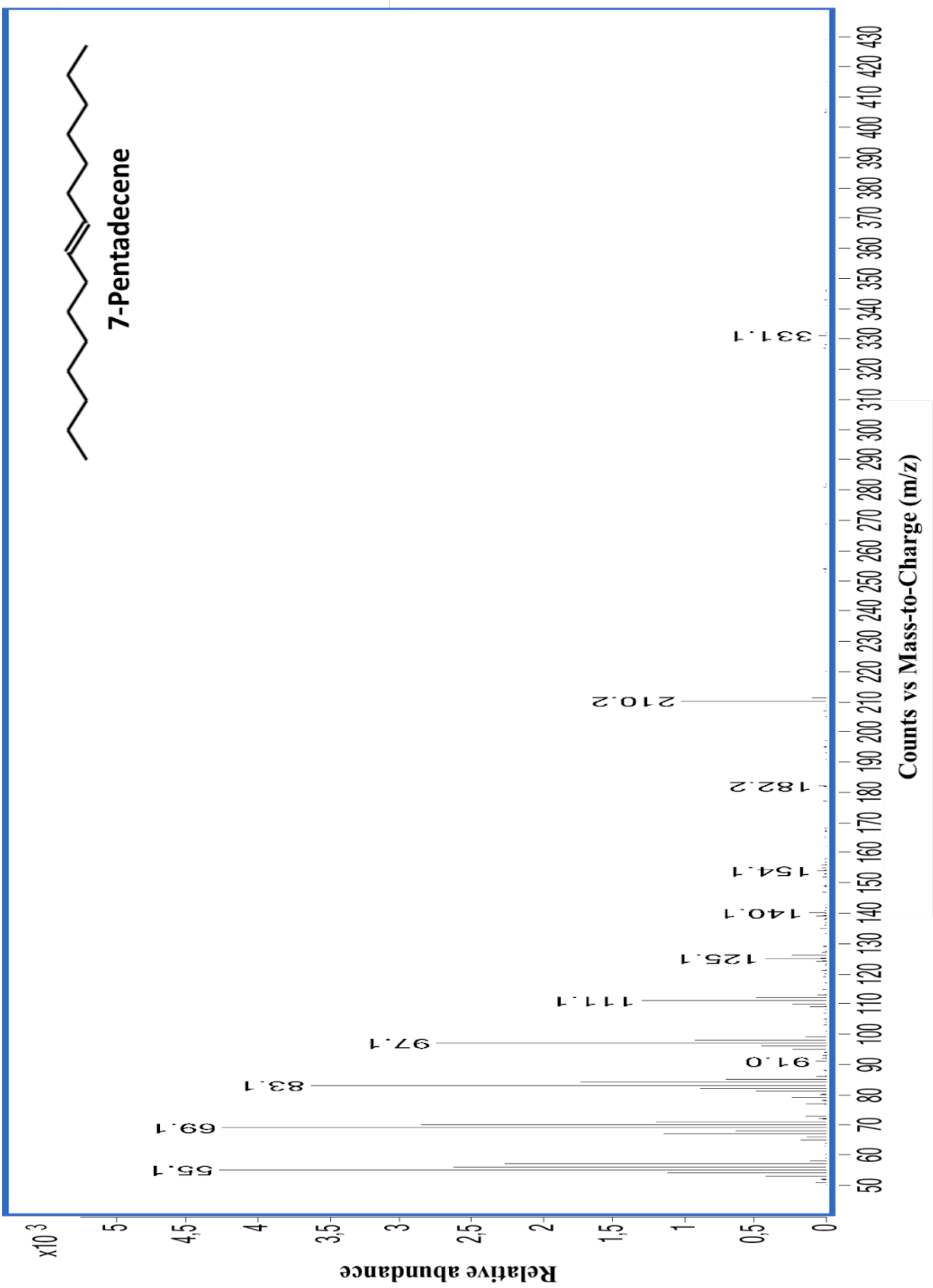


**Figure S6: Mass spectrum of the C15:1 HC.**

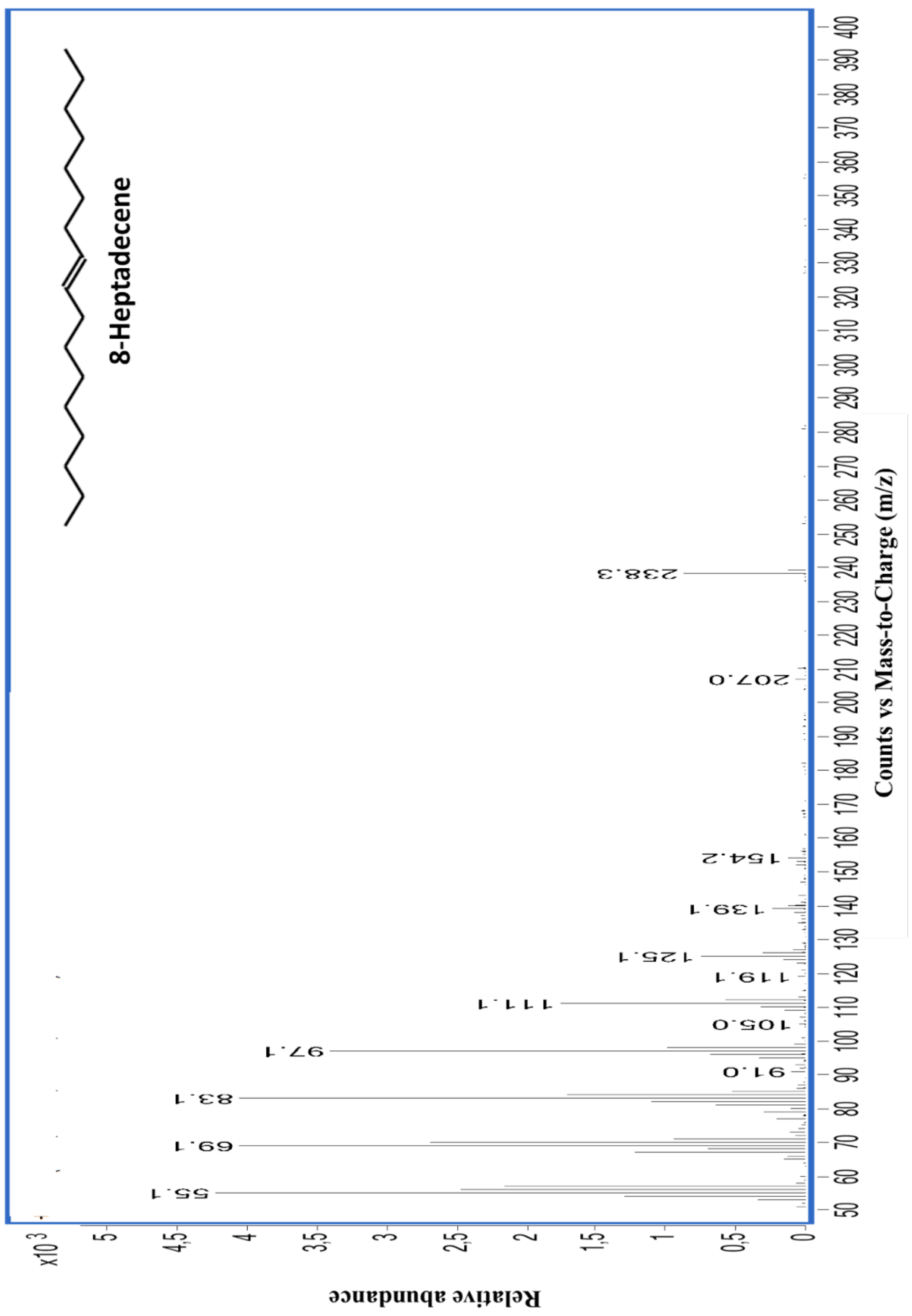


**Figure S7: Mass spectrum of the C17:1 HC.**

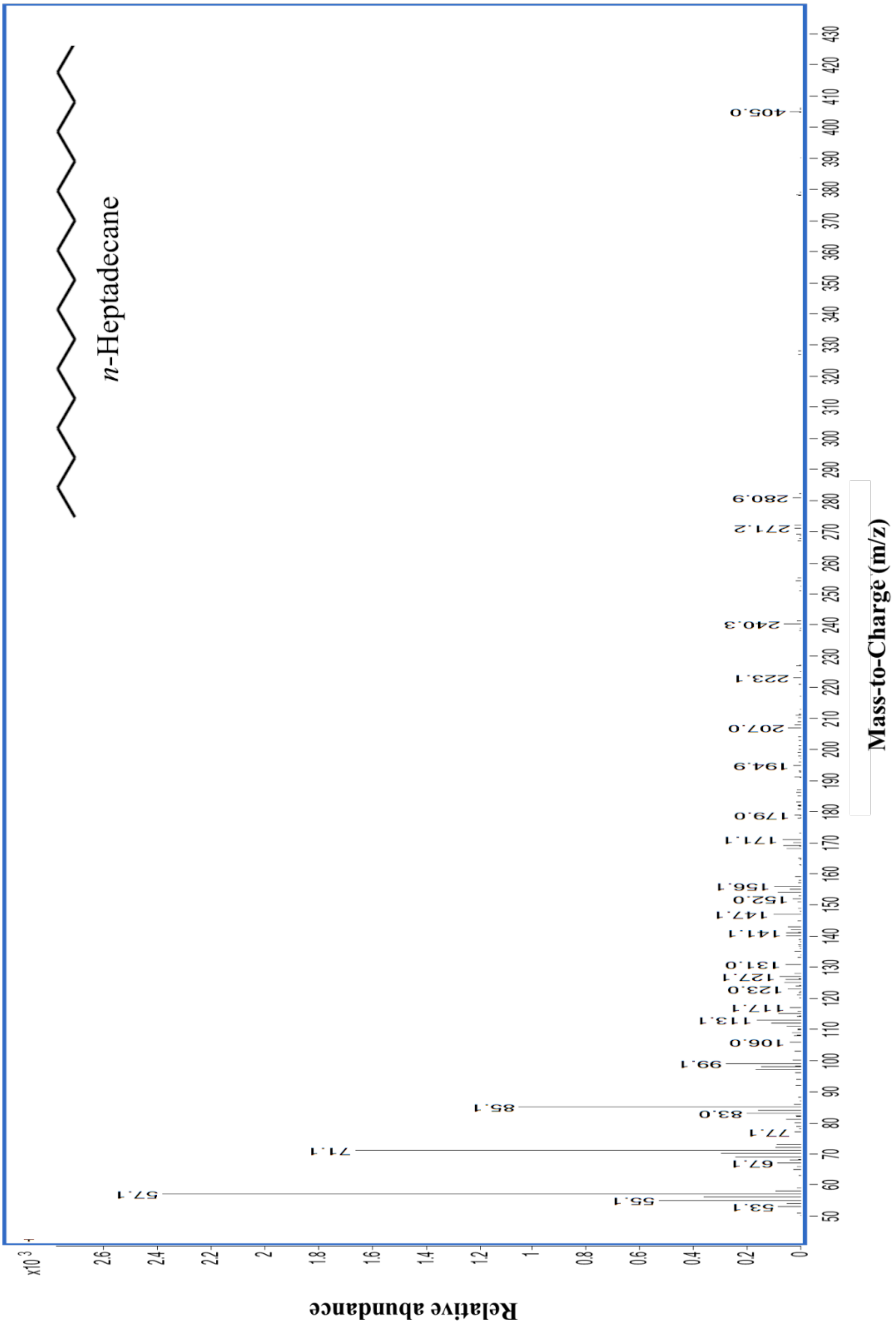


**Figure S8: Mass spectrum of the C17:0 HC.**

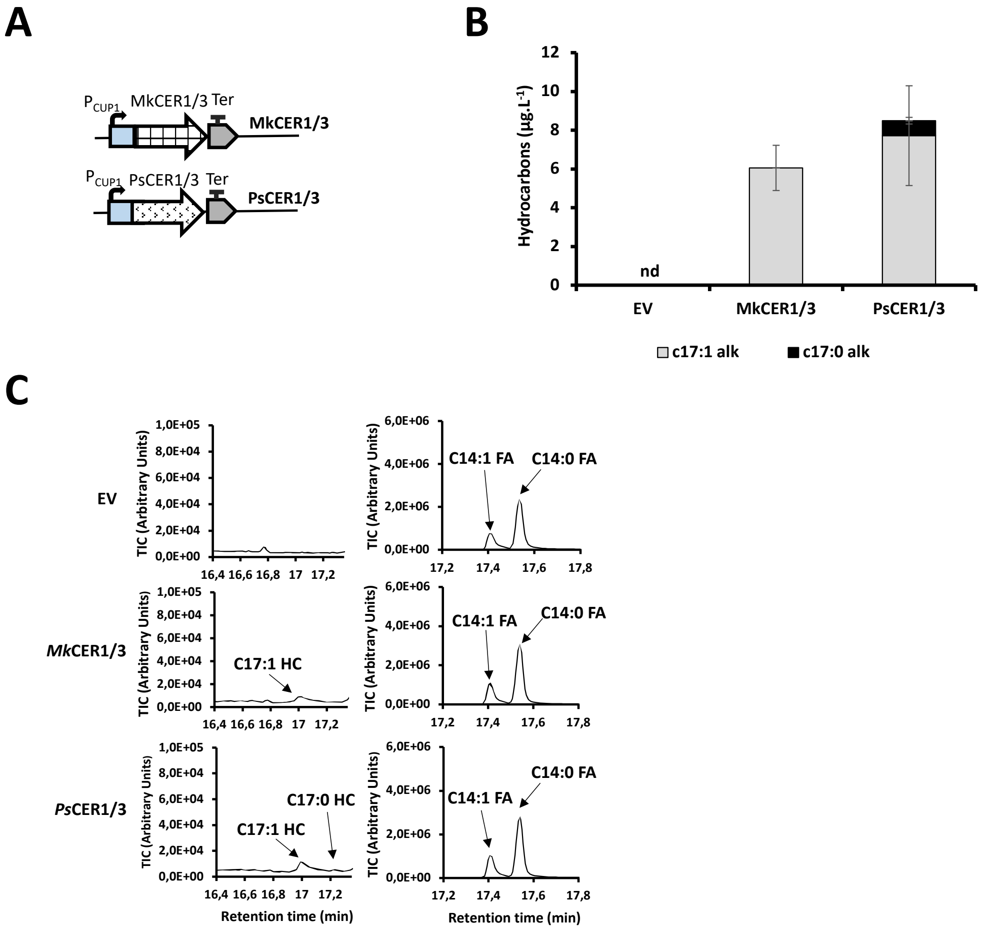


**Figure S9: Expression of *M. kramstrae* CER1/3 (*Mk*CER1/3) and *P. sulcata* CER1/3 (*Ps*CER1/3) in yeast. A)** Scheme of the genetic constructs showing that *Mk*CER1/3 and *Ps*CER1/3 were expressed in yeast strains under the control of a copper-inducible promoter ($P_{CUP1}$). **B)** HCs produced by yeast strains expressing *Mk*CER1/3 and *Ps*CER1/3. **C)** Chromatograms showing the different HCs, and the C14:1 and C14:0 fatty acids (used as loading control), present in yeast strains expressing *Mk*CER1/3 and *Ps*CER1/3. Error bars represent the standard error based on six biological replicates. 'HC' = hydrocarbon, 'FA' = fatty acid, 'nd' = non detected.

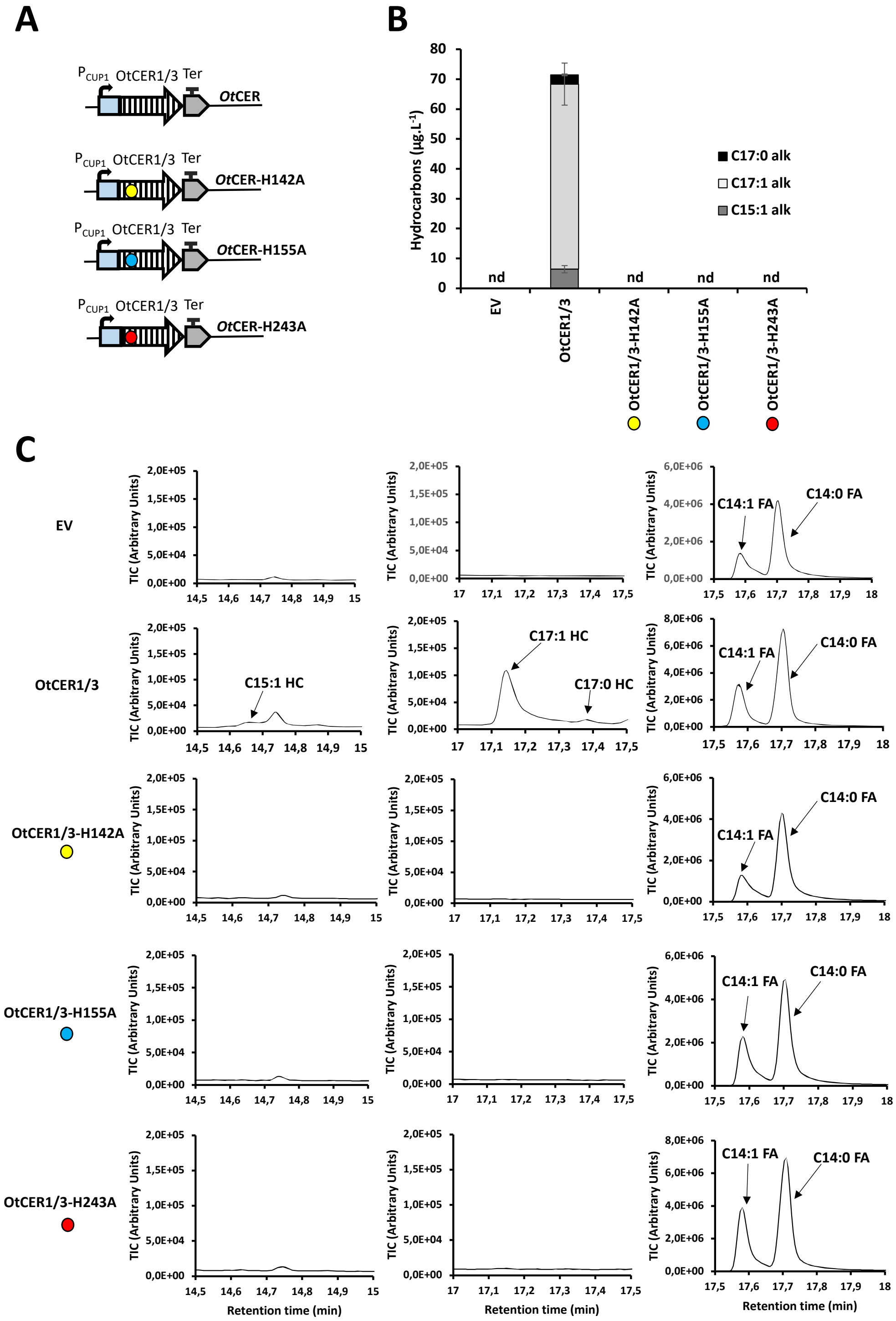


**Figure S10: Expression of different versions of OtCER1/3 in its NTD in yeast. A)** Scheme of the genetic constructs. OtCER1/3 mutated or not in its different histidine-rich motifs were expressed in yeast strains under the control of a copper inducible promoter ($P_{CUP1}$). **B)** HCs produced by the different yeast strains. **C)** Chromatograms showing the different HCs, and the C14:1 and C14:0 fatty acids (used as loading control), present in the different yeast strains. Error bars represent the standard error based on six biological replicates. 'nd' = non detected, 'HC' = hydrocarbon, 'FA' = fatty acid, yellow cercle = H142A mutation, blue cercle = H155A mutation, red cercle = H243A mutation.

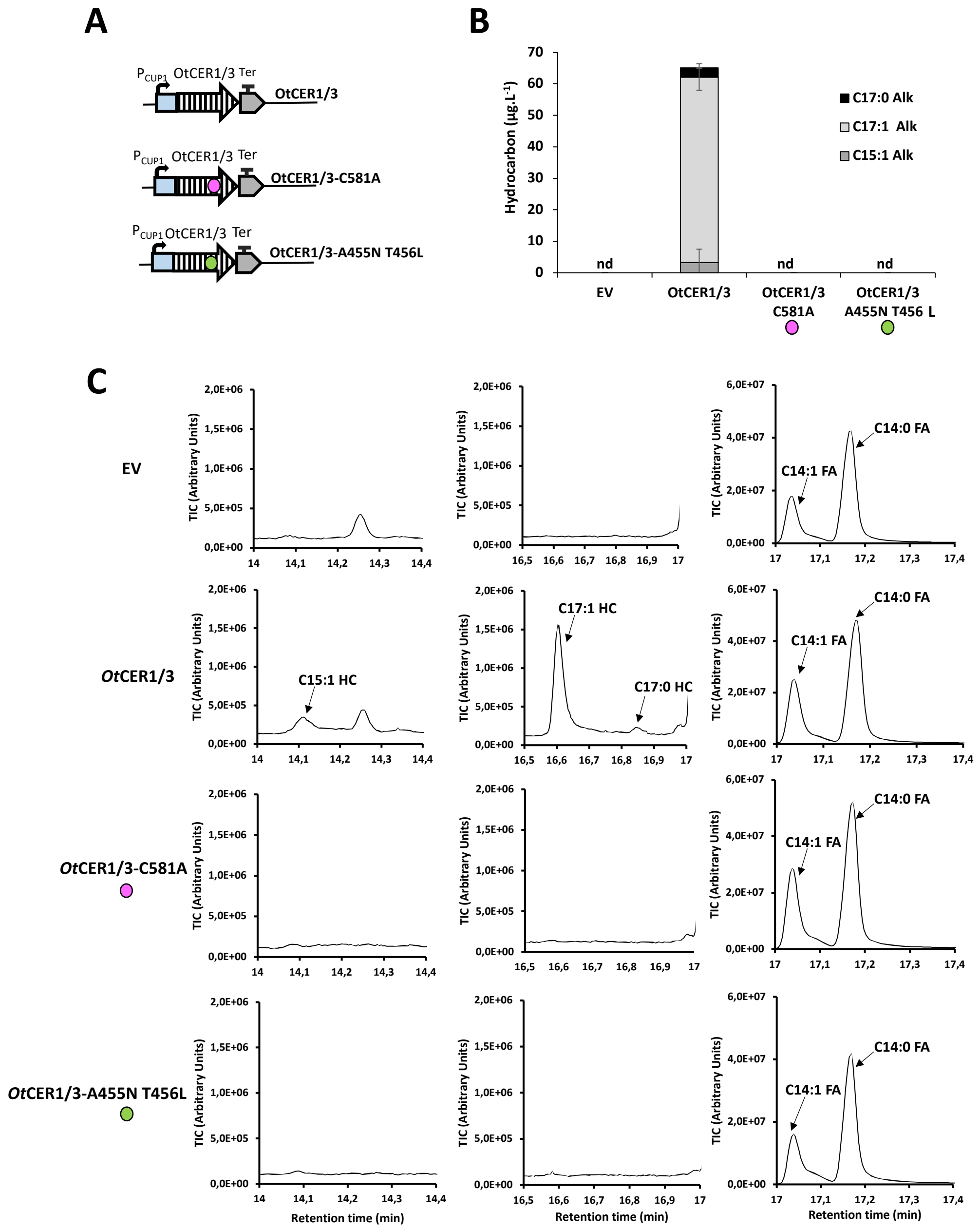


**Figure S11: Expression of different versions of OtCER1/3 in its CTD in yeast**. **A)** Scheme of the genetic constructs. OtCER1/3 mutated or not in its putative conserved catalytic cysteine, or in its putative conserved NADPH binding site were expressed in yeast strains under the control of a copper inducible promoter ($P_{CUP1}$). **B)** HCs produced by the different yeast strains. **C)** Chromatograms showing the different HCs, and the C14:1 and C14:0 fatty acids (used as loading control), present in the different yeast strains. Error bars represent the standard error based on six biological replicates. 'nd' = non detected, 'HC' = hydrocarbon, 'FA' = fatty acid, pink cercle = C581A mutation, green cercle = A455N and T456C mutations.

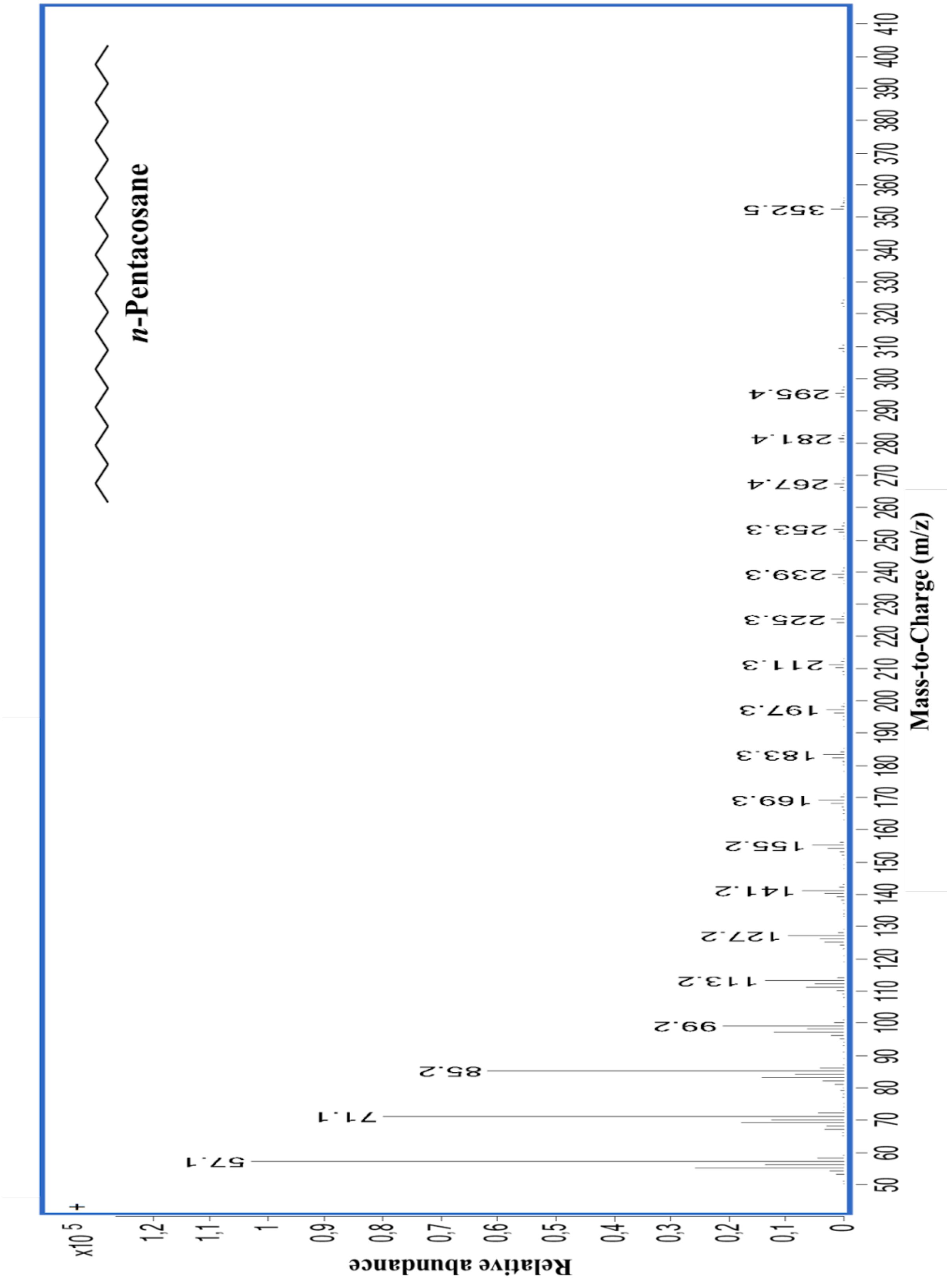


**Figure S12: Mass spectrum of the C25:0 HC.**

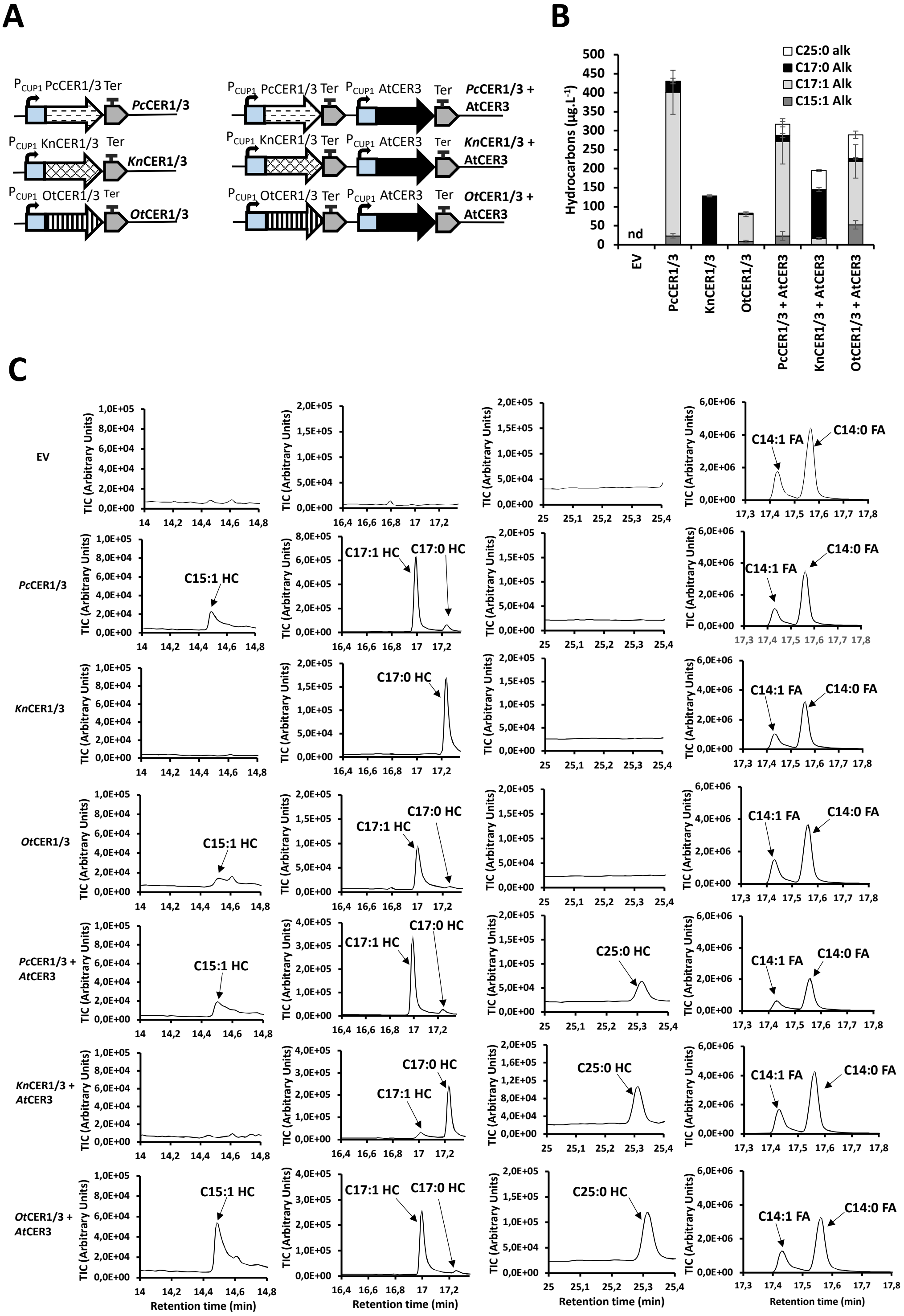


**Figure S13: Expression of *P. coloniale* CER1/3, *K. nitens* CER1/3 or *O. tauri* CER1/3 together or not with *A. thaliana* CER3 in yeast. A)** Scheme of the genetic constructs. PcCER1/3, KnCER1/3 or OtCER1/3 were expressed together or not with AtCER3 in yeast strains under the control of a copper inducible promoter ($P_{CUP1}$). **B)** HCs produced by the different yeast strains. **C)** Chromatograms showing the different HCs, and the C14:1 and C14:0 fatty acids (used as loading control), present in the different yeast strains. Error bars represent the standard error based on six biological replicates. 'nd' = non detected, 'HC' = hydrocarbon, 'FA' = fatty acid.

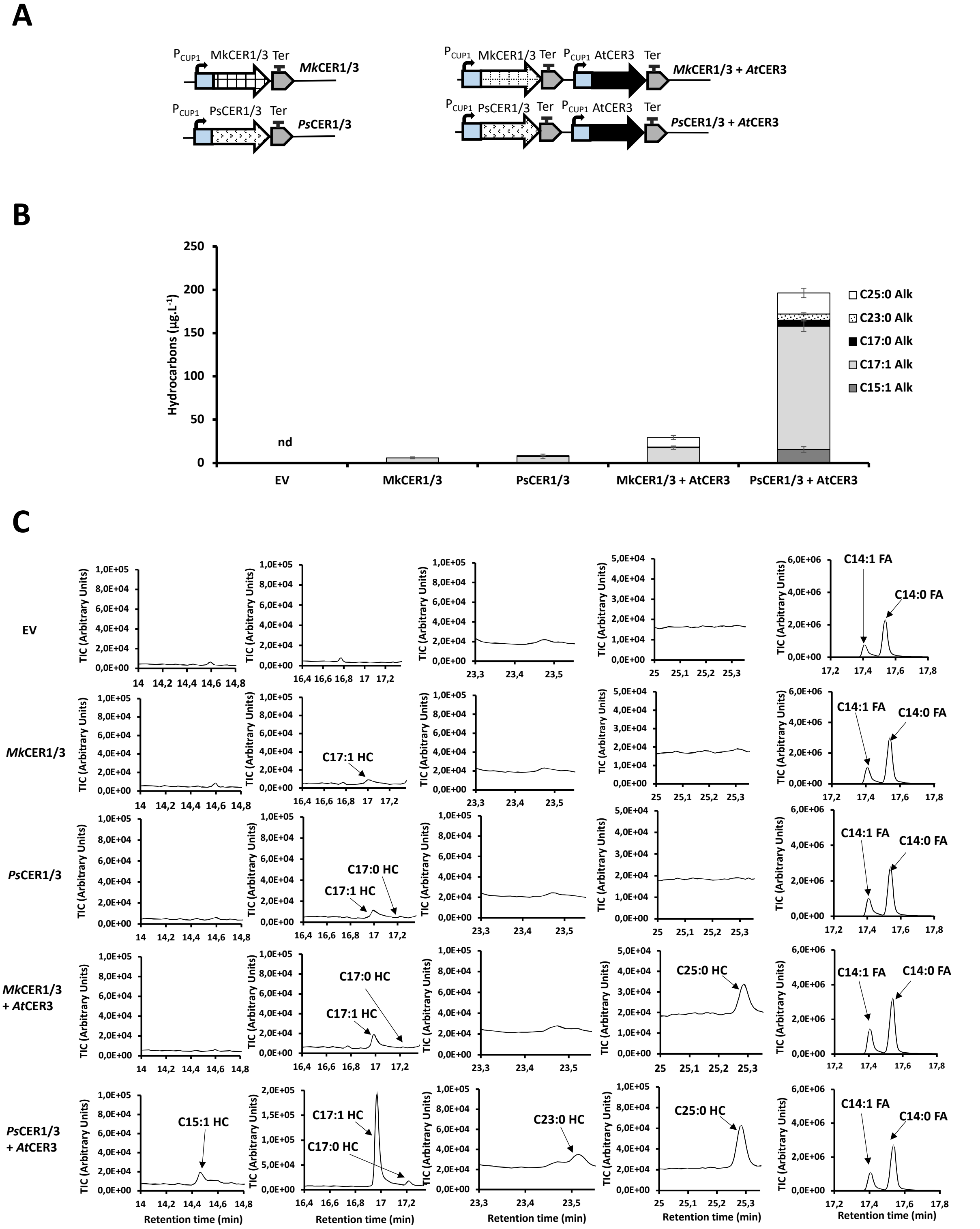


**Figure S14: Expression of *M. kramstae* CER1/3 or *P. sulcata* CER1/3 together or not with *A. thaliana* CER3 in yeast. A)** Scheme of the genetic constructs. MkCER1/3 or PsCER1/3 were expressed together or not with AtCER3 in yeast strains under the control of a copper inducible promoter ($P_{CUP1}$). **B)** HCs produced by the different yeast strains. **C)** Chromatograms showing the different HCs, and the C14:1 and C14:0 fatty acids (used as loading control), present in the different yeast strains. Error bars represent the standard error based on six biological replicates. 'nd' = non detected.

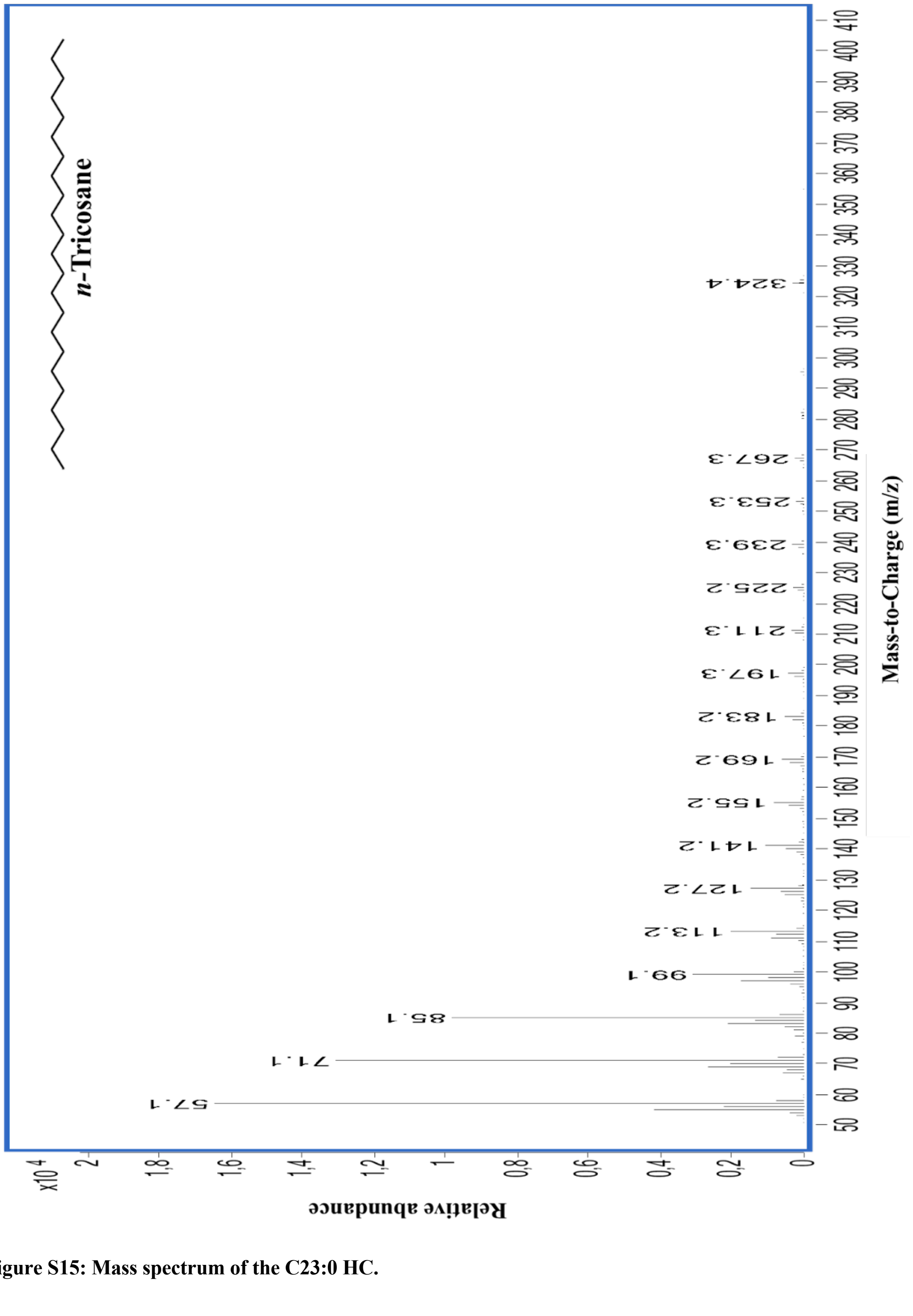


**Figure S15: Mass spectrum of the C23:0 HC.**

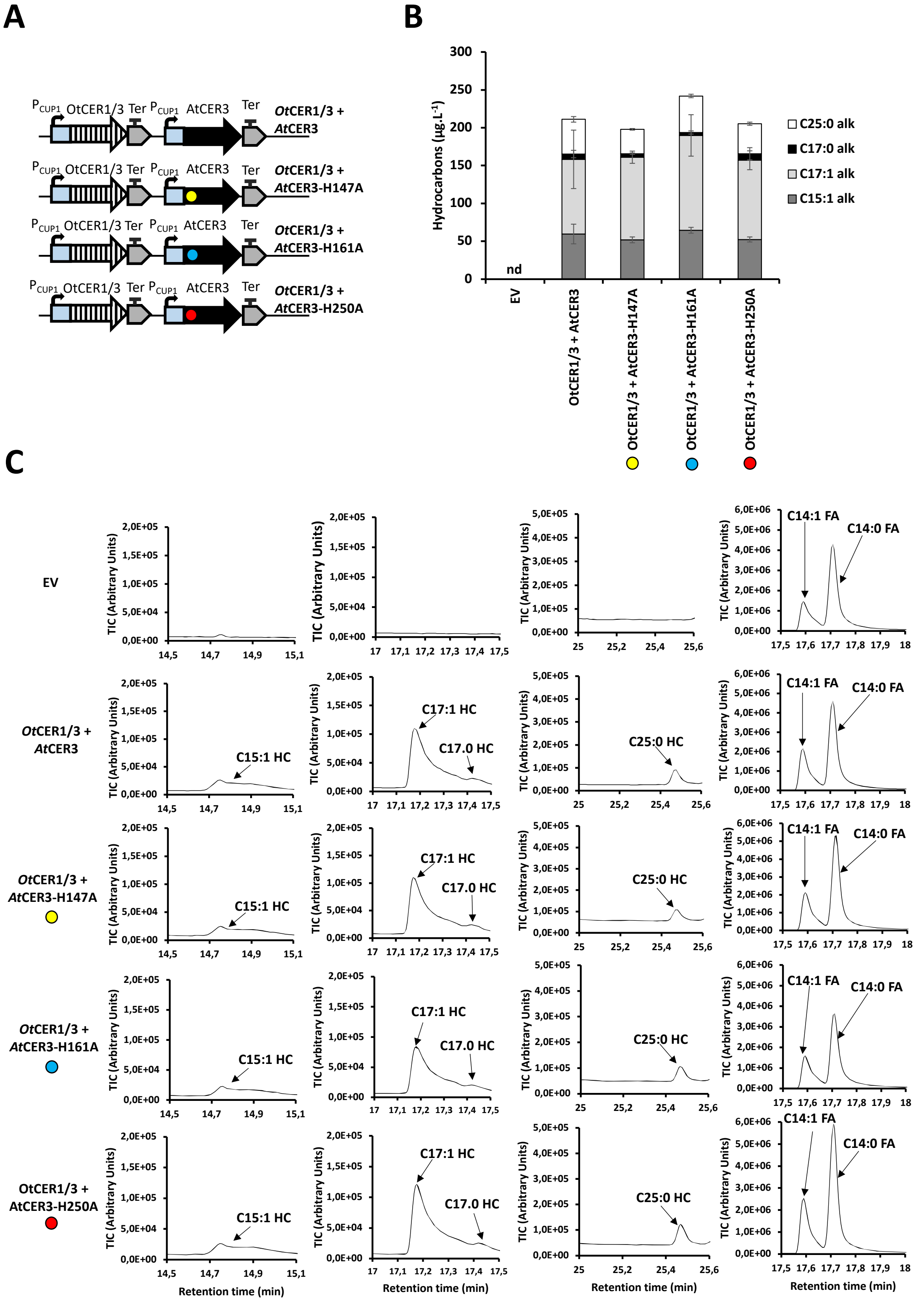


**Figure S16: Expression of *O. tauri* CER1/3 and *A. thaliana* CER3 mutated or not in its different histidine conserved domains in yeast**. **A)** Scheme of the genetic constructs. OtCER1/3 and AtCER3 mutated or not in its different His domains were co-expressed in yeast strains under the control of a copper inducible promoter ($P_{CUP1}$). **B)** HCs produced by the different yeast strains. **C)** Chromatograms showing the different HCs, and the C14:1 and C14:0 fatty acids (used as loading control), present in the different yeast strains. Error bars represent the standard error based on six biological replicates. 'nd' = non detected, 'HC' = hydrocarbon, 'FA' = fatty acid. Yellow cercle = H147A mutation in OtCER1/3, Blue cercle = H161A mutation in OtCER1/3, Red cercle = H250A mutation in OtCER1/3.

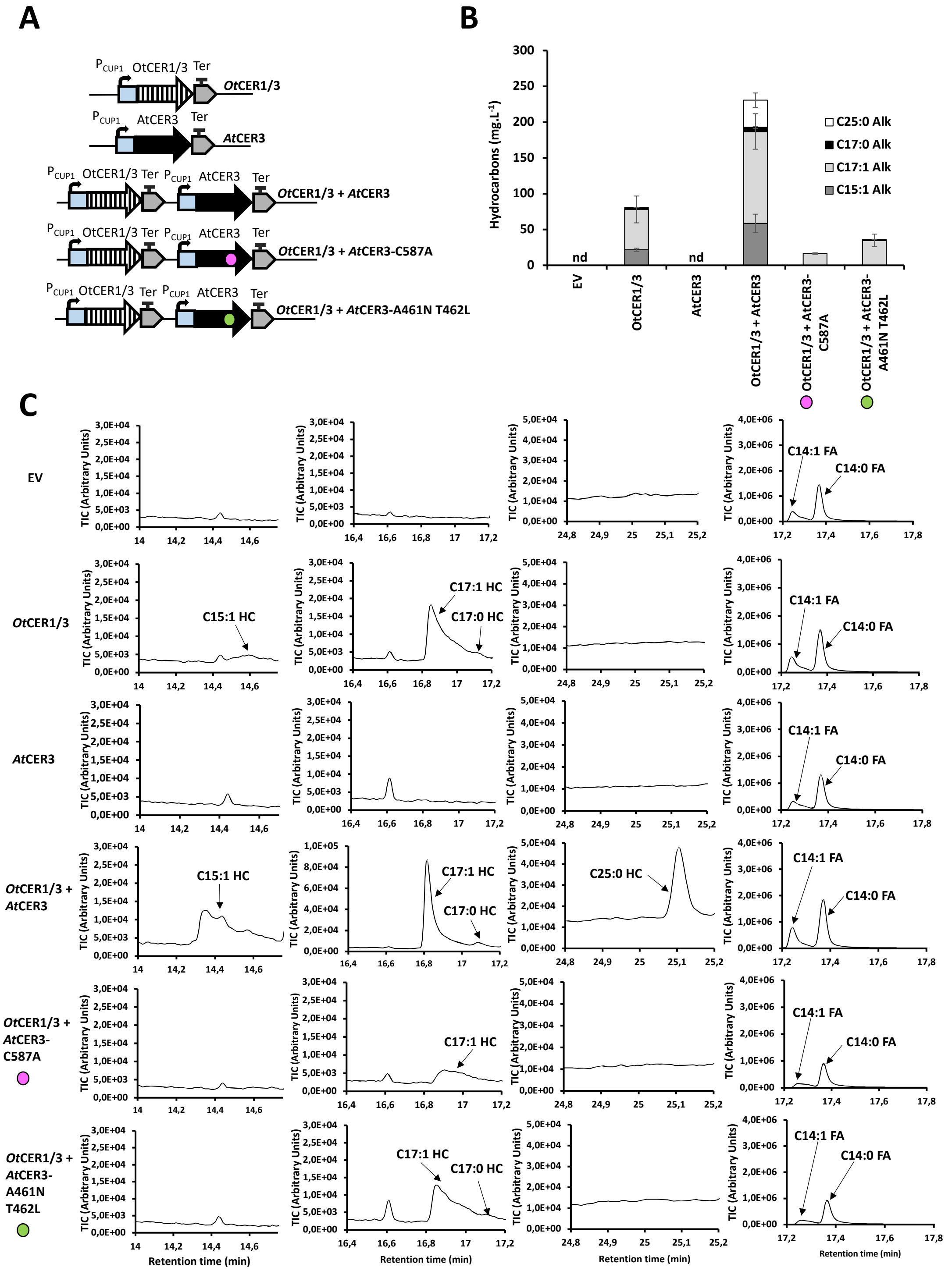


**Figure S17: Expression of *O. tauri* CER1/3 and *A. thaliana* CER3 mutated or not in the catalytic cysteine (C587A) or in the putative NADPH binding site (A461N T462C) in yeast. A)** Scheme of the genetic constructs. OtCER1/3 and AtCER3 mutated or not in its putative conserved catalytic cysteine or in its putative conserved NADPH binding site were expressed in yeast strains under the control of a copper inducible promoter ($P_{CUP1}$). **B)** Hydrocarbons produced by the different yeast strains. **C)** Chromatograms showing the different HCs, and the C14:1 and C14:0 fatty acids (used as loading control), present in the different yeast strains. Error bars represent the standard error based on six biological replicates. ‘nd’ = non detected, ‘HC’ = hydrocarbon, ‘FA’ = fatty acid, Pink cercle = C587A mutation in AtCER3, Green cercle = A461N T462C mutation in AtCER3.

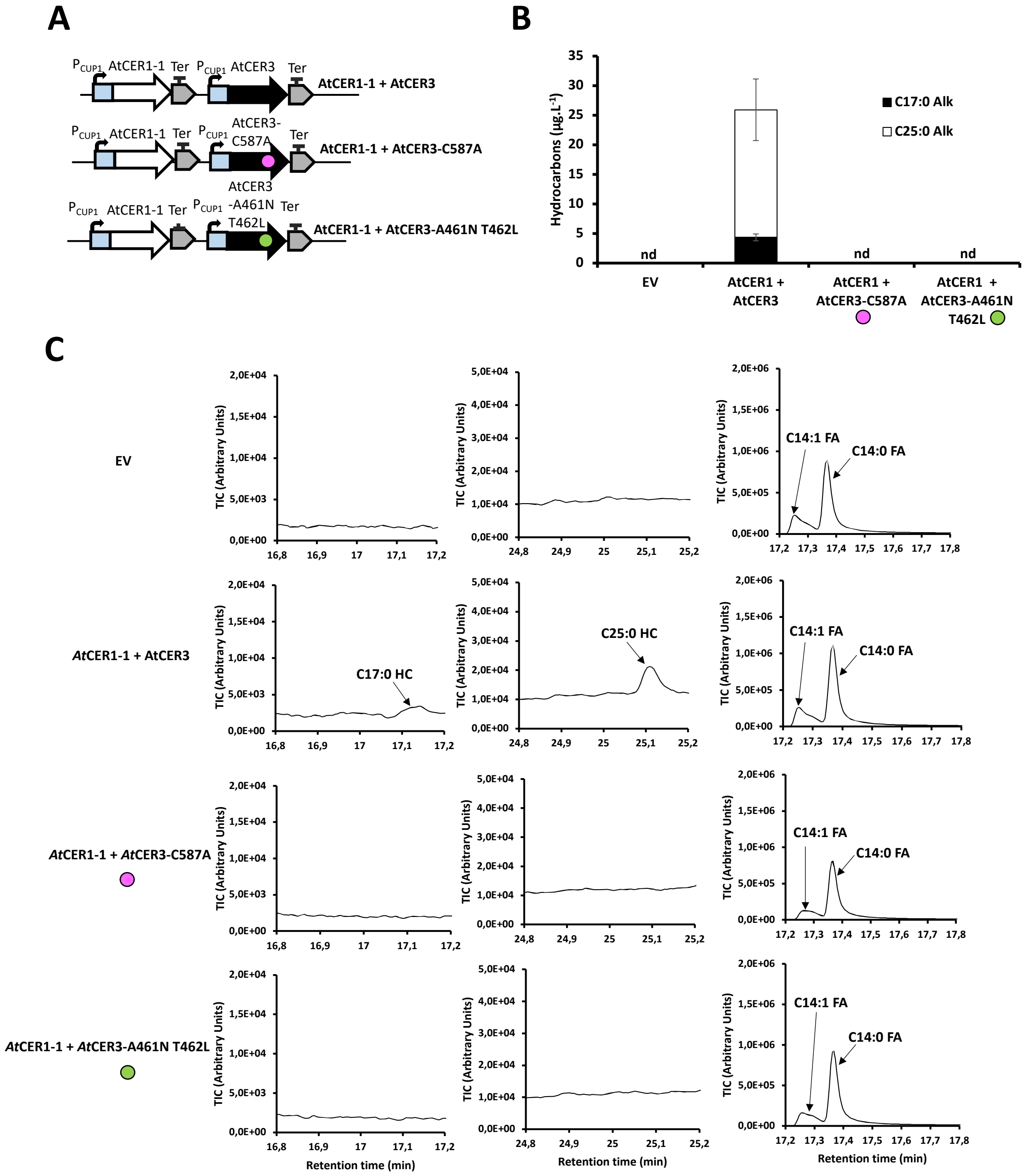


**Figure S18: Expression of *A. thaliana* CER1 and *A. thaliana* CER3 mutated or not in the catalytic cysteine (C587A) or in the putative NADPH binding site (A461N T462C) in yeast. A)** Scheme of the genetic constructs. AtCER1 and AtCER3 mutated or not in its putative conserved catalytic cysteine or in its putative conserved NADPH binding site were expressed in yeast strains under the control of a copper inducible promoter ($P_{CUP1}$). **B)** HCs produced by the different yeast strains. **C)** Chromatograms showing the different HCs, and the C14:1 and C14:0 fatty acids (used as loading control), present in the different yeast strains. Error bars represent the standard error based on six biological replicates. 'nd' = non detected, 'HC' = hydrocarbon, 'FA' = fatty acid, pink cercle = C587A mutation in *At*CER3, green cercle = A461N T462C mutation in AtCER3.

**Table S1: Protein sequences of all the genes analyzed or used in this work.** In addition to the protein sequences, information about the source organisms and the databases from which the sequences were obtained was also included. The protein sequence of *Tc*CER1/3 was provided by Xavier Bailly from the Roscoff Biological Station.

| Protein | Organism | Protein sequence | Database(s) |
|---|---|---|---|
| *Ot*CER1/3 | *Ostreococcus tauri (RCC4221)* | MATRPGVWYDFPWANVGALKYAVFAPFVLAVALGKDDADSFCWHLLAIAAARYVNAQLWISLSRVHAWTRNTR<br>IQAKGIDFKQVDREDHWDDYILLQTLVIAAVHWMPGLGFKDFPLYSGKSFAQLALLHAGPTEFIYYWLHRALHHH<br>KLYSAYHSHHHASFVTEPITGSVHPFMEHLMYTANFAIPLLGTWALGGGDIAMFYTYLIGFDILNAIGHCNFEFVPR<br>WFMRLPGMKYLIYTPSYHSLHHSRVHTNFCLFMPLYDYVYGTADVTSDELYEKAITGNAVPVKAPEVVFMAHGTE<br>LLSVFHLPFVLRSFSSRPFVSEWWLKPFWPLCVPFVLLLRVFGKSFVADRHRLKTLNCETWVTPAWGFQFFMKSEF<br>NHINKKIEEAILDADKSGVQVVGLGALNKNEALNGGGALFVNKHGKSLKTRVVHGNTLTAAAILQKIPNDCKEIFL<br>TGATSKLGRAIALYCAERGVRVVMYTTSEERFEMIRAEAPKKDQHLFVQSTSLTDGANIKDWVIGKHCSMKDQKS<br>APRGATFHQFVVPPIPESRKDCVYTDLPAFKLPRESKDFRSCEMTMPRGHVHACHAGALVHALEGWDHHEVGAID<br>HTRIDLTWEAALKHGFSLA | PhycoCosm (algae)<br><br>NCBI |
| *Mc*CER1/3 | *Micromonas comoda (RCC299)* | MALRPGPAYKFPWEDMGSFKYLLFVPFVATAALGLDDADNWAYHMLVIAAIRYVHAQFWISLSRIHAVTQHTKIQ<br>AKGIDYKQVDREDHWDDYIILQAIIMTLVHKMPYLGYNNFPQYNAMGMWQLLLLHAGPTEFIYYWLHRALHHHT<br>LYSWYHSHHHASFVTEPITGSVHPFMEHIMYTANFAIPLVGTWAFGGASIAMFYAYLIGFDLLNNIGHCNFEFMPQ<br>WFMNIPGVKYLIYTPTYHSLHHSKVHVNFCLFMPIYDYAYGTNDPSSDELYRKAINGEAAPNKAPDVVFVAHGTEL<br>LSLFHLPFALRSFSSKPFKSVWWLQPFLPLCIPFVALLRIFGKPFTADRHRLLHLNTATWVTPAWGFQFFIKSEFNHIN<br>RQIERAILEADATGTKVIGLGALNKNEALNGGGQLFVDKHPNLRVRVVHGNTLTAAAILKKIPADVKEIFLTGSTSK<br>LGRAIALYLSARGVRVVMYTTAKDRFEKIKAEAREEHRELLVQATTLEEGSGIKDWVVGKFCSARDQAKAPKHAT<br>FHQFVVPPLEESRRDCAYTDLPAFKLPKEAKDFRSCEMTMERGHVHACHAGALVHALEGWTYNEVGAIDHTKIDV<br>TWDAAVKHGFALA | PhycoCosm (algae)<br><br>NCBI |
| *Kn*CER1/3 | *Klebsormidium nitens (NIES-2285)* | MATKPGVLSEWPWQRMGNYKYVLYAPYVAAAAYEWGYQGKANEFIAYNAHLGVIAALRYVVMQIWTSIARFP<br>WLVGKYQISSKGIDFKQVDREDSWDDFILFNSLGLTLLGLVLPGWQTLPLVNGHGLVVMLLLHASVAEWLYYWF<br>HRALHHHWLYTRYHSHHHQSFVTEPITGSVHPFLEHVGYIAIFALPGVLTYVGGQGSIAMIYAYLLTFDFLNGLGHC<br>NFEFIPTWFFKAFPPLKYLIYTPSYHSLHHTRVHTNFSLFMPLYDYLYGTVDNDSDTLYEQTIKGVHPGPDAIFLAHG<br>MDVGSFVHGCFGIRGLASHPYYPSWVVYPLLPLATLLTIFCWLIDAKPWAVDRRSLRGNKIELWTVPRMGLQYFFP<br>SERPKINRLILQTLKKCDANGVKVFGLGALNKSEALNQGGEIYRKDIEGLRMRVVHGNTLTAASILHRLPQDCDEIF<br>LTGGTSKLGRALALYLVRKGVRVLLHTKSQERFQAIVDEAAPEHRKYLVWSPNLTDGSHCKNWVVGRWHSRKEQ<br>AFAPAGTHFHQFVVPLMTPARKDCTYSKQICMRMPKDTQGLGHCELNLQRNTVHACHAAVIVHLLEGWDFHELG<br>SINVDRIDENWEAALKHGFEPVS | PhycoCosm (algae) |
| *Pc*CER1/3 | *Prasinoderma coloniale (CCMP1413)* | MASKPGFMYEFPWEPMGNFKYLLLAPFAACAALGLDDADNWCWHMVTLVAVRYALAQLFITASRLHAISGKHRI<br>QKRGIDFEQMDREDNWDDFLILQALVMTAVHWILPGFKNFALYDGTGLAQCALLHAGPTEFVYYWFHRALHNHA<br>LYGAYHSHHHKSFVTEPTSGSCHPFMEHLGYTANFAIPMLGTWFLGGASMAMFYAYTIAFDTLNMIGHCNWEFVP<br>VRLYQAVPLLRYLVYTPSYHSLHHSRVHTNFCLFMPLYDYLGGTLEMERDGRTWTSDKLQAEAMARTYEVPDAV<br>FLAHGTDLLSTFHLPMFFRGFAKNPYSESWLLYPLWPLAWLAMAVLWVVGEAFPSHVERVNGHPVQTWVTPRYG<br>FQYFLKFERRRINGLIERAIRKADAQGVKVFGLGALNKAEHVNGGGVIFPEAIKDLRTRVVHGNTLTAACLVHELP<br>ADVREVFLTGATSKLGRAAALWLCRRGVRVRMLTSSVERFEAIRAEAAPEHRELLVRVEHHRDGGDCKTWIVGK<br>MCGAKDQACAPPGTHFHQVVVPPIAKHRSDCTYGELAAMDLPDTATDVKACELFMPRRRVFACHAGALTHMLEG<br>WDHHEVGAIDIDRIDVTWEAALRHGFRPAVAPAGCAVGCAQPAAVAARASDAAAIPAAVAAA | PhycoCosm (algae) |
| *Mk*CER1/3 | *Mesotaenium kramstae (Lemmermann NIES-657)* | MATTPGAFYEWPWQSLGNFKYAIYAPFVAKAVHANLLGGEDPDNWCLHMVLSSALRMFHGQLWMSASRVHYW<br>TQKHKIQLKGIKFEQVDREERWDDYIILHVLVATFVHAVLPGFQNFPLTDGWGLLILLLLHMGPAEWVYYWGHRL<br>LHHHYLYTRYHSHHHASFITEPVSGSVHPFAEHVMYTAMFAIPFLGTWALGGASIAMFYAYWLGFDFLNAIGHCN<br>WEFMPSWLFQAFPPLKYLIYTPSYHSLHHSQVHTNFALFMPLYDYLGGTVDGMSDLLHSSVRKGRANQPDFVFLA<br>HGTELLSLFHLPFGIPSFASRPYRPSWVMYLLWPLTLPIMAIIWLIGKVFVSDSYTLESLRMQTWVVPRLGFQYFLSG<br>EKKRINKHIEQAILDADELGVKVFTLGALNKNEALNGGGTLFVEKHKDLRVRVVHGNTLTAAVILDKIPKDAREIFL<br>TGATSKLGRAIALYLCQRGVRVLMLTAARDRFEAIQREAPEECRHLLVQAESYAEGANCKSWVVGKWMTARQQA<br>HAPPGTHFHQFVVPPTPEVRKDCTYGKLAAMRLPNTVQNMRTCEMTMDRGCVHACHAGGLVHALEGWDHHEV<br>GAIDPSKIDVTWEAALRHGFQPV | PhycoCosm (algae) |
| *Tc*CER1/3 | *Tetraselmis convolutae* | MATKPGPLYEWPWAAAGDLKYLLFTPFIAAVALGKDDEDNWASHMLAIAALRYVQNYLWIFVSRCDWLSGRNKI<br>QHKPVTFKQVDRENHWDDYIILQAFVMTAVHWVLPGFASFPVNNWNGLWQTLMLHVGPTEFIYYWFHRALHHH<br>SLYSQYHSHHHASFVTEAITGSVHPFLEHVGYTANFAIPLLGTWLMGGASWTMFYAYLAAFDFLNAMGHCNFEFV<br>PTGILKACPWIKYLIYTPTFHALHHAHVRTNFCLFMPIYDYIYGTWDPDSDPWQETAWKGRQTEEGVVFLAHGTE<br>MLSLFHLPFMTRAFASKPYQTTWAMYLLWPLTIPFLLVFWAFGSVFVTDRNLLGKLKMQTWCTPRFGFQYFLKFD<br>KPRINRLIKDAVIKADKLGLQVIGLGALNKAEPLNGGGKAILESALAERDLKVRVVHGNTLTAGCVIKELPAGVKEI<br>FLTGSTSKLGRAIALYMSAKGTRVIMFTKSAERFAAVQEEAKLEHRHLLVHSTDYTDGSKCKTWVVGTLLGAKQQ<br>AFAPKGAFFHQFVVPPISETRSDCSYGKLAAMRLPPKTLMRACEMTMERNCVHACHAGALVHALEGWEHHEVGA<br>IDVSRIDLTWKAAMSHGFKPVFESQAAPVKEAAAAAA | |
| *Bp*CER1/3 | *Bathycoccus prasinos* | MASRPGFAYDFPWSKIGKMKYAIYLPMLYRGIVSPENDDSDQWHFHMTMIVLLRYVMAQFFISLSRIHAITEKTRIQ<br>SKGIDFKQVDREDHWDDYILLQYIVMSMVHFCPGLGFKNFPLFEKKGMWQLLLLHVGPTEYVYYWLHRLLHHHT<br>LYSAYHSHHHASFVTEPITGSVHPFMEHIMYTANFAIPLLGTWMCNGASMAMFYVYLMGFDLLNAIGHCNFEFVP<br>KFFAKFPGVKYLLYTPSYHSLHHSRVHTNFCLFMPIYDYAYGTMDKSSEELYDKAIEGKASPKTTPDVVFMAHGTE<br>LLSMFHLPFAFRSFSSRPFTTDSWMLKMLWPLTLPAVAALRFLPGVKAFVSDKHRLKNMNIETWVTPAWGFQFFIR<br>SEFKHINAKIERAILDADERGVRVLGLGALNKNEALNGGGAFFVQKHEKNLKNTKVVHGNTLTAAAIIDKIPENVK<br>EIFLTGATSKLGRAIALYMATKKNCRVLMCTTSEERFEKIKMECPEKFRHLLFRVNNANEKVEITQESTSNVLKKSG<br>SFLLSRLGSLKNNNNNNNREVETEEKNDTKTNYSSGRTCRNWVVGRHCDKNEQSLAPSKTTFHQFVVPPIPETRSD<br>CAYTDLPAFRLPEKEAKDFKTCEMTMERGCVHACHAGALIHALEGWQHHEVGA<br>IDPEKIDVTWKASKKHGFACL | PhycoCosm (algae) |

| | | | |
|---|---|---|---|
| *Cs*CER1/3 | *Crustomastix stigmatica* | MATAPGPLYAFPWESWGECGKYALLLPFAAAVVLGRDDGDNWAWHMCVIAVARYVHAQLWISISRLHRLTSKT<br>RIQARSVGFKQVDREDHWDDYIILQALIMTLVHWCPGLGFKGFPVWGSWVGMAQLLLLHAGPTEFIYYWLHRAL<br>HHHSLYSRYHSHHHASFVTEPITGSVHPFMEHLMYTANFAVPLLGTWACGGGSVAMFYAYLLGFDFMNAVGHCN<br>WEFVPRQIFRALPFLKYMVYTPSFHSLHHSKVHVNFCLFMPIYDYAYGTLDESSWDLHERASLGVAVPDVAPDTVF<br>LAHGTELLSMFHLPFALRSFSSVPFVPRWFLWPMYPVAALTVVLLRLFGTVFTQDKFSIGHLKGETWVTPAFAFQFF<br>LKREHSFINRQIAGAIRQADARGVRVFGLGALNKAEFVNGGGVLFTKHLPDLRIRVVHGNTLTAAAILQKLPEGVR<br>EVFLTGSTSKLGRAIALYLAERDVRVRMYTTSEERYRAIADEVEPKRRHLLSRHDRLADGKDVEHWVVGKWLTA<br>KEEQVAAPGTTFHQFVVPPLEGTRKDCVFTQLPAFRLPQKKTRGVKACEMTMPRGCVHACHAGALVHGLEGWDH<br>NEVGRIDHTKIDTCWEAAMRHGFELV | |
| *Ts*CER1/3 | *Tetraselmis striata* | MSGLRNDPRWCTPGIAATEAEHGKLAADRASSAPVQGVHRYYPFVVEDRGRLGKSALTVVYIFAALIANSAPDRP<br>HSAVPAMATKPGILHDWPWAPLGDFKYLLFVPFVATVALGCDDKDNWAMHMLLISALRYIMNWFWIFVSRNDAL<br>SGKNRIQHKPVTFQQVDRENHWDDYIILQAIVMTAVHNVLPGFQSFPMNNWKGLGQMVAWHVGPTEFVYYWLH<br>RALHHHSLYSRYHSHHHASFVTEAITGSVHPFLEHVAYTANFAIPLLGAWAMGGASWAMFYSYLLGFDFLNAWG<br>HCNFEFFPLWVLQTFPFIKYLIYTPTFHALHHSHVRTNFCLFMPIYDYLYGTWDADSDKLHAQSRYGRKQEEGVVF<br>LAHGTELLSFFHLPFITRAFASKPFRASWAMYLLWPLTLPAMLVVWAMGGVFCTDRNQLGKLQMQTWSMPRFGF<br>QYFLSLTSPMFKFEKARINKLIKSAIVKADDKGLQVIGLGALNKAEPLNGGGKQLLEEILAERELKVRVVHGNTLTA<br>GAVLKELPGNVEEIFLTGSTSKLGRAIALYLSARGTRVIMYTKSAERFAAVRDEATPAAQKFLVHATSLAQGSGCK<br>TWVVGTALRAKEQAAAPKGTFFHQFVVPPIEEARKDCSYGKLAAMRLPPKALMRACEMTMERGCVHACHAGAL<br>VHALEGWEHHEVGAIDVSRIDQTWEAALRHGFKPVFTQQ | PhycoCosm (algae) |
| *Smu*CER1/3 | *Spirogloea muscicola* | MATKPGLLSEWPWQPLGQFKLPTTGYSCLLSQPGSLVRCLSGVQYLLFAPFAAKAIQANFYGGHDKDNWCMHMLI<br>ASALRYLHGQIWMSASRFHYLVEKYQIQTKGIKFEQVDRESSWDDYIILHVLVATFVHSFLPGFQNFPLYNGWGLVI<br>LLLLHAGPAEFIYYWGHRLLHHHYLYTRYHSHHHASFVTEPVTGSVHPFAEHVMYTAMFAVPFLGTWALGGASLS<br>MFYAYWLGFDLLNCIGHCNWEFIPTWVFKAFPPLKYLVYTPSYHSLHHTQVHTNFALFMPLYDYLGGTVDGMS<br>DLLHSSIRKGRTDRPDFIFLAHGTELLSAFHVPFGLPQFAAHPYAPSWLMYLLWPLTLPLLALVWLFGQTFVADKY<br>KLRDLRMQTWVIPRFGFQYFLPFEKKHINNHIEKAILDADRIGAKVFTLGALNKNEGLNGGGTLFVERHKHLRVRV<br>VHGNTLTAAVILDKIPKDVEEVFLTGATSKLGRAISLYLCRRGVRVLMLTSALDRFCAIQQAAPAECRHLLVQATSY<br>ADGKDCKTWILGKWLTSKDQACASAGSHFHQFVVPPVVETRADCTYGKLAAMRLPPEVQGLRTCEMTMERG<br>AVHACHAGGLVHALEGWPHHEVGAIDPDRIDITWEAALKHGFRPV | PhycoCosm (algae) |
| *Ps*CER1/3 | *Tetraselmis convolutae* | MWSGIARAGAFAQWFNDSSKKQKVRGALKPGILYDSPWESLGSYKYSLFLPFLYAVVSGNDDEDNWCYHMLMIA<br>ALRYIQAWIWNFLSRNHHVSGKNRIQAKGVDFVQVDREDNWDDYILLQIYVASAVHLLPFLYCWDGECYAYKNF<br>PLHNNKGLIKLLLIHMGPTEFIYYWLHRALHTHQLYANYHSHHHASFVPEVVTGSVHPFMEHLMYTANFAIPLVGT<br>WLWGGASIAMFYTYLLGFDLLNMIGHCNFEFFPVWPFKYIPGLKYLIYTPSFHSLHHSRVHTNFALFMPVYDWIGG<br>TLDKKSWSLFESAAQGEAVPQKAPDAVFLGHGTEMLSVFHLPWMSRSFSAQPYEAQWWMYILWPIALVPLAVIRL<br>VGKALIQDKYRLGDFNCESWFTPVVALEFMFKSQWKSINSFIENAIVEADKSGVRVFGLGALNKNEALNGGGLQFI<br>QKHPELKMCVVHGNTLTAAALLQRINQDVKRVFVLGSTSKLGRAISLYLAKKGVKVMMMTNSRERYENTIEDCP<br>KEYRGNLIHSTDMKDGANCKSWVVGRFLSKREQALAPSGTTFHQFVVPELERVRTDCSYTALPAFTLPKMAQGLR<br>SCEMTMGRRNVHACHAGALVHLLEGWKHHEVGAIDPSRIDATWEAAMRHGFKLVEEEDA | PhycoCosm (algae) |
| *Cc*CER1/*3* | *Cryptophyceae sp. CCMP2293* | MWPGAARLKAMGSWMGGTVTAPNEKEKDSLRHKGATQPGLWYDFPWHAMGDWKYLLYLPFVVVLALGLDDA<br>DRWCFHMMAVCVLRFTQAWVWNLLSRNHFISGKTRIQGKGVEFKQIDRESNWDDYIILQLYVATAVHCLPFLYCS<br>NGVCHAYRNFPLYDGKGLVKMLLIHAGPTEFIYYWLHRALHLHSLYARYHSHHHASFVPEAVTGSVHPFMEHLM<br>YTINFAIPLLGTWLAGGASVAMIYTYLLGFDLMNIIGHCNFEIFPVWPFKVFPFLKYIIYTPSFHSLHHSRVVTNFCLF<br>MPMYDWMYGTLDPKSWDLFDKAAAGNAVQHAPPETVFLAHGTELLSVFHLPFMSREFSSKPFQPSMWMYILWPI<br>AVPMLLLARLFGTVFVADKHRLGAMRIETWVTPAWAIDFFFKSQWGRINRAIDAAITDADKAGVKVFGLGALNKN<br>EALNGGGALFVKNHPDLTMRVVHGNTLTAAAIIKTIPADVKKAFVLGATSKLGRGISLYLAKRGVTVVMLTQSEER<br>VACWVVGRFLSKEEQHKAPKGAAFHQFVVPPLDELRKDCVYTKLPAFTLPENAAGFKSCEMTMQRRDVHACHAG<br>ALVHALEKWTHHEVGAIDEEKIDRTWDAAMKHGFKMVC | PhycoCosm (algae) |
| *At*CER1 | *Arabidopsis thaliana* | MATKPGVLTDWPWTPLGSFKYIVIAPWAVHSTYRFVTDDPEKRDLGYFLVFPFLLFRILHNQVWISLSRYYTSSGKR<br>RIVDKGIDFNQVDRETNWDDQILFNGVLFYIGINLLPEAKQLPWWRTDGVLMAALIHTGPVEFLYYWLHKALHHH<br>FLYSRYHSHHHSSIVTEPITSVIHPFAEHIAYFILFAIPLLTTLLTKTASIISFAGYIIYIDFMNNMGHCNFELIPKRLFHLF<br>PPLKFLCYTPSYHSLHHTQFRTNYSLFMPLYDYIYGTMDESTDTLYEKTLERGDDIVDVVHLTHLTTPESIYHLRIGL<br>ASFASYPFAYRWFMRLLWPFTSLSMIFTLFYARLFVAERNSFNKLNLQSWVIPRYNLQYLLKWRKEAINNMIEKAIL<br>EADKKGVKVLSLGLMNQGEELNRNGEVYIHNHPDMKVRLVDGSRLAAAVVINSVPKATTSVVMTGNLTKVAYTI<br>ASALCQRGVQVSTLRLDEYEKIRSCVPQECRDHLVYLTSEALSSNKVWLVGEGTTREEQEKATKGTLFIPFSQFPLK<br>QLRRDCIYHTTPALIVPKSLVNVHSCENWLPRKAMSATRVAGILHALEGWEMHECGTSLLLSDLDQVWEACLSHG<br>FQPLLLPHH | Phytozome |
| *At*CER3 | *Arabidopsis thaliana* | MVAFLSAWPWENFGNLKYLLYAPLAAQVVYSWVYEEDISKVLWCIHILIICGLKALVHELWSVFNNMLFVTRTLRI<br>NPKGIDFKQIDHEWHWDNYIILQAIIVSLICYMSPPLMMMINSLPLWNTKGLIALIVLHVTFSEPLYYFLHRSFHRNN<br>YFFTHYHSFHHSSPVPHPMTAGNATLLENIILCVVAGVPLIGCCLFGVGSLSAIYGYAVMFDFMRCLGHCNVEIFSH<br>KLFEILPVLRYLIYTPTYHSLHHQEMGTNFCLFMPLFDVLGDTQNPNSWELQKKIRLSAGERKRVPEFVFLAHGVDV<br>MSAMHAPFVFRSFASMPYTTRIFLLPMWPFTFCVMLGMWAWSKTFLFSFYTLRNNLCQTWGVPRFGFQYFLPFAT<br>KGINDQIEAAILRADKIGVKVISLAALNKNEALNGGGTLFVNKHPDLRVRVVHGNTLTAAVILYEIPKDVNEVFLTG<br>ATSKLGRAIALYLCRRGVRVLMLTLSMERFQKIQKEAPVEFQNNLVQVTKYNAAQHCKTWIVGKWLTPREQSWA<br>PAGTHFHQFVVPPILKFRRNCTYGDLAAMKLPKDVEGLGTCEYTMERGVVHACHAGGVVHMLEGWKHHEVGAI<br>DVDRIDLVWEAAMKYGLSAVSSLTN | Phytozome |
| *No*CER1 | *Nymphaea odorata* | MASHPGPLTDWPWEKLGSYKYAVLLPFIGHAVHTLYNSEPADRDYTLLCPFYLLVTRLVHDQLWISWSRFQNARS<br>KHQIQSRGIEFQQVDRERRWDDQAIMHALAIYVAHVFIGGASHLPLWNSKGLLFTALVHAGPVEFVYYWAHRALH<br>HYWLFTRYHSHHHSSFVTEPITSVVHPFAEHLLYLAIFAPGFVVPWITGTGSLIALCGYMTFIDLMNNLGHCNFEFIP<br>KWAFTIFPPLKYIMYTPTFHSLHHSQVHINFCLFMPIYDYMYGTVDKTTDTLYETSISGREQMTDVVHLTHPTSIHSI<br>WQIRFGFAYLAAEPYSTKWYFWLLWPFTAALALLTWMFGATFTVEKIRLDKLKIQTWAIPRFGFQYNVASQKQPIN<br>SMIKKAIKDADSKGVKVITLGLHNQSEELNENGKSYLDSVGNMKVKVVDGCSLAAAVVMNNIPQGARQVLVCGR<br>LTKTGYAVVRALCQRGTKVLTVTEEELQGLKSKIPAELHDRFELVHYYDCKVWLVGDGLSTQVQRKAPKGTLFVP<br>FSQFPTKAVRSDCTYHTTPAMAIPKALENVHSCENWLPRRVMSAWRIAGIVHALEGWDAHECGEKMLDMKKVLD<br>AAVSHGFRPLGVVRSM | NCBI |

| | | | |
|---|---|---|---|
| *No*CER3 | *Nymphaea odorata* | MVAPLSAWPWNELCNLKYFLYAPLLANCITSWGNDEAGGWCFHILALCALRGVVHQFWYSYSCMLFITDKYRVL<br>KQGVDYKQIDREHNWDNFIILHAFMAAIACYSSTFMDSLPLFNFRGYIYALILHMGITESLYYFIHRMFHSDYLFQN<br>YHSLHHLSVVAQSYTAGTASILENLVMGFLMGIPMLGASWMGGASVTMFYGYVLVFDFIRCMGHCNVEVMPVSL<br>FDHMPVLRYLIYTPSYHSLHHTDMSSNFCLFMPLYDALGNTLNKNSWEFHRRIRTGEDVRVPDFVFLAHVIDVHAS<br>MHAPFMFRSVSSKAFKANLLVLPLWPVALVVLQLMWAWAKPFLVSFYCLRNRLLHTWIVPRYGFQYFLPFAKDGI<br>NRSIEEAILRADKIGVKVISLAALNKNEALNGGGTLFVGKLPHLRVRVVHGNTLTAAVILHEIPQDVKEVFLTGATS<br>KLGRAIALYLCRKAVRVMMLTHSTERFQAIQKEAPTECQKFLVQVTKYQVAQNCKAWIVGKWLSPMEQAWAPSG<br>THFHQFVVPPILELRRDCTYGKLAAMRLPDDVEGLGNCEYTMGRGIVHACHAGGLVHLLEGWEHHEVGAIDLDRI<br>DVVWECALKHGLRPLS | NCBI |
| *Pp*CER1 | *Physcomitrium patens* | MKAIQPGAWTKFPWHSMGDCKYLLYLSLVGRLLYGLIREDRGRYDLYFHVLLLAVLRHFFGQLGISLSRWPYLSS<br>RYQIQKKGFSFDAVDLSSNWDDYIILDTLLLSVTVMIPMFGNRYYPPWDWTGLVICALLHMGPAEAIYYWLHRAL<br>HGHYLYTRYHSHHHSLFVTEANSGTVHPFLEHLMYASNFAIPLFGTWALGRFSISTLYVYTLTFDTLNAIGHCNVEF<br>VPSWLFDAFPPLKYLIYTPSYHSLHHSQVHTNFCLFMPIYDYWGGTMDKNSDALYRSVRRSDSQERADNVYLTHG<br>MDLLHMMHVTLGIQSFAATPYKGPNWRLWLLYPLALIAMPLLWILGQPFAADKYWIPRTLRGETWLIPRYRFHYS<br>LPVEKVRINALIEQAIVMAEDEGCRVVSLGQLNKEMRLNGSGAAIVVRNPHLKVRIVTGLTLTAAVVINRLPKQTK<br>EVFLVGSSDLIRSVEIYLVRRGVRVLVLTNSPRYFGSTQPKVTKVNQQLIVNVMSFQEGQHCREWILDEYVEGKDL<br>KWAPPGADLHHVCQGSKPLPRTRKDCTYAMYPAMHVPKSMKGLRSCEGGLPRGVISASHAAGVVHSLEKWTHN<br>EVGPIDVERIDTVWAAALKHGFQMAV | Phytozome |
| *Pp*CER1/3 | *Physcomitrium patens* | MATKPGALTEYPWTSLGAWKYTLFLPFAAKAVQTNLLGGHEVDNWCFHMLLSSALRYLHGQAWMSLSRCHWLT<br>GKYRIQTKGINFDQVDRESNWDDYILLHIITATLVHEILPGFANFPVWDLRGIAILLLLHAGPTEFLYYWLHRALHH<br>HFLYNKYHSHHHASFVTEPVSGSVHPFAEHLMYTATFALPFLGTWALGGASIGMFYFYWLFFDFMNAIGHCNFEFF<br>PTWMFRVFPPLKYLVYTPTFHSLHHSHVHTNFALFMPLYDYLGGTADKVSDELYEQVREGKQEKPDFVFLAHGTE<br>LLSTFHLPFGIPSFAAWPYAPKWFIWPLWPLTLPILAILWLFGKPFTSDTYKLKHLRTETWVVPRFGFQYFLPFEKKR<br>INRLIEHAILSAQKKGVRVISLGALNKNESLNGGGTLFVQKHKDLRIRVVHGNTLTAAVILNEIPKDVKEIFLTGATS<br>KLGRAIALYFCHRGVRVLMLTTSRDRFEMIQSELAPQHRENMIQVTKYQAGQNCKRWVLGKWATPSEQKWAPPG<br>THFHQFVVPPVMECRKDCTYGKLSAMQVPKEMKGLRSCEMTMPRGVVHACHAGGLVHALEGWEFHEVGAIDVG<br>RIDETWAAALKQGFKPVC | Phytozome |
| *Pp*CER3 | *Physcomitrium patens* | MVAKEAFLAEWPWERLGHFKYLVYAPFVGRLLQTIIYGTGLEPDNWAMHMFFLMVARYFHQQLWVSASRVPWL<br>TEKFVVDERQSGYEQVDREYHSDNHLMLQLIFISVAHSWFPGFSNVVAWNTQGFLYVLLFHVGVVEVLYYWIHRA<br>FHTEVLFRNYHFYHHMSVVPEPPTGSITTMLEQILQSLLVCVPLLGAAALGGGSMAMIYIYLIAFDFFKCWGHSNFE<br>FVPEWFRGFPGVKYLLYTPSYHSLHHLEQNSNFCLFMPLFDYLGGTVDPKTESLYAELRKGRLLKVPDFVFLAHCI<br>DVLSSLQVSFCCRTMAAHPYKCHWFIWWTWPITVFFLMIFWYWGQTFTAMTIYVNKLKCTSWVIPKHGFQFFLPF<br>GLDSINKHIEKAILEADKQGVKVISLAALNKNEALNGGGLLFVKKHPNLKVRVVHGNTLTAAVIIKTLPPDVKEVF<br>MTGATSKLGRAIALYLCARGIRVLMLTTSTERFDAIQREAPADCRNNLIHVTKYQAGKNCKTWIVGKWTFAKDQQ<br>WAPPGTFFHQFVVPVISEVRKDCTYGQLAGMVLPKEGVKGLRTCEFTMERGAVHACHAGGMIHTLEGWTHHEVG<br>SIDVSRIDVVWEAAMRHGFAPIGS | Phytozome |
| *Ac*CER1 | *Ananas comosus* | SIHDHFLFPLHMHACMQYVVLAPWVLHGLRMVATKGWRELDVITYNAIFPSLLLRMIHNQIWITLSRFQNARSKHQ<br>IVDRSIEFEQVDRERNWDDQIIFNGLLLYVGFTFIPSSAQRLPMWRTDGAVMIALLHAGPVEFLYYWFHRALHHHFL<br>YSRYHSHHHASIVTEPITSVIHPFAEHVVYYALFTIPMLTTVLTGRGSIIALLAYTTYLDFMNNMGHCNFELVPKWLF<br>HAFPPLKYLMYTPSFHSLHHTQFRTNYSLFMPFYDYIYGTMDKSSDDLYESSLKGKEEAPDVVHLTHPTTLQSIYHL<br>RLGFASLASRPYNSKWYMLIMWPISSVSMLLTWIYGSSFTVERNTLKKLKLQTWAIPRYNFQYGLKWEKEAINDLI<br>EKAILEADQRGVRVLSLGLLNQACYSLALAKELNGSGELYIEKHPKLRVRIVDGSSLAAAVVLKSIPAGTNEVLLAG<br>NLSKVARAVAAALCQKGVQVIMTRKHEFKMLKGQMTEKAANYLVFSRNYTTKVWLVGDGLEHEEQKRAPKGA<br>RLIPYSQFPPKKIRNDCSYYTTPCMKIPETLQNMHSCENWLPRRVMSAWRAAGIVHALEGWDVHECGETVQDVEK<br>TWSAAIRHGFLPVTQI | Phytozome |
| *Ac*CER3 | *Ananas comosus* | CDKAKMTSAPFSSWPWENLGVYKYVLYGPLIAKAFTSKAWEFGNPNKWCLHILILFALRGAVHQLWYTFSNMLFL<br>TRRRRIFTDSVDFDQIDKEWDWDNFLILQCLIGAMSLYTFPALRDLPQWDVWGILLAFFLHVTISEPLFYFVHRAFH<br>RGHLFSLYHSLHHSSKVPQSFTAGFATPLEHLILSVVMGVPLLVPCLVGNGSLGLIYGYVLLFDFLRCMGHSNVEVF<br>PHKLFEALPLLKYFIYTPTYHSVHHMEKNSNFCLFMPLFDLLGGTLDNKTWKLHKEISLEVFTSLVSTVVHVWRMV<br>TEERSKFLAGRNDQVPDFVFLAHVLDLVSAMHVQFIFRWHSSMPFAVKPLLLVFWPVAIVIMLCMWAWSKTFLVS<br>MYRLRGRLHQIWAVPRFGFQYFLPFAKDGINHHIELAILRADKMGVKVLSLAALNKNEALNGGGMLFTSKHQDLR<br>VRVVHGNTLTAAVILNEIPKDVQEVFLTGATSKLGRAIALHLCRKKIRVLMLTLSTERFQKIQKEAPPQFQQYLVQV<br>TKIQAAKNCKTWIAGKWLSPREQLWAPPGTHFHQFVIPPIIGFRKDCTYGRLAAMRLPKDVQGLGMCEYTLERGV<br>VHACHAGGVVHLLEGWTHHEVGPIDVDRIDVVWRAALKHGLAPQLHY | Phytozome |
| *Sm*CER1 | *Selaginella moellendorfii* | MAINPGLLTHWPWERLGSFKYLLYLPLVANAVRSAMTPEGRSRDNFSLHILVLAALRYIQGQLWITVTSVHDIVKK<br>HQVQTKGMKFDQLDRERDWEDFILLQALMLLAYQFSPLCLPNHAVSDWRGLVITILWHLGPVEFLYYWFHRALH<br>HHSLYRRYHSHHHLSFVTQAVTGNVHPFAEHLSYAVLFGSTLIVNLFLGTASLALIYSYMLWFDFMNYIGHCNWEF<br>MPSWMFQALPLLKYLVYTPSFHSLHHTQVHTNFCLFVPLYDYIYGTVDKTSGQLHLAARQGRTELVDFVFLTHPTD<br>PLSIFHLSFGIPSFAAQPYGRRWYIWLLYPLALPVMLLLWAFGSPFTVEEHTVDKVLAQTWAIPRFSFHFGMTSEIGS<br>LNALIERAILAAQDKGAKFICLGLHNKDEHLNASGALFLKNHPDLSIKVVDGSTLTSAIVLDKLPKDASEVFLVGAE<br>HKVGRAIANYLCRHRATEVTSLKKSVPQESQHKLVAVESLEHGRHCKAWIVGEPLRAMEQLHAPSGACFYQFTEE<br>AMEETRPDCLYAKLPAMRLPPEYKGIRACEGSMPRGVVQASHAGGILATMENWNHHEVGNTIDVDKIDAVMRAA<br>VNRGFVPYY | Phytozome |
| *Sm*CER3 | *Selaginella moellendorfii* | MEESKILGCWPWQRMGTYKYHLFLPIFLSAAHSHYLGTSPRDNWCFHILVIAALRYALYQAWSSFARLHAVVKHH<br>QIISYALTYEQVDREFDCDNGIILHSLLAYALGPNDISGFSIWNLRGLVYLIAFHAGVTESAYYWLHRAFHTKSLFRS<br>FHSYHHASTAPEPATAFTHTFLEALLQTVLMSVPIFASCFLGGSCLALFYVYPLAFDFFKYLGHFNCEIVPLWAFQKL<br>PLLKYLIYTPSYHSLHHLDLKSNFCLFMPLYDYLGGTQHPNTHAFYRSIRKDGREAVPQFVFLVHCIDILSSLHVAFS<br>GRTASSVPFRGEWYAWLVFPIGLVSCFCVWIWGKTFVATKYLLDGLHAQSWVVPRYGFHYFIPACAAGINRHIERA<br>ILDADELGVKVISLAALNKNESLNGGGLLFVKKHPNLKVRVVHGNTLTAALVLRELPAETSEVFLTGSTSKIGRAIA<br>LYLCRRNVRIMMLTTSRERYQSIVDEAPADCRHNLVQVTKYQAGQTCKTWIVGKWATSQDQSWAPHGSHFHQFV<br>VPPVHEYRKDCTYGKLAGMKLPQSVEGVHSCEYTFDRGVVAACHAGGLVHALENWTHHEVGSIDIDHIDLVWEA<br>ALKHGLEPVL | Phytozome |

| | | | |
|---|---|---|---|
| *Af*CER1 | *Azolla filiculoides* | MEWMWMWMVLGRRRRRRQYLAYVLVGGAALWEGSPWTTNISLNLLLLATLRYTTFQIWGSVSRLPFFYRPFIIQ<br>KKSIPIDQVEHEASWENSVIFTTLLASSFKVMFPSIFTLPLWDGKGVALALLLHVGPVEWLYYWGHRALHHHYLFS<br>RYHTHHHSSFVTQPVSANNHPMLEIMFYTMLFAIPIVGAVLVGSASIAMYHIYVFFVDTMNAWGHCNFEFIPTSVY<br>DAFPPLKYLIYSPSFHSLHHSKVHTNFALFMPLYDYMYGTADPTSDELHRQVREGNHDKVDCVYLTHPTDLPSIFH<br>QPFAFQTFATQPYSKPWYVWLTWPFSFVIMVAFWLFAKPFVATSQLIGKSFELQTWMIPRYNFQYRMKPVSAKIKR<br>LVEQCVLDAQEAGAKVISLGLLNKELLDGDFLRTRSIEIPIVTGETLTSAIIVKKIEAQKVDKVFLASANSRVGQAVA<br>TYLCQHGVTVVALLPTKSCFHELQANIDAEFQKNLVFATTYKNGKDCKVWVVNDMVAAEEQRWAPAGTCFHHL<br>GVFPLSTTTRTSDCTYEYIPAMRLPSNARTMNGCEDVLPKYVVRAAYAAGMLHALEGWKHSDFSNEISTYLLDKT<br>WEAAVKHGFSIYSLNGDDQAKH | FernBase |
| *Af*CER3 | *Azolla filiculoides* | MALPSAPAPLYSWPWRWMGCFKYLLYLPIAWAAYSSLEQKTPISQNFSLHILILTAARLILYQLWQTFSRFLPLTESI<br>QIRPQGLSYAQVDREWHCDDHHLLQALVLFGAHSWLGFLKGAPVWRGSGLLMTLLLHTGPAEMIYYWFHRFLHS<br>APMFERYHKLHHESVTPEPSTPGLSTFLEQISMVVIMAVPIVGGSVKGQGSLSALYAYVLLFDCMRCMGHCNAEMF<br>SPTFLANFPLIKYLIYTPSYHTLHHEERDSNYCLFMPIYDYMFGTVSKNREAFLKEIRERVLRVPDYVFLLHAVDFLS<br>CLHVHFLFRAFASHGYRTLLFLIPIWPFVIPIAIAMWIWGKPFKGWNYYLDNRFHQIRLIPRFGFMYFLPFARNNINN<br>LIEKAILDADRMGVKVLSLAALNKNEALNGGGNLFVMKHPNLNIRVCHGNTLTAAVIIKEIPDHVTEVFLTGATSK<br>LGRAIALYLCQKGVRVLMLTESRKRFESITSESPAEFRHNLVQVTKHQAGKNCKTWILGKWTTYSDQMFAPPGTHF<br>HQFVVPPVIPFRRDCTYGALAAMRLPPNTKGLASCEYTLDRNVVHACHAGGVVHALEGWTHNEVGALDVDRIDL<br>VWEAAMRHGFRPA | FernBase |
| *Sl*CER1 | *Solanum lycopersicum* | MASKPGILTEWPWTWLGNFKYVVLAPFVGRSIESLLNREDGSKIDIGYLIIFPFLLFRMLHNQIWISLSRYKTAKGDN<br>RILDKTIEFDQVDRERNWDDQILLNGLLFYYGYMKLEQSHYLPIWRSDGIILTTLLHIGPVEFLYYWLHRALHHHFL<br>YSRYHSHHHSSIVTEPITSVIHPFAEHIAYFLLFSIPLLTTVVTKTASIVSFGGYITYIDFMNNMGHCNFEIIPKWMFSTF<br>PPLKYLMYTPSYHSLHHTQFRTNYSLFMPIYDYIYGTLDKSSDTLYEKSLERQGKSPDVVHLTHLTTPESIYHLRLGF<br>ASFASQPYTSKWYFWLMWPVTLWSMMVTWIYGHTFTVERNVFNNLNLQTWAIPKYRVQYFMQWQRETINNLIEE<br>AIMEADQKGIKVLSLGLLNQDEKLNKNGEVYIRRHPQLKVKLVDGSSLAVAVVLNSLPKGTTQVVLGGHLSKVAN<br>AIALALCQGGVKVMTLREEEYKKLKSSLTPEAATNLLLSKTYTSKIWLVGDGLNEDEQLKVPKGTIFIPFSQFPPRKT<br>RKDCFYFHTPAMITPKHFENVDSCENWLPRRVMSAWRIAGILHALEDWHEHECGNLMFDIEKVWKASLDHGFQPI<br>SVVSASESKA | Phytozome |
| *Sl*CER3 | *Solanum lycopersicum* | MEKQNEEVENGNDTLLRKTDAALFAWPWNNLGNYKYLLYGPFLAKFIHSMYWKESMEDIWCLHILVLCSLRGLV<br>HQLWSTFSNMLYLNSTRRVSYEGIDYDQIDNEWDWDNFLILQAVVGSFVYLNFPSLANLPVWDVRGLISCLILHIGI<br>SEPLFYWMHRLLHSSYLFPLYHWHHHESKITHPFTAGHGTFLEHLLLCVVIGIPTLGTAFIGYGSISVMYSYILAFDFL<br>RCMGHSNVEIIPHSLYQRVPLLRYVIYCPTYHSLHHQEMKTNFCLFMPLYDMLGNTLNTASWSLHKEISSRTNERAP<br>DFVFLAHIVDIMSSMHAPLLFRSFSSVPFSTRLFLLPMWPFAFVVVLTMWLKSKTFLFSFYNIRGRLNQTWIVPRAGF<br>QYFLPFAAEGINKLIEEAILRADRIGVKVISLAALNKNESLNGGGTLFVNKHPNLRVRVVHGNTLTAAVILNEIPRDV<br>NEVFLTGATSKLGRAIALYLARRRVRVLMLTKSTERFMKIQREATVECQKYLVQVTNCKEAKQCKTWIIGKWSTP<br>REQSWAPSGTHFYQFVVPPIIPFRRDCTYGKLAAMRLPKDVTGLGTCEYTMGRGIVHACHAGGLVHLLEGWTHHE<br>VGAIDVDQIDVVWEAALKHGLKPLYNSS | Phytozome |
| *Psit*CER3 | *Picea sitchensis* | MSWKSCFLLDWPWAYLESLKYVLYGPLIAKAVHTNLYGGKEADNWCFHILLLTSLRYVTYQLWATFSNMYCLSH<br>RYKICKKGAEFDQMDREWDWDNFLLLQAFMATAAHHFLPFFRDMPAWNAGGLICLAILRMGPAEVLYYWAHRA<br>FHKDFLFQRYHSLHHAAIVLQPQTAGTATFLEHIGLTIIMAVPMVGASWMGGASMGMIYIYCLLFDFLRYMGHSNV<br>EIVPETIFRCLPPLKYLIYTPLYHTLHHTEMDTNFCPFMPLYDYLGHTINSKSWDLHRSMSAGQVEDVPDYVFLAHI<br>VDVLSSLHVRFLLRGFCSTPFATWWFLLPLWPVVIPVALAMWVWAKTFVNTGHRLKGRLHQTWIVPRFGFQYFIP<br>FAQAGINNLIQDAILSADKMGVKVISLAALNKNEALNGGGTLFVNRLPDLRVRVVHGNTLTAAVILNELNPDVKEV<br>FLTGATSKLGRVIALYLCRKGIRVMMLTYSKERFKSIQSEAPPEFQNFLVQVTKYEAAQNCKTWIVGKWIAYKEQT<br>WAPVGCHLHQFVVPPIFELRKDCTYGKLAGMQLPDAVEGLSTCEYTMPRRCVHACHAGGILHSLEGWEHHEVGAI<br>DVNKIDMVWEAALKHGFKPMK | NCBI |
| *Zm*CER1 | *Zea mays* | MASKPGPLSRWPWQDLGNYKYALVAPWAVRSTYRFVRSGSGERDLLAFAVLPVLLLRLLYSQLWITVSRHQTARS<br>RHRIVNKSLDFEQVDRERNWDDQILLTALLFYAVNAAVPVAQSVPWWDSRGLLVAALLHVGPVEFLYYWLHRAL<br>HHHYLYARYHSHHHASIVTEPITSVIHPFAEELVYFTLFAIPLLTMVGTGTASVAVANGYLAYIDFMNYLGHCNFEL<br>VPRLLFDVFPPLKYLMYTPSFHSLHHTQFRSNYSLFMPLYDHLYGTADKSSDDLYERALQGRAGEDAPDVVHLTHL<br>TTPASLLRLRLGFASLAAAPAPPASRYGAGSSSSSSSSLAAVACPLAALLGWTRTAFRSEANRLHKLKLETWVVPRY<br>TSQYLSKQGLYAVGRVVEKAVADAEASGARVLTLGLLNQANELNKNGELYVIRKPSMRTKIVDGTSLAAAAVLH<br>MIPEGTDEVLLLGDAGGNKMAGVLASALCEREIQVHVVDKDLYESVKQQLRPETHEHLLHLAEWWSHSAKTTKV<br>WLVGDRLTGEEQRKAQGGAHFVPYSQFPPGAVVRADCVYHSTPALVVPDAFEDLHACENWLPRRVMSAWRAAG<br>IVHALEGWDAHECGARVTGVDKAWRAALAHGFRPYDRYGAN | Phytozome |
| *Zm*CER3 | *Zea mays* | MGKPRIIQGKQTTTRAAALLILGSLLFVWTGVHAQYLLYGPLVAKVAHAWRETGSLPLGSWCLHLLLLLALRSLTF<br>QLWFSYGNMLFFTRRRRVVKDGVDFRQIDAEWDWDNMVILQTLVAAVAMGSAAFPAVSELRAWDPRGWALALL<br>LHMAVSEPVFYWTHRALHRGPLFSQYHARHHSSPVTQPFTAGFGTPLEALLLTLAMGAPLAGAFLAGAGSVSLVY<br>GHVLLFDCLRCLGYSNVEVISHRAFAAFPPLRYLVYTATYLSLHHREKDCNFCLFMPLYDALGGTISSRSWGLQRE<br>VDQGMNDRVPDFVFLAHVVDVVSSMHVPFAFRSCSSLPWAMRPVLLPLWPVAFAFMLLQWFFSKTFTVSFYFLRG<br>RLHQTWSVPRYGFQYFIPSAKKGINRQIELAILRADKMGVKVISLAALNKNEALNGGGTLFVNKHPNLRVRVVHGN<br>TLTAAVILNEIPSSVREVFLTGATSKLGRAIALYLCRKRIRVLMLTLSTERFLKIQREAPPEFQQYIVQVTKYQAAQG<br>CKTWIVGKWLSPREQRWAPPGTHFHQFVVPPIIGFRRDCTYGKLAAMRLPKDVEGLGSCEYTMERGVVHACHAG<br>GVVHCLEGWEHHEVGALEVDRIDVVWEAALKHGLTPA | Phytozome |
| *Atri*CER1 | *Amborella trichopoda* | MASKPGPLTDWPWKKLGNFKYMVIVPCAIHAVYRIYGAQKQKRDHATIISLGLFLSRIIHDQLWITLSRFQTARSKH<br>RIQSRGIDFEQVDRERNWDDQAILHAIIFYIGELFLPGAQNLPLWNTKGVIIAVLCHAGPVEYIYYWAHRALHHHFL<br>YTRYHSHHHSSFVTEPITSVVHPFAEHILYALIFAVPFLATVLTYTASYTVLIGYMTFIDFMNNMGHCNFEFFPKWTF<br>KIFPPLKYLIYTPSYHSLHHSQVHTNFSLFMPIYDYMYNTVDKTTDSVYESSIEGREDTTDVVHLTHPTSLHSIYHLR<br>LGFAYLAAEPYSSKWYLWLMCPLSFVLMLVTWIFGCTFVVEKNKLYEHKMQTWAIPRYNFQYSLKWQKAPINNM<br>IEKAILEADAMGVKVISLGLLNQGEEFNKNGELYLHNREKLKTRIVDGSTLSAATVLNGIPHGTQKVLIRGCLTKTL<br>FTTALALLERRTKIVVIRTEEYENLKMRIPSKYQSGISLSNNYDTKVWLVGEDLRAEEQSRANRGTLFIPFTQFPPRA<br>VREDCIYHTTPALVIPRTLEDVHSCENWLPRRVMSAWRVAGIVHALEGWDSHECGNMTLDVEKVWSTTLSHGFKP<br>INNVKFN | Phytozome |

| | | | |
|---|---|---|---|
| *Atri*CER3 | *Amborella trichopoda* | MLFLTKKLRVLQQGVDFKQIDREWHWDNFILLQALMATFACYYFPSFTKSLPLYNTTGFICVLLLHMGFSESLYYG<br>VHRLFHSDYLYTNYHSFHHKSMVSQSFTAGSASLLEHIVLCFIMGIPIVGASWMDCASLSMVYAYIMAFDFMRCLG<br>HCNVEIIPHSLFEWMPLLRYIFYTPTYHSLHHNEKSTNFCLFMPIYDALFNTLNPKSWDLHRTVRSGCGDKVPDFVF<br>LGHVIEFDSALHVSFLFRSISSMPCVFRPILLPTWPVVFVVMLFMWASATTFLISFYTLRDRLHQTWVIPRFGFQYFLP<br>FAKDGINKHIEDAILSADKLGVKVISLAALNKNEALNGGGIIYVQKHRNLKVKVVHGNTLTAAVILNELPNGVKEV<br>FLTGSTSKLGRAIALYLARKGVRVLMLTLSQERFQIIQKEAPVEYQKNLVQVTKYQAGQNCKTWIVGKWILPREQK<br>WAPSGTHFHQFVVPPILRLRRDCTYGKLAAMHLPDDVEGLGTCEYTMGRGIVHACHAGGVVHLLEGWTHHEVGA<br>ISVDRIDLVWEAALKHGLKPV | Phytozome |
| *Sf*CER1/3 | *Sphagnum fallax* | MASSPGVWSEWPWQRMGAWKYTLFAPYAVKAVHANFLGGQDVDNWCLHMLIVSALRYLHGQIWMSLSRCHTL<br>TGKYQIQTKGITFEQVDRESNWDDYILLHVIVATIVHVVLPGFCNFPVFWDQKGIIILLLLHMGPAECIYYWLHRAL<br>HHHSLFSRYHSHHHASFITEPVTGSVHPFAEHLMYTAIFAIPFLGTWALGGASMGMFYFYWLFFDFLNAIGHCNFEF<br>MPTWMFQVFPPLKYLVYTPTYHSLHHSRVHTNFSLFMPIYDYLGGTVDATSDELHSFVRKSQQQEKPDFVFLAHGT<br>ELLSAFHLPFGIPSFAAWPYSVRWYLWLLWPLTLPVLGMTWIFGRPFVSDKYRLPNLRTETWVIPRFGFQYFLPFEK<br>KAINKLIEQAILEAENEGARVIGLGALNKNESLNGGGTYFVNKHKDLRIRIVHGNTLTVAVILNTIGTDVKQIFLTGA<br>TSKIGRAVAIYLCRQGVRVMMLTASVDRYESILAEVPVEYKQNLVQVTKYQDGQYCPVWVIGKWATPNDQKWAP<br>PGTHFHQFVVPNIIESRQDCTYGKLAAMHLPKEIKGLRSCEMTMPRGVVHACHAGALVHALEQWTHHEVGVVDID<br>RIDITWAAALKHGFKPVQ | Phytozome |
| *Sf*CER3 | *Sphagnum fallax* | MGVRTPIFFEWHWERLGNFKYVLAAPVVANGIHGLISGAKDVDNWCYHLLLLGALRCLCQQMWTTYSRLHCLVK<br>KHQINACGIEFEQVDREFHSDNHMLLQLLLLLAAHSWIPGFSNLPMWNMRGLVILFLLHTGPVEFLYYWMHRAFH<br>SEPLFQRYHSLHHLSVVTEPPTGSVTTMLEQVLQTGLIGLAMVVTVMLGGASISMIYIYLLTFDFLKCMGHCNCEFV<br>PARVFQLFPAAKYLLYTPSYHSLHHTQLHYNFCLFMPLYDYLGGTVHPDTLPFYASLRKGKEEEVPDFVFLAHCID<br>VLTSLHVSFCIRTLAAHPFTPYWFIRLMWLPTLFCLFIFWTWAQTFVAYRYMLDNLNCVTWVIPRHGFQYFLPFYK<br>DNINKHIEKAILNADTLGVKVISLAALNKNEELNGGGMLFVEKHKDLRVRVVHGNTLTAAVIIRQLPTNVKEVFMT<br>GSTSKLGRAIALYLCRRGVRVLMLTKSKERFDKIVAEASHDVRHNLIRVTKFQAGQNCKTWLFGKWTLNADQQW<br>APPGTNFHQFVVPAVREIRKDCTYAPVAGMILPKQTIGIHTCEVMMPRRAVHACHAGGLVHALEGWKHHEVGAID<br>VDRIDVVWEAAMKHGFSSVT | Phytozome |
| *Mp*CER1/3 | *Marchantia polymorpha* | MATAPGALYDWPWQKLGSFKYLLFVPFVVKATHVNVFGGHEADNWCLHMLISTALRYLHGQIWMSLSRFYHLTE<br>KYQIQKKGINFQQVDRESNWDDFTILQLIVITIVHSYIPGFSNIPLWDTRGIIYLLLLHAGPAEFIYYWAHRALHHHYL<br>YTRYHSHHHASFVTEPISGNVHPFAEHILYTAIFAVPMLGSWALGGVSMGMVYFYWLWFDFMNAIGHCNWEFMP<br>PWAFKAFPPLKYLIYTPTYHSLHHSQVHTNFALFMPLYDYLGGTVDKNSDSLYETVRKGNSNKLDFVFLAHGT<br>ELLSTFHLPFGIASFAARPYKAQWYMWPLWPLTLPVMALLWVFGHPFVADNYRLDDLFTQTWVIPRYGFQYFLNF<br>EKNRINRLIEEAILKAQDKGCKVISLGALNKNEALNGGGTLFTEKHPDLKIRVVHGNTLTAATILDKIPRDVTEIFLT<br>GATSKLGRAIALYLCRRGVRVLMLTTSRDRFESILHEAPVDCQRYLVQVTKYQDGADCKTWVMGKWATPKQQSY<br>APSGTHFHQFVVPPLEPARKDCTYGTLAAMQLPKAMRGLRTCEMTMPRGCVHACHAGGLVHALEGWTHHEVGA<br>IDVERIDLTWKAAMKHGFFPVD | Phytozome |
| *Mp*CER3 | *Marchantia polymorpha* | MGTKLNFMWSSPWQILGGYKYVIGIPFLVKAIHANYYGGYDVDNWCFHMIWVTIMRLTLSQLWQIYSRLHGIVKK<br>HQISTVGFTFEQIDREFHSDDYIILQALVATAAHVWVPGFRQLPLWDSRGLVTIILLHVGPTEFVYYWLHRALHTDF<br>LFTNYHSFHHASINTEPPTSGVGTFLEHVLFSGVMGIALAGPVIFGGASISMIYLYCLFFDFMKHMAHSNTEIIPITLFK<br>AFPLLKYLLITPSYHSLHHSELHSNYCLFMPLYDHLGGTVNVKSEALHARLREGRAEEVPEFVFLAHCVDLLSSMH<br>VSFVLRQFASRPYSARWFLYPFLVLITPVMFMMWAWGKVFVAYKYTLDKFSCQTWVVPRYGFQYFIPVGLNSING<br>LIESAILEADKKGVKVISLAALNKNEALNGGGVLFTDKHTNLRVRVVHGNTLTAACILKGIPEDVTEVFITGATSKL<br>GRAIALHLCRRRVRVLMLTSSQERFEAILKEAPKELKRYLVRVTKYQAGINCKTWIMGKWISHKDQQMAPRGTHF<br>HQFVVPGIPEIRKDCTYGKLAAMRLPTQVKGISTCEFSCERGVVHACHAGGLIHCLEGWTHHEVGSIDVDRIDEVW<br>EAAMRHGFSAVY | Phytozome |
| *Dc*CER1/3 | *Diphasiastrum complanatum* | MATKPGALTDWPWKKLGNWKYILFAPFVVKAVHANLLGGHDADNWSLHMLLTSALRYLHAQIWMSLSRCHFLT<br>SKYQIQSKEITFEQVDRERNWDDYILMYIFGMTVLHAMLPGFSNFPVFWDSRGIVMLLLLHMGPTELVYYWAHRA<br>LHHHFLYTRYHSHHHASFITEPITGIVHPFAEHMMYMAIFAIPILGTWALGGELSVGMLYFYWLFFDFMNALGHCN<br>FEFVPTWAFQVFPPLKYLIYTPSYHSLHHSHVHTNFALFMPLYDYLGGTVDKISDELYETVRQGRKEKIDFIFLAH<br>GMELLSTFHLPFGIPSFAAWPYAPKWYLWPLWPLTLPILAILWIFGKPFASDKYRLLNLHTETWVVPRFGFQYFLSF<br>EKKRITRLIEQAILEAEKGGARVISLSALNKNESLNGGGTLFVNKHKDLRIRIINGNTLTAAVILNKLPTDLKQIFLTG<br>AMSHIGRAIAIYLCQRGVQVMMLTTSQDQYETILAEVPVQYRGNLVQVVKYQDGQHCSVWVIGKALTPSEQKWA<br>PSGTHFHQFVLAPILESRKDCIYEKLPAMQVPKDMKGLRSCEMTMPRGVLHACHVGGIVHTLERWEHHEV<br>GAIDVDRIDQTWDAAVKHGFKPM | Phytozome |
| *Dc*CER3 | *Diphasiastrum complanatum* | MALLRDWPWERIGNYKYYLLLPLVIKLFHARYWGGSDTDNWGFHILLICSLRYLQHMCWMSFSRLYCLVKRYQV<br>QSCGTDFDQVDREFHFDNYIIFQALVATIAHEYLPGFKNLPLWNSTGVLYLAIFHMGATESLYYWVHRALHSEAFW<br>QYHSKHHASTTPEPSTSGTHTFLEEILQSGMMALPIVGAGLMGASSISSLYIYILAFDFLKQMGHCNCEVVPMWAFQ<br>IFPPMKYLLYTPSYHSLHHTDCNSNFCLFMPIYDYLGGTVNKRTRSLYLSLRKGNKNAVPDFVFLAHMIDIMSCLHV<br>SFVDRTLASVPFAPRWFLWLQWPFVLPFICALWIWGRTFVAYEYMLRGLHAQTWVVPRCGFHYFLPFEKERINQLI<br>ESTILDADVLGVKVISLAALNKNEALNGGGLLFVNKNKDLRVRVVHGNTLTAAVILYDLPKSVREVFMTGATSKL<br>GRAIALYLCNKNVRVLMLTSSRERYESILKEAPEDCRQNLIRVTKYQAGKNCKTWIIGKWTTGRDQSWAPSGTHFH<br>QFVVPPVQELRKDCTYGKLAGMRLPQDIKGLRTCEYTFGRSVVAACHAGGLLHALEGWEHHEVGSIDVDRIDVV<br>WEAALKHGMTPTWEDN | Phytozome |
| *Cr*CER3 | *Ceratopteris richardii* | MAIGLASWPWERMGNYKYALYLPLVWEVYQTSIPLRQNVCVHILAVTFLRVVVYGLWQAASRFLPLVDGLHIRPQ<br>GFSYRQIDREYHCDNPHLLQALLLYAGHKWVGLFECMPFWSTRGLVLVLLLHVGPAEMLYYFIHRALHTSYFFEN<br>YHKLHHESVTPEPSTAGVSSFLEQIWISAIMGIVVIGSTVIGGASVFNLYAYVLFFDCLRCYGHSGIEIVPVSIFKFFPL<br>LRYFLYTPSYHALHHECQDCNFCLFMPFYDYFGNTVNPRTQDFHAESRDKSTGRVPDFVFLLHAVDFTSALHSLFV<br>FRTFASRPYASSIFLIPFLPIVVLIASGMLIWGTAFRYWNYYIRTYHIQVWLVPRFGFMYFLPFARDNINSLIEKAILDA<br>DKTGVKVFGLAALNKNEALNKGGSLFVAKHPNLRVRVCHGNTLTAAVILKEIPIHVKEVFLTGATSKLGRAIALYL<br>CRKGIRVLMLTESRKRFEAIFAEAPSNVRNNLVHVTKHQAGKNCKTWILGKWLTYWEQMFAPSGTHFHQFVVPPV<br>VPFRRDCTYGKLAAMRLPDDVRGLPCCEYALDRNVVHACHAGGLVHLLEGWTHHEVGALDVDRIDVVWNAALK<br>HGLLPV | Phytozome |

| | | | |
|---|---|---|---|
| *Tp*CER1 | *Thuja plicata* | MATKPGPLTQWPWEKLGNFKYLLLAPLVIKAIHTNFLGGYEADNISFVLLLLTAVRYIHNQLWISISRYQNARSKHQ<br>IQSKSIHFEQVDREGHWDDYILLYTFIFVIVHAYIPGASNVPLWNTKGLFVISLAHVGPSEFIYYWAHRALHHHFLFS<br>RYHSHHHSSFVTEPITSVVHPFAEHLMYVTMFSVALLATLFTNTQSVGLIFGYMMWFDFMNNLGHCNFEFIPKWAF<br>KVFPPLKYLMYTPSYHSLHHSQVHTNFCLFVPLYDYLYGTVDKSTDTLYETAYKGRQEKVDLVYVTHATSLFSFFH<br>LRFGFASFAAKPYSTKWYLWILWPISNAIMLFLWLFGKTFVVEKNRLNDLHLQTWVIPRYTFQYYMSSERARINKFI<br>EDSILEADKKEVKVINLGLLNQSEDLNGGGELFLKKYKNLKVRMVDGSTLAAAVVLNSIPLETTEVFMCVGRSKIG<br>SAIVCLLSQRGVTIQLLTESREQLDQMKSNVPSQFQHNIVPVNSYQEGKNCKIWIVGRMVNGEDQRKAPKGAIFIPIL<br>PFPIHRTRKDCIYHSIPAMKVPANVENVHACENWLPRKVMSAWRVGGMVHALEEWNHHECGQTMNIADIDKVW<br>KAALKHGFLPFNTSPAQLQLAFKICSSSFYKYIYKVKHRNA | Phytozome |
| *Tp*CER3 | *Thuja plicata* | MEQGRGPLFGWPWERLNNFKYLLYAPLLFKAADEIWENRGEGYWFVHILVISFLRCFLYQVWHSYSRAGFLSRRH<br>QIHWNSVDFDQIDREFHWDNYVILQAWLATVGHKLVPGLVGMLPMWNSRGILTVLLLHMGPTELLYYYFHRAFH<br>KGYLFQNYHYLHHESTRTEPSTSWTSSFLEQLVMVALMSIPIGAAAVMGEACIGMFYVYLLGFDFLKFMQHSNVE<br>VVPLWLFDKLPFLKYLICTPSYHSIHHTKLDCNYCLFMPLYDYLGGTSNSKESSILHTKLRTQGQERKVPNVVFLC<br>HLVHYIHSLHIPYFGRTFSARPCTYGAWYAWIYLPLIWICFPIASLLTSAHAISKYHIKGSFFVETWGIAKSGFQFFLPF<br>GKNHINKLIEDAILYADRLGVKVFTLGALTKNEVLNGGGELFVRRNPNLSIRVVHGNTLTAAVLLHELPENVDQVF<br>LSGSTSKIGKAIALYLCRKRVRVLMLTASLERFKAIKNEAPAEYRDYLVHSSSYEGGSRCKTWIVGKWAGYKNQK<br>WAPTGTQFYQFTLPKIIEFRSDCTYRDSVSLMLPRAAEGVNCCEYTMPRRVVHACHAGGVVHALEGYENHEVGAI<br>NVDHIDLVWNAAMKHGFTLVPQPLVK | Phytozome |

**Table S2: Plasmids constructed and used in this work for protein expression in yeast. All plasmids were assembled using Golden Gate cloning.** The replication origins are shown in bold red, promoters in bold blue, coding sequences (CDSs) in bold orange, and terminators in bold pink. Abbreviations: 2micron: A yeast origin of replications, ScURA3p: *S. cerevisiae* orotidine 5-phosphate decarboxylase promoter, URA3: *S. cerevisiae* orotidine 5-phosphate decarboxylase, ScURA3t: *S. cerevisiae* orotidine 5-phosphate decarboxylase terminator, ScCUP1p: *S. cerevisiae* copper inducible promoter, ScPGK1t: *S. cerevisiae* 3-phosphoglycerate kinase terminator, ScTDH1t: *S. cerevisiae* glyceraldehyde-3-phosphate dehydrogenase 1 terminator, ScADH1: S. cerevisiae alcohol dehydrogenase terminator.

| Plasmid name | Description | Genetic features |
|---|---|---|
| pFV261 | L2 empty plasmid | - |
| pFV334 | Empty vector (EV) plasmid | **2micron-URA3p-URA3-URA3t** |
| pFV570 | *At*CER1-1 plasmid | **2micron-ScURA3p-URA3-ScURA3t-ScCUP1p-*At*CER1-ScPGK1t** |
| pFV571 | *At*CER3 plasmid | **2micron-ScURA3p-URA3-ScURA3t-ScCUP1p-*At*CER3-ScTDH1t** |
| pFV573 | *At*CER1-1 + *At*CER3 plasmid | **2micron-ScURA3p-URA3-ScURA3t-ScCUP1p-*At*CER1-1-ScPGK1t-ScCUP1p-*At*CER3-ScTDH1t** |
| pFV575 | *At*CER1-1-H159A + *At*CER3 plasmid | **2micron-ScURA3p-URA3-ScURA3t-ScCUP1p-*At*CER1-1-H159A-ScPGK1t-ScCUP1p-*At*CER3-ScTDH1t** |
| pFV896 | *At*CER1-1 + *At*CER3-C587A plasmid | **2micron-ScURA3p-URA3-ScURA3t-ScCUP1p-*At*CER1-1-ScPGK1t-ScCUP1p-*At*CER3-C587A-ScTDH1t** |
| pFV897 | *At*CER1-1 + *At*CER3--A461N-T462L plasmid | **2micron-ScURA3p-URA3-ScURA3t-ScCUP1p-*At*CER1-1-ScPGK1t-ScCUP1p-*At*CER3-A461N-T462L-ScTDH1t** |
| pFV796 | *No*CER1-1 + *No*CER3-1 plasmid | **2micron-ScURA3p-URA3-ScURA3t-ScCUP1p-*No*CER1-ScPGK1t-SScCUP1p-*No*CER3-1-ScTDH1t** |
| pFV576 | *Ot*CER1/3 plasmid | **2micron-ScURA3p-URA3-ScURA3t-ScCUP1p-*Ot*CER1/3-ScADH1t** |
| pFV818 | *Kn*CER1/3 plasmid | **2micron-ScURA3p-URA3-ScURA3t-ScCUP1p-*Kn*CER1/3-ScADH1t** |
| pFV817 | *Pc*CER1/3 plasmid | **2micron-ScURA3p-URA3-ScURA3t-ScCUP1p-*Pc*CER1/3-ScADH1t** |
| pFV819 | *Mk*CER1/3 plasmid | **2micron-ScURA3p-URA3-ScURA3t-ScCUP1p-*Mk*CER1/3-ScADH1t** |
| pFV816 | *Ps*CER1/3 plasmid | **2micron-ScURA3p-URA3-ScURA3t-ScCUP1p-*Ps*CER1/3-ScADH1t** |
| pFV579 | *Ot*CER1/3 + *At*CER1-1 plasmid | **2micron-ScURA3p-URA3-ScURA3t-ScCUP1p-*Ot*CER1/3-ScADH1t-ScCUP1p-*At*CER1-1-ScPGK1t** |
| pFV582 | *Ot*CER1/3 + *At*CER3 plasmid | **2micron-ScURA3p-URA3-ScURA3t-ScCUP1p-*Ot*CER1/3-ScADH1t-ScCUP1p-*At*CER3-ScTDH1t** |
| pFV832 | *Kn*CER1/3 + *At*CER3 plasmid | **2micron-ScURA3p-URA3-ScURA3t-ScCUP1p-*Kn*CER1/3-ScADH1t-ScCUP1p-*At*CER3-ScTDH1t** |
| pFV831 | *Pc*CER1/3 + *At*CER3 plasmid | **2micron-ScURA3p-URA3-ScURA3t-ScCUP1p-*Pc*CER1/3-ScADH1t-ScCUP1p-*At*CER3-ScTDH1t** |

| | | |
|---|---|---|
| pFV833 | *Mk*CER1/3 + *At*CER3 plasmid | **2micron-ScURA3p-URA3-ScURA3t-ScCUP1p-*Mk*CER1/3-ScADH1t-ScCUP1p-*At*CER3-SsTDH1t** |
| pFV830 | *Ps*CER1/3 + *At*CER3 plasmid | **2micron-ScURA3p-URA3-ScURA3t-ScCUP1p-*Ps*CER1/3-ScADH1t-ScCUP1p-*At*CER3-ScTDH1t** |
| pFV701 | *Ot*CER1/3-H142A Plasmid | **2micron-ScURA3p-URA3-ScURA3t-ScCUP1p-*Ot*CER1/3-H142A-ScADH1t** |
| pFV702 | *Ot*CER1/3-H155A Plasmid | **2micron-ScURA3p-URA3-ScURA3t-ScCUP1p-*Ot*CER1/3-H155A-ScADH1t** |
| pFV703 | *Ot*CER1/3-H243A Plasmid | **2micron-ScURA3p-URA3-ScURA3t-ScCUP1p-*Ot*CER1/3-H243A-ScADH1t** |
| pFV812 | *Ot*CER1/3-C581A Plasmid | **2micron-ScURA3p-URA3-ScURA3t-ScCUP1p-*Ot*CER1/3-C581A-ScADH1t** |
| pFV813 | *Ot*CER1/3-A455N-T456L plasmid | **2micron-ScURA3p-URA3-ScURA3t-ScCUP1p-*Ot*CER1/3-A455N-T456L-ScADH1t** |
| pFV704 | *Ot*CER1/3+*At*CER3-H147A plasmid | **2micron-ScURA3p-URA3-ScURA3t-ScCUP1p-*Ot*CER1/3-ScADH1t-ScCUP1p-*At*CER3-H147A-ScTDH1t** |
| pFV705 | *Ot*CER1/3+*At*CER3-H161A plasmid | **2micron-ScURA3p-URA3-ScURA3t-ScCUP1p-*Ot*CER1/3-ScADH1t-ScCUP1p-*At*CER3-H161A-ScTDH1t** |
| pFV706 | *Ot*CER1/3+*At*CER3-H250A plasmid | **2micron-ScURA3p-URA3-ScURA3t-ScCUP1p-*Ot*CER1/3-ScADH1t-ScCUP1p-*At*CER3-H250A-ScTDH1t** |
| pFV892 | *Ot*CER1/3+*At*CER3-C587A plasmid | **2micron-ScURA3p-URA3-ScURA3t-ScCUP1p-*Ot*CER1/3-ScADH1t-ScCUP1p-*At*CER3-C587A-ScTDH1t** |
| pFV893 | OtCER1/3+AtCER3-A461N-T462L plasmid | **2micron-ScURA3p-URA3-ScURA3t-ScCUP1p-*Ot*CER1/3-ScADH1t-ScCUP1p-*At*CER3-N461N-T462L-ScTDH1t** |

**Table S3:** ***S. cerevisiae*** **strains constructed and used in this work.**

| Strain name | Plasmid | Genetic features | Origin |
|---|---|---|---|
| INVSc1 | - | *MATa his3D1 leu2 trp1-289 ura3-52 MAT his3D1 leu2 trp1-289 ura3-52,* diploid strain. | Thermo Fisher Scientific |
| EV | pFV334 | INVSc1 strain transformed with pFV334 plasmid. | This work |
| *At*CER1-1 | pFV570 | INVSc1 strain transformed with pFV570 plasmid. | This work |
| *At*CER3 | pFV571 | INVSc1 strain transformed with pFV571 plasmid. | This work |
| *At*CER1-1 + *At*CER3 | pFV573 | INVSc1 strain transformed with pFV573 plasmid. | This work |
| *At*CER1-1-H159A + *At*CER3 | pFV575 | INVSc1 strain transformed with pFV575 plasmid. | This work |
| *At*CER1-1 + *At*CER3-C587A | pFV896 | INVSc1 strain transformed with pFV896 plasmid. | This work |
| *At*CER1-1 + *At*CER3--A461N T462L | pFV897 | INVSc1 strain transformed with pFV897 plasmid. | This work |
| *No*CER1-1 + *No*CER3-1 | pFV796 | INVSc1 strain transformed with pFV796 plasmid | This work |
| *Ot*CER1/3 | pFV576 | INVSc1 strain transformed with pFV576 plasmid | This work |
| *Kn*CER1/3 | pFV818 | INVSc1 strain transformed with pFV818 plasmid | This work |
| *Pc*CER1/3 | pFV817 | INVSc1 strain transformed with pFV817 plasmid | This work |
| *Mk*CER1/3 | pFV819 | INVSc1 strain transformed with pFV819 plasmid | This work |
| *Ps*CER1/3 | pFV816 | INVSc1 strain transformed with pFV816 plasmid | This work |
| *Ot*CER1/3 + *At*CER1-1 | pFV579 | INVSc1 strain transformed with pFV579 plasmid | This work |
| *Ot*CER1/3 + *At*CER3 | pFV582 | INVSc1 strain transformed with pFV582 plasmid | This work |
| *Kn*CER1/3 + *At*CER3 | pFV832 | INVSc1 strain transformed with pFV832 plasmid | This work |
| *Pc*CER1/3 + *At*CER3 | pFV831 | INVSc1 strain transformed with pFV831 plasmid | This work |
| *Mk*CER1/3 + *At*CER3 | pFV833 | INVSc1 strain transformed with pFV833 plasmid | This work |
| *Ps*CER1/3 + *At*CER3 | pFV830 | INVSc1 strain transformed with pFV830 plasmid | This work |
| *Ot*CER1/3-H142A | pFV701 | INVSc1 strain transformed with pFV701 plasmid | This work |
| *Ot*CER1/3-H155A | pFV702 | INVSc1 strain transformed with pFV702 plasmid | This work |
| *Ot*CER1/3-H243A | pFV703 | INVSc1 strain transformed with pFV703 plasmid | This work |
| *Ot*CER1/3-A455N -T456L | pFV813 | INVSc1 strain transformed with pFV813 plasmid | This work |
| *Ot*CER1/3+*At*CER3-H147A | pFV704 | INVSc1 strain transformed with pFV704 plasmid | This work |
| *Ot*CER1/3+*At*CER3-H161A | pFV705 | INVSc1 strain transformed with pFV705 plasmid | This work |
| *Ot*CER1/3+*At*CER3-H250A | pFV706 | INVSc1 strain transformed with pFV706 plasmid | This work |
| *Ot*CER1/3+*At*CER3-C587A | pFV892 | INVSc1 strain transformed with pFV892 plasmid | This work |
| OtCER1/3+AtCER3-A461N-T462L | pFV893 | INVSc1 strain transformed with pFV892 plasmid | This work |

**Table S4: List of genomes from different species or strains of green algae investigated for the presence of FAP and CER1/CER3 homologs.** The genomes of this list were obtained from the PhycoCosm database (https://phycocosm.jgi.doe.gov/algae/algae.info.html).

| Genomes investigated to look for presence of FAP and CER1/CER3 homologs | Classification |
|---|---|
| *Prasinoderma coloniale CCMP1413* | Prasinodermophytes |
| *Mesostigma viride CCAC 1140* | Streptophytes Mesostigmatophyceae |
| *Mesostigma viride NIES-296* | Streptophytes Mesostigmatophyceae |
| *Chlorokybus atmophyticus CCAC 0220* | Streptophytes Chlorokybophyceae |
| *Klebsormidium nitens* | Streptophytes Klebsormidiophyceae |
| *Chara braunii S276* | Streptophytes Charophyceae |
| *Spirogloea muscicola CCAC 0214* | Streptophytes Zygnematophyceae |
| *Zygnema circumcarinatum UTEX 1559* | Streptophytes Zygnematophyceae |
| *Zygnema cf. cylindricum SAG 698-1a* | Streptophytes Zygnematophyceae |
| *Zygnema circumcarinatum SAG 698-1b* | Streptophytes Zygnematophyceae |
| *Zygnema circumcarinatum UTEX 1560* | Streptophytes Zygnematophyceae |
| *Mesotaenium endlicherianum SAG12.97* | Streptophytes Zygnematophyceae |
| *Mesotaenium kramstae Lemmermann NIES-658 v3.0* | Streptophytes Zygnematophyceae |
| *Mesotaenium kramstae Lemmermann NIES-657 v3.0* | Streptophytes Zygnematophyceae |
| *Bathycoccus prasinos RCC1105* | Chlorophytes Mamiellophyceae |
| *Crustomastix stigmatica CCMP3273* | Chlorophytes Mamiellophyceae |
| *Micromonas commoda NOUM17 RCC299 v3.0* | Chlorophytes Mamiellophyceae |
| *Micromonas pusillaCCMP1545 v3.0* | Chlorophytes Mamiellophyceae |
| *Ostreococcus lucimarinus CCE9901 v2.0* | Chlorophytes Mamiellophyceae |
| *Ostreococcus 'lucimarinus' clade-A-BCC118000* | Chlorophytes Mamiellophyceae |
| *Ostreococcus mediterraneus clade-D-RCC2572* | Chlorophytes Mamiellophyceae |
| *Ostreococcus mediterraneus clade-D-RCC1621* | Chlorophytes Mamiellophyceae |
| *Ostreococcus mediterraneus clade-D-RCC1107* | Chlorophytes Mamiellophyceae |
| *Ostreococcus tauri RCC1115 v1.0* | Chlorophytes Mamiellophyceae |
| *Ostreococcus tauri RCC4221 v3.0* | Chlorophytes Mamiellophyceae |
| *Ostreococcus sp. RCC809* | Chlorophytes Mamiellophyceae |
| *Asterochloris glomerata Cgr/DA1pho v2.0* | Chlorophytes Trebouxiophyceae |
| *Auxenochlorella protothecoides 0710* | Chlorophytes Trebouxiophyceae |
| *Botryococcus braunii Showa* | Chlorophytes Trebouxiophyceae |
| *Chlorella variabilis NC64A v1.0* | Chlorophytes Trebouxiophyceae |
| *Chlorellaceae sp. bin 3300059473_4402 v1.0* | Chlorophytes Trebouxiophyceae |
| *Chlorellaceae sp. bin 3300059473_4966 v1.0* | Chlorophytes Trebouxiophyceae |
| *Coccomyxa subellipsoidea C-169 v3.0* | Chlorophytes Trebouxiophyceae |
| *Helicosporidium sp. ATCC 50920* | Chlorophytes Trebouxiophyceae |
| *Nannochloris desiccata UTEX 2437* | Chlorophytes Trebouxiophyceae |
| *Nannochloris desiccata UTEX 2526* | Chlorophytes Trebouxiophyceae |
| *Picochlorum soloecismus DOE101* | Chlorophytes Trebouxiophyceae |
| *Symbiochloris reticulata Africa extracted metagenome v1.0* | Chlorophytes Trebouxiophyceae |
| *Symbiochloris reticulata Scotland extracted metagenome v1.0* | Chlorophytes Trebouxiophyceae |
| *Symbiochloris reticulata Spain extracted metagenome v1.0* | Chlorophytes Trebouxiophyceae |
| *Symbiochloris reticulata Spain reference genome v1.0* | Chlorophytes Trebouxiophyceae |

| | |
|---|---|
| *Symbiochloris reticulata Switzerland extracted metagenome v1.0* | Chlorophytes Trebouxiophyceae |
| *Trebouxiophyceae sp. bin 3300059473_6682 v1.0* | Chlorophytes Trebouxiophyceae |
| *Trebouxiophyceae sp. bin 3300059473_978 v1.0* | Chlorophytes Trebouxiophyceae |
| *Tetraselmis striata* | Chlorophytes Chlorodendrophyceae |
| *Tetraselmis GSL018* | Chlorophytes Chlorodendrophyceae |
| *Tetraselmis astigmatica CCMP880* | Chlorophytes Chlorodendrophyceae |
| *Chloropicon primus CCMP1205* | Chlorophytes Chloropicophyceae |
| *Picocystis sp. ML* | Chlorophytes Picocystophyceae |
| *Astrephomene gubernaculifera NIES-4017* | Chlorophytes Chlorophyceae |
| *Chlamydomonas priscuii UWO241* | Chlorophytes Chlorophyceae |
| *Chlamydomonas reinhardtii CC-4532 v6.1* | Chlorophytes Chlorophyceae |
| *Chlamydomonas reinhardtii CC-503 v5.6* | Chlorophytes Chlorophyceae |
| *Chloromonas remiasii CCCryo 005-99 v1.0* | Chlorophytes Chlorophyceae |
| *Chromochloris zofingiensis SAG 211-14 v5.0* | Chlorophytes Chlorophyceae |
| *Desmodesmus armatus UTEX B 2533 v2.0* | Chlorophytes Chlorophyceae |
| *Dunaliella salina CCAP19/18* | Chlorophytes Chlorophyceae |
| *Enallax costatus CCAP 276/31 v1.1* | Chlorophytes Chlorophyceae |
| *Flechtneria rotunda SEV3VF49 v1.1* | Chlorophytes Chlorophyceae |
| *Gonium pectorale NIES-2863* | Chlorophytes Chlorophyceae |
| *Monoraphidium minutum 26B-AM v1.0* | Chlorophytes Chlorophyceae |
| *Monoraphidium neglectum SAG 48.87* | Chlorophytes Chlorophyceae |
| *Raphidocelis subcapitata NIES-35* | Chlorophytes Chlorophyceae |
| *Raphidocelis subcapitata NIES-35* | Chlorophytes Chlorophyceae |
| *Scenedesmus obliquus EN0004 v1.0* | Chlorophytes Chlorophyceae |
| *Scenedesmus obliquus UTEX 3031* | Chlorophytes Chlorophyceae |
| *Scenedesmus obliquus UTEX 393* | Chlorophytes Chlorophyceae |
| *Scenedesmus obliquus UTEX 393 v2.0* | Chlorophytes Chlorophyceae |
| *Scenedesmus obliquus var. DOE0013 v1.0* | Chlorophytes Chlorophyceae |
| *Scenedesmus obliquus var. UTEX 1450 v1.0* | Chlorophytes Chlorophyceae |
| *Scenedesmus obliquus var. UTEX2630 v1.0* | Chlorophytes Chlorophyceae |
| *Scenedesmus sp. NREL 46B-D3 v1.0* | Chlorophytes Chlorophyceae |
| *Tetradesmus deserticola SNI-2 v1.1* | Chlorophytes Chlorophyceae |
| *Tetradesmus obliquus UTEX 72 v1.1* | Chlorophytes Chlorophyceae |
| *Volvox carteri v2.1* | Chlorophytes Chlorophyceae |
| *Ulva mutabilis Foyn* | Chlorophytes Ulvophyceae |
| *Caulerpa lentillifera* | Chlorophytes Ulvophyceae |
| *Ostreobium quekettii SAG 6.99* | Chlorophytes Ulvophyceae |

**Table S5: Table summarizing several algal CER1/3 proteins analyzed in this study and their identity towards *A. thaliana* CER1-1 (*At*CER1-1) and CER3 (*At*CER3) proteins.**

| Algal CER1/3 | Specie | Subkingdom | Phylum | % identity with *At*CER1-1 | % identity with *At*CER3 |
|---|---|---|---|---|---|
| *Ot*CER1/3 | *Ostreococcus tauri (RCC4221)* | *Viriplantae* | *Chlorophyta* | 40.1% | 46.9% |
| *Mc*CER1/3 | *Micromonas comoda (RCC299)* | *Viriplantae* | *Chlorophyta* | 40.7% | 46.4% |
| *Tc*CER1/3 | *Tetraselmis convolutae* | *Viriplantae* | *Chlorophyta* | 39.0% | 44.8% |
| *Pc*CER1/3 | *Prasinoderma coloniale (CCMP1413)* | *Viriplantae* | *Prasinodermophyta* | 38.1% | 45.2% |
| *Kn*CER1/3 | *Klebsormidium nitens (NIES-2285)* | *Viriplantae* | *Streptophyta* | 39.9% | 44.0% |
| *Mk*CER1/3 | *Mesotaenium kramstae (Lemmermann NIES-657)* | *Viriplantae* | *Streptophyta* | 43.5% | 50.2% |
| *Ps*CER1/3 | *Proteomonas sulcata (CCMP1175)* | *Cryptophyta* | *Cryptista* | 38.3% | 43.0% |
| *Cc*CER1/3 | *Cryptophyceae sp. CCMP2293* | *Cryptophyta* | *Cryptista* | 37.3% | 42.6% |
